# Large-Scale Deep Multi-Layer Analysis of Alzheimer’s Disease Brain Reveals Strong Proteomic Disease-Related Changes Not Observed at the RNA Level

**DOI:** 10.1101/2021.04.05.438450

**Authors:** Erik C.B. Johnson, E. Kathleen Carter, Eric B. Dammer, Duc M. Duong, Ekaterina S. Gerasimov, Yue Liu, Jiaqi Liu, Ranjita Betarbet, Lingyan Ping, Luming Yin, Geidy E. Serrano, Thomas G. Beach, Junmin Peng, Philip L. De Jager, Vahram Haroutunian, Bin Zhang, Chris Gaiteri, David A. Bennett, Marla Gearing, Thomas S. Wingo, Aliza P. Wingo, James J. Lah, Allan I. Levey, Nicholas T. Seyfried

**Affiliations:** Goizueta Alzheimer’s Disease Research Center, Emory University School of Medicine, Atlanta, GA, USA; Department of Neurology, Emory University School of Medicine, Atlanta, GA, USA; Department of Biochemistry, Emory University School of Medicine, Atlanta, GA, USA; Department of Genetics, Emory University School of Medicine, Atlanta, GA, USA; Department of Psychiatry, Emory University School of Medicine, Atlanta, GA, USA; Department of Pathology and Laboratory Medicine, Emory University School of Medicine, Atlanta, GA, USA; Banner Sun Health Research Institute, Sun City, AZ, USA; Departments of Structural Biology and Developmental Neurobiology, St. Jude Children’s Research Hospital, Memphis, TN, USA; Center for Proteomics and Metabolomics, St. Jude Children’s Research Hospital, Memphis, TN, USA; Center for Translational & Computational Neuroimmunology, Department of Neurology, Taub Institute, Columbia University Irving Medical Center, New York Presbyterian Hospital, New York, NY, USA; Departments of Psychiatry and Neuroscience, Icahn School of Medicine at Mount Sinai, One Gustave L. Levy Place, New York, NY, USA; JJ Peters VA Medical Center MIRECC, 130 West Kingsbridge Road, Bronx, NY, USA; Department of Genetics and Genomic Sciences, Mount Sinai Center for Transformative Disease Modeling, Icahn School of Medicine at Mount Sinai, One Gustave L. Levy Place, New York, NY, USA; Rush Alzheimer’s Disease Center, Rush University Medical Center, Chicago, IL, USA; Division of Mental Health, Atlanta VA Medical Center, GA, USA

## Abstract

The biological processes that are disrupted in the Alzheimer’s disease (AD) brain remain incompletely understood. We recently performed a proteomic analysis of >2000 brains to better understand these changes, which highlighted alterations in astrocytes and microglia as likely key drivers of disease. Here, we extend this analysis by analyzing >1000 brain tissues using a tandem mass tag mass spectrometry (TMT-MS) pipeline, which allowed us to nearly triple the number of quantified proteins across cases. A consensus protein co-expression network analysis of this deeper dataset revealed new co-expression modules that were highly preserved across cohorts and brain regions, and strongly altered in AD. Nearly half of the protein co-expression modules, including modules significantly altered in AD, were not observed in RNA networks from the same cohorts and brain regions, highlighting the proteopathic nature of AD. Two such AD-associated modules unique to the proteomic network included a module related to MAPK signaling and metabolism, and a module related to the matrisome. Analysis of paired genomic and proteomic data within subjects showed that expression level of the matrisome module was influenced by the *APOE ε4* allele, but was not related to the rate of cognitive decline after adjustment for neuropathology. In contrast, the MAPK/metabolism module was strongly associated with the rate of cognitive decline. Disease-associated modules unique to the proteome are sources of promising therapeutic targets and biomarkers for AD.

## Introduction

Alzheimer’s disease (AD) remains an immense personal and public health burden without effective disease-modifying therapies. As part of a national effort to develop therapeutics and biomarkers for AD, the Accelerated Medicines Partnership—Alzheimer’s Disease (AMP-AD) Consortium has been leveraging unbiased molecular profiling data at the genomic, transcriptomic, proteomic, and metabolomic levels to further our understanding of AD pathogenesis. Genetics has significantly advanced our understanding of AD heritable risk, yet how genetic risk factors affect biological pathways that influence AD pathophysiology is not always clear^1^. Understanding the biological effects of AD risk factor polymorphisms in human brain often requires additional levels of analysis using other –omics approaches. To this end, transcriptomics has been widely used to measure mRNA transcripts in AD brain, and the resulting transcriptomic data have been integrated with AD genetic risk^2^. However, the ultimate biological effectors of AD genetic and environmental risk are often the proteins and the metabolic pathways they modulate. Compared to genomics and transcriptomics, proteomics approaches have to date provided comparatively less in-depth coverage of the target analyte due to increased complexity and technical demands of analyzing amino acid polymers *versus* nucleic acid polymers.

As the proteomics team of AMP-AD, we have been working to better characterize AD at the proteomic level, and relate observed proteomic changes to other levels of –omics data in order to advance our understanding of the disease. As part of our initial efforts towards this goal, we recently reported a large multi-center study using an unbiased discovery approach to better understand the proteomic changes that occur in AD brain in both early and late stages of the disease^3^. We used protein co-expression network analysis to cluster proteins into groups that reflect distinct biological functions, processes, pathways, and cell types, and related these protein groups—or modules—to early and late AD stages. This protein co-expression network of 13 modules revealed that the module most strongly correlated to AD was related to astrocyte and microglia metabolism. Furthermore, proteins from this module could be detected in cerebrospinal fluid, suggesting that they may be useful biomarkers for tracking this AD brain pathophysiology.

Our consensus AD protein network was constructed on 3334 proteins measured across >450 brain tissues by label-free quantitation mass spectrometry (LFQ-MS), which was an approach we have previously used to generate other protein co-expression networks of smaller scale^4, 5^. LFQ-MS is typically performed using a “single-shot” approach where a single fractionation of the sample is performed prior to MS analysis and protein identification and quantification. LFQ-MS suffers from missing protein measurements that accumulate across sample sets and that ultimately reduce the final number of reliably quantified proteins^6^. This problem becomes particularly acute when analyzing large cohorts, and limits the depth of proteomic analysis. More recently, a different quantitative mass spectrometry approach based on peptide labeling and quantification using tandem mass tags (TMT-MS) has been developed that helps to address, in part, some of the limitations inherent in LFQ-MS^7–9^. The TMT-MS approach allows for orthogonal pre-fractionation of samples prior to a second fractionation step and analysis on the mass spectrometer which, along with batching of samples to partially mitigate missing measurements, allows for deeper proteome coverage than what can be achieved through LFQ-MS^10^.

In this study, we used the TMT-MS approach and the AMP-AD consortium of postmortem brain tissues to generate a deeper TMT AD protein network that significantly expanded our previous LFQ network and revealed new AD-related protein co-expression modules. We leveraged brain tissues from cohorts that also have been profiled using other –omics modalities, including genomics and transcriptomics. Multi-layer –omics data provide the opportunity to relate and integrate different levels of biological information into a holistic understanding of disease pathophysiology, and identify key molecular drivers of disease based on observations across different levels of data. For instance, the relationship of RNA to protein levels, as well as the genetic variants that affect these levels, are important considerations when identifying causal variants and prioritizing drug discovery efforts^11^. To this end, we undertook a multi-layer genomic, transcriptomic, and proteomic analysis of the TMT AD protein network to better understand the relationships among these different data types. We found that some of the most strongly AD-related modules in the proteomic network had no cognate modules in RNA networks. Many of these modules also correlated with cognitive trajectory during life, and one of them—the matrisome module—was influenced by genetic variation in apolipoprotein E (*APOE*), a known AD genetic risk factor. Our findings highlight the importance of including protein measurements when studying AD pathophysiology and selecting potential targets for disease-modifying therapeutic development.

## Results

### Construction of a TMT Consensus AD Protein Co-expression Network

For this study, we analyzed a total of 516 dorsolateral prefrontal cortex (DLPFC) tissues from control, asymptomatic AD (AsymAD), and AD brains from the Religious Orders Study and Memory and Aging Project (ROSMAP, *n*=84 control, 148 AsymAD, 108 AD)^12–14^ and Banner Sun Health Research Institute (Banner, *n*=26 control, 58 AsymAD, 92 AD)^15^ by tandem mass tag mass spectrometry (TMT-MS)-based quantitative proteomics (**Figure 1A, Supplementary Table 1**). Cases were defined based on a unified classification scheme using semi-quantitative histopathological measures of Aβ and tau neurofibrillary tangle deposition^16–19^, as well as cognitive function near time of death, as previously described^3^. AsymAD cases were those with neuropathological burden of Aβ plaques and tau tangles similar to AD cases, but without significant cognitive impairment near time of death, which is considered to be an early preclinical stage of AD^20^. After data processing and outlier removal, a total of 8619 proteins were used to build a protein co-expression network using the weighted co-expression network (WGCNA) algorithm^21^ (**Figure 1B, Supplementary Figure 1, Supplementary Tables 2-4**). This network consisted of 44 modules, or communities of proteins related to one another by their co-expression across control and disease tissues. All modules contained proteins that were quantified using multiple peptides per protein (**Supplementary Table 3**). Compared to our previous AD consensus network constructed using label-free quantitative (LFQ) proteomic data, the TMT consensus network contained over five times as many proteins that could be assigned to a module (6337 *versus* 1205), as well as a larger fraction of quantified proteins that could be assigned to a module (73% *versus* 36%), highlighting the improved depth and coherence of the TMT data compared to the LFQ consensus data. Of the 13 modules previously identified in the LFQ consensus network^3^, every module except the smallest module (module 13 consisting of 20 proteins) was preserved in the TMT network (**Supplementary Figure 2A**), also highlighting the consistency of the LFQ and TMT proteomic data. Because different network clustering algorithms can produce disparate networks, we tested the robustness of the TMT consensus network generated by the WGCNA algorithm by also generating a co-expression network using an independent algorithm—the MONET M1 algorithm. MONET M1 was identified as one of the top performers in the Disease Module Identification DREAM challenge, and is based on a modularity optimization algorithm rather than the hierarchical clustering approach used in WGCNA^22, 23^. We found that all 44 WGCNA modules were highly preserved in the MONET M1 network (**Supplementary Figure 2B**), demonstrating the robustness of the TMT consensus network to clustering algorithm.

**Figure 1.**
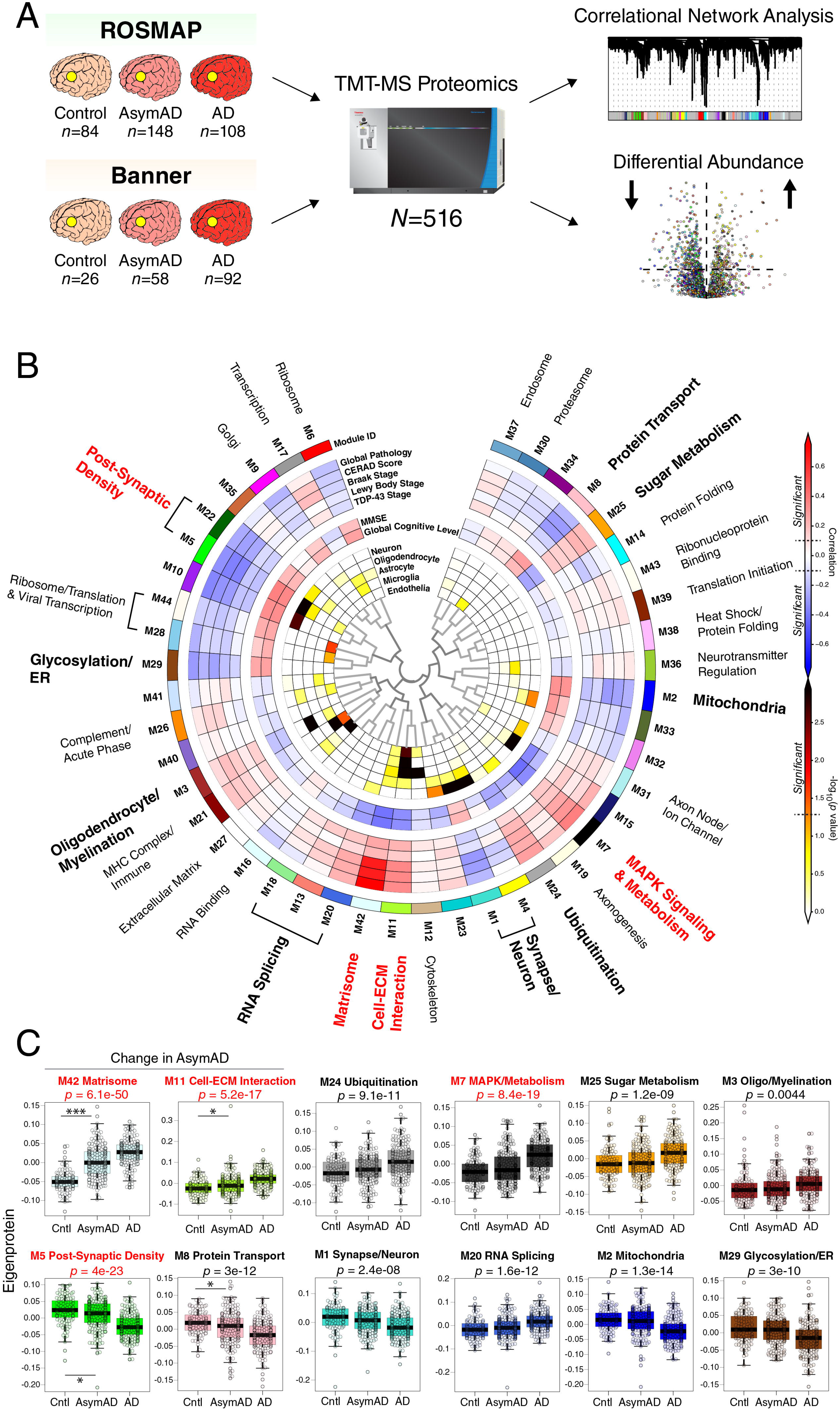
TMT AD Protein Co-Expression Network. (A-C) 516 dorsolateral prefrontal cortex (DLPFC) tissues from the Religious Orders Study and Memory and Aging Project (ROSMAP, *n*=84 control, 148 AsymAD, 108 AD) and Banner Sun Health Brain Bank (Banner, *n*=26 control, 58 AsymAD, 92 AD) were analyzed by tandem mass tag mass spectrometry (TMT-MS)-based proteomics (A). After outlier removal and data processing, a total of 8619 proteins were quantified across 488 cases, which were analyzed by both differential expression and co-expression approaches. (B) A protein co-expression network was built using the weighted co-expression network algorithm (WGCNA), which consisted of 44 protein co-expression modules. Module relatedness is shown in the central dendrogram. Gene ontology analysis was used to identify the principal biology represented by each module. Modules that did not have a clear ontology were not assigned an ontology term. Module eigenproteins were correlated with neuropathological and cognitive traits present in the ROSMAP and Banner cohorts (red, positive correlation; blue, negative correlation). The global pathology, Lewy body stage, TDP-43 stage, and global cognitive level traits were present only in ROSMAP. Twelve of the 44 modules that were most highly correlated to neuropathological and/or cognitive traits are in bold, with the four most strongly trait-related modules highlighted in red. The cell type nature of each module was assessed by module protein overlap with cell type specific marker lists of neurons, oligodendrocytes, astrocytes, microglia, and endothelia. (C) Module eigenprotein levels by case status for the twelve most strongly trait-correlated modules bolded in (B). Modules are grouped by those that change in AsymAD (*n*=4, left), and those that change only in AD (*n*=8, right). *n*=106 control, 200 AsymAD, and 182 AD. Correlations were performed using biweight midcorrelation and corrected by the Benjamini-Hochberg procedure. Cell type marker overlap was assessed using Fisher’s exact test (FET) and corrected using the Benjamini-Hochberg procedure. Differences in module eigenprotein by case status were assessed by one-way ANOVA with Tukey test. \**p*<0.05, \*\*\**p*<0.001. Boxplots represent the median, 25^th^, and 75^th^ percentiles, and box hinges represent the interquartile range of the two middle quartiles within a group. Datapoints up to 1.5 times the interquartile range from box hinge define the extent of whiskers (error bars). AD, Alzheimer’s disease; AsymAD, asymptomatic Alzheimer’s disease; Cntl, control; ECM, extracellular matrix; ER, endoplasmic reticulum; MAPK, mitogen-activated protein kinase; MHC, major histocompatibility complex.

The biology represented by each TMT consensus network module was determined using gene ontology (GO) analysis of its constituent proteins (**Figure 1B, Extended Data 1**). Most modules could be assigned a primary ontology, and those that were ambiguous in their ontology were left unannotated or assigned as “ambiguous.” Module 42 was assigned the term “matrisome,” which refers to the collection of extracellular matrix-associated proteins^24, 25^, due to its strong enrichment in extracellular matrix and glycosaminoglycan binding proteins. To assess whether a module was related to features of AD, we correlated each module eigenprotein, or the first principal component of module protein expression, to neuropathologic or cognitive traits present in the ROSMAP and Banner cohorts (**Figure 1B, Supplementary Table 5; individual protein trait correlations are provided in Supplementary Table 6**). We also assessed the cell type nature of each module by determining whether it was enriched in cell type-specific protein markers (**Figure 1B, Supplementary Tables 7 and 8**). Because the network was highly powered, with the ability to observe a significant correlation of 0.1 at *p*=0.05 for most pathological and cognitive traits, we were able to observe a large fraction of the 44 modules that significantly correlated with at least one pathological or cognitive trait. Twelve modules or module families were noted to correlate more strongly to AD traits than the others. These included post-synaptic density, glycosylation/endoplasmic reticulum (ER), oligodendrocyte/myelination, RNA splicing, matrisome, cell-extracellular matrix (ECM) interaction, synapse/neuron, ubiquitination, MAPK signaling and metabolism, mitochondria, sugar metabolism, and protein transport modules (**Figure 1B**). Four of these modules—M5 post-synaptic density (global pathology *r*=–0.32, *p*=2.7e^-9^; global cognitive function *r*=0.35, *p*=7.4e^-11^), M7 MAPK signaling and metabolism (global pathology *r*=0.37, *p*=4.9e^-12^; global cognitive function *r*=–0.42, *p*=1.2e^-15^), M11 cell-ECM interaction (global pathology *r*=0.34, *p*=4.1e^-10^; global cognitive function *r*=–0.33, *p*=1.1e^-9^), and M42 matrisome (global pathology *r*=0.75, *p*=1.1e^-60^; global cognitive function *r*=–0.4, *p*=2.3e^-14^)— were the most strongly correlated to AD neuropathology or cognition out of the twelve. The increased depth of the TMT network allowed for additional resolution of AD correlated modules identified in the prior LFQ consensus network. For instance, the M4 astrocyte/microglia metabolism module that was most highly correlated to AD in the LFQ network was split into two predominant modules in the TMT network: the M7 MAPK/metabolism module and the M11 cell-ECM interaction module (**Supplementary Figure 2C**), both of which were correlated to AD. Similarly, the large M1 synapse/neuron module in the LFQ network was resolved into three large neuronal/synaptic modules (M1, M4, and M5), one of which, M5, reflected post-synaptic biology and was strongly correlated to AD pathology and cognitive impairment. Sugar metabolism, which was the primary ontology for M4 in the LFQ network, was split into the M7 MAPK/metabolism and M25 sugar metabolism modules in the TMT network, both of which were significantly correlated to AD but did not have strong cell type character as suggested by co-expression analysis in the LFQ network. Two modules in the LFQ network that had ambiguous ontologies, M12 and M13, could now be assigned with additional depth in the TMT network as reflecting protein transport (M8) and protein folding (M14) functions. The additional analytical depth afforded by the TMT pipeline also allowed us to identify a significant number of new modules that had little to no overlap with the LFQ network. Among these were the M17 transcription, M21 MHC complex/immune, M6 ribosome, M19 axonogenesis, and M9 Golgi modules, in which approximately 80% or more of the module proteins were not quantified in the LFQ network, including a majority of proteins with strong correlation to the module eigenprotein (i.e., “hub” proteins) (**Supplementary Figure 2D**). Three of these new modules—the M24 ubiquitination, M29 glycosylation/ER, and M42 matrisome modules—were strongly correlated to AD endophenotypes.

To assess whether a given TMT network module was altered in the early stages of AD, we compared the module eigenprotein across control, AsymAD, and AD cases (**Figure 1C, Supplementary Table 9, Extended Data 2**). Four of the twelve most highly AD correlated modules were either significantly increased or decreased in AsymAD compared to control, whereas the other eight were largely altered in AD only. Modules that were increased in AsymAD included M42 matrisome and M11 cell-ECM interaction, whereas modules that were decreased in AsymAD included M5 post-synaptic density and M8 protein transport. Modules that did not significantly change in AsymAD included M3 oligo/myelination, M20 RNA splicing, and M2 mitochondria, among others, consistent with findings in the LFQ network^3^.

The well-known heterogeneity in levels of tau protein domains in AD complicated the measurement and co-expression patterns of the full-length tau isoform in our proteomic dataset^5, 26, 27^. To address this issue, we performed an additional database search in which the tau microtubule binding domain (MTBR) and all other non-MTBR peptides (dMTBR) were entered as separate protein entries for quantification, and correlated each module to MTBR, dMTBR, the MTBR/dMTBR ratio^5^, as well as to all 41 tau peptides measured in our dataset (**Supplementary Tables 10-12**). Modules that correlated most strongly with elevated MTBR protein and peptide levels were the M42 matrisome and M11 cell-ECM modules, whereas modules that correlated most strongly in a negative direction with MTBR were M5 post-synpatic density, and M1 and M4 synapse/neuron modules. Modules that correlated with MTBR tended to be also altered in AsymAD.

In summary, TMT-MS analysis of >500 brain tissues allowed us to quantify >8600 proteins and construct a robust protein co-expression network that was highly powered to detect AD-correlated modules, including a significant number of new modules not present in the previous LFQ consensus network. Some of these new modules were also altered in early stages of AD, likely reflecting pathophysiologic processes that develop in the presence of AD neuropathology but prior to cognitive decline, and correlated with tau dyshomeostasis.

### TMT AD Network Modules Are Preserved Across Different Cohorts and Brain Regions

The TMT AD protein network was generated from DLPFC Brodmann area 9 (BA9) tissues from two centers analyzed at one institution. To determine whether the network modules were also present in other brain regions and robust to center and analytical pipeline, we analyzed 226 paired tissues from frontal cortex BA6 (*n*=113) and temporal cortex BA37 (*n*=113) from 113 subjects in the ROSMAP cohort, 151 parahippocampal gyrus (PHG) BA36 tissues from the Mount Sinai Brain Bank, and 40 tissues from DLPFC BA9 and anterior cingulate BA24 from the Emory Brain Bank, which also included Parkinson’s disease cases (**Figure 2A, Supplementary Tables 13-16**). The Mt. Sinai tissues were analyzed at a different center using a similar mass spectrometry pipeline^28^. All tissues were analyzed using the TMT approach, with the Emory tissues analyzed using the synchronous precursor selection (SPS)-MS3 TMT quantification approach^7, 29^. We generated protein co-expression networks for each cohort, then assessed whether the TMT AD network modules were preserved in each cohort and brain region^3, 30^. We found that nearly all TMT AD network modules were preserved across both cohort and brain region (**Figure 2B, Supplementary Figure 3**).

**Figure 2.**
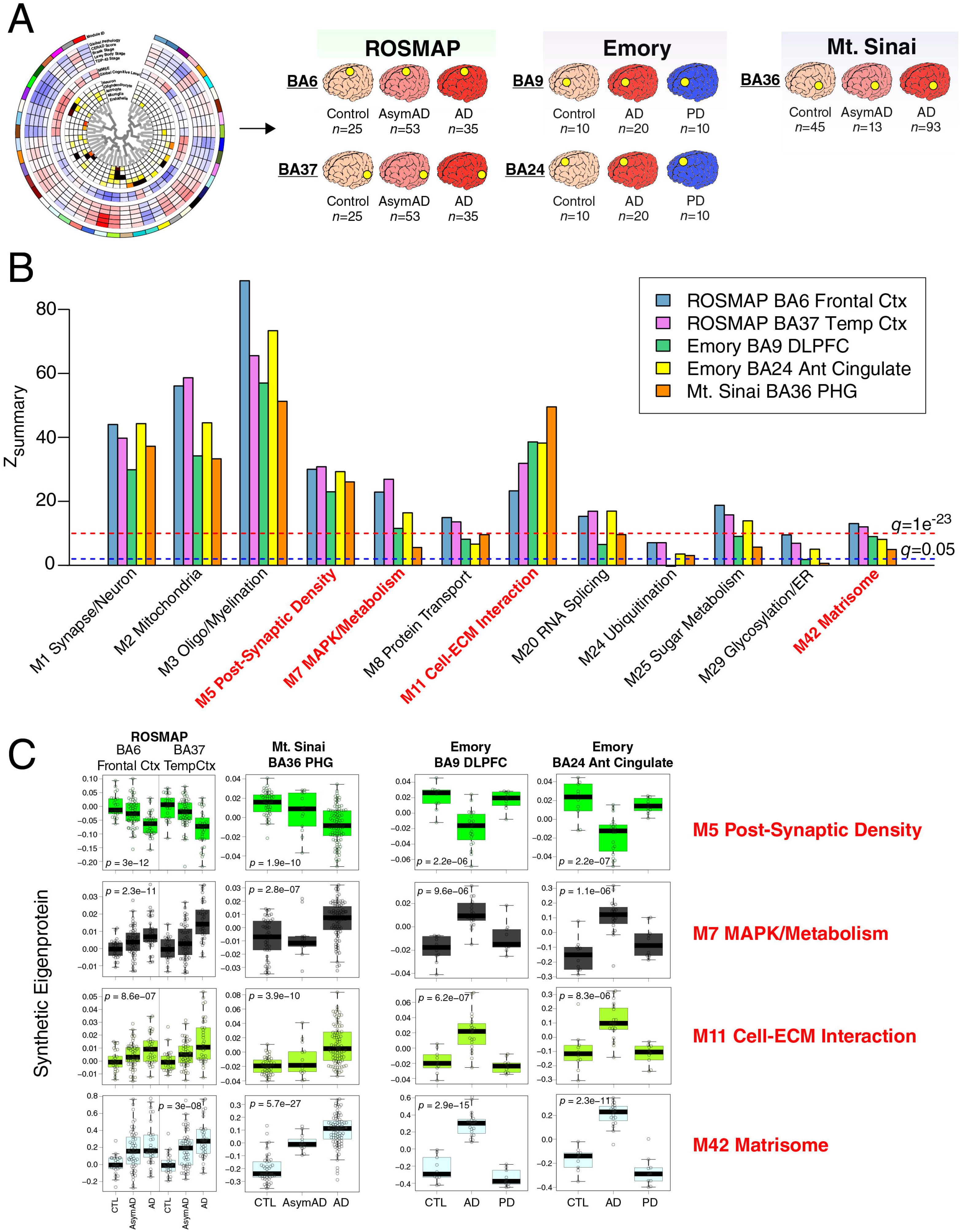
Preservation of the TMT AD Network Across Different Cohorts, Centers, Methods, and Brain Regions. (A-C) Module preservation and synthetic eigenprotein analysis of the TMT AD network generated from dorsolateral prefrontal cortex (DLPFC) Brodmann area 9 (BA9) tissues was performed in ROSMAP Brodmann area 6 (BA6, frontal cortex) and BA37 (temporal cortex), Emory BA9 (DLPFC) and BA24 (anterior cingulate), and Mt. Sinai Brain Bank BA36 (parahippocampal gyrus, PHG) tissues (A). The Emory tissues included Parkinson’s disease cases, and were analyzed using a different TMT quantification approach (synchronous precursor selection (SPS)-MS3). The Mt. Sinai tissues were processed and analyzed by MS2-based TMT-MS at a different center. (B) Module preservation of the twelve trait-correlated modules highlighted in Figure 1B**, C**. Modules that had a *Z*_summary_ score of greater than or equal to 1.96 (or *q*=0.05, blue dotted line) were considered to be preserved, whereas modules that had a *Z*_summary_ score greater than or equal to 10 (or *q*=1e^-23^, red dotted line) were considered to be highly preserved. Preservation statistics for all TMT AD network modules are provided in **Supplementary Figure 3**. (C) Module eigenprotein level by case status was assessed in the different cohorts and brain regions by measuring a TMT AD network synthetic eigenprotein, representing the top 20% of module proteins by module eigenprotein correlation value (kME), in each cohort and region. Synthetic eigenprotein levels are shown for the four most highly trait-correlated TMT AD network modules. Differences and statistics for all modules are provided in **Supplementary Table 17**. Differences in synthetic eigenprotein levels were assessed by one-way ANOVA. Boxplots represent the median, 25^th^, and 75^th^ percentiles, and box hinges represent the interquartile range of the two middle quartiles within a group. Datapoints up to 1.5 times the interquartile range from box hinge define the extent of whiskers (error bars). AD, Alzheimer’s disease; Aβ, amyloid-β; AsymAD, asymptomatic Alzheimer’s disease; ECM, extracellular matrix; ER, endoplasmic reticulum; MAPK, mitogen-activated protein kinase.

We assessed how TMT AD network modules were different by case status in each cohort and brain region by measuring TMT consensus AD network “synthetic” eigenproteins, or the top 20% of proteins within each consensus module, in each separate network (**Figure 2C, Supplementary Table 17**)^3^. Because the ROSMAP BA6 and BA37 tissues were sampled within subject, we were able to compare the module synthetic eigenproteins directly between these two regions within the same individual. All TMT AD network modules were altered in a similar direction to that observed in DLPFC across brain region and cohort. Interestingly, the M7 MAPK/metabolism and M42 matrisome modules were increased more strongly in temporal cortex than frontal cortex when assessed in the same individual, perhaps due to earlier and more severe involvement of this brain region in AD^31^. Most AD associated modules were not significantly altered in PD in either frontal cortex or anterior cingulate, although there appeared to be a trend for the M7 MAPK/metabolism module to increase and the M5 post-synaptic density module to decrease in anterior cingulate, consistent with this brain region being more severely affected in PD compared to DLPFC^32, 33^. In summary, we observed that nearly all TMT AD network modules were preserved across different cohorts, centers, MS methods, and brain regions, demonstrating that the protein co-expression relationships observed are robust to technical artefact and are not unique to the DLPFC.

### TMT AD Protein Network Modules Not Observed in Transcriptomic Networks

Most co-expression network analysis in AD has been performed to date using quantitative RNAseq data. However, not all mRNA transcripts correlate well with protein levels^34, 35^. To compare the similarities and differences between RNA and protein AD co-expression networks, we generated an AD RNA network on 15,582 transcripts measured across 532 ROSMAP DLPFC tissues, 168 of which overlapped with tissues used to generate the TMT AD protein network (**Figure 3A, Supplementary Tables 18-20**). We took care to ensure that the case classification and WGCNA pipeline used for network construction was consistent between protein and RNA datasets. Given the greater number of transcripts measured compared to proteins measured (*n*=15,582 *versus* 8619), the resulting RNA network contained more modules compared to the protein network (n=88 *versus* 44) (**Supplementary Figure 4A, Supplementary Table 20**). We used network preservation statistics to determine which modules in the protein network were preserved in the RNA network^30^. We found that slightly greater than half of the protein modules were preserved in the RNA network (26 out of 44 modules at *Z*_summary_ > 1.96, or *p ≤* 0.05) (**Supplementary Figure 5A**). Among the modules preserved in the RNA network were the AD-associated modules M1 synapse/neuron, M2 mitochondria, M3 oligo/myelination, M5 and M22 post-synaptic density, M8 protein transport, M11 cell-ECM interaction, M20 RNA splicing, and M25 sugar metabolism. However, there were also 18 protein network modules that were not preserved in the RNA network, including the AD-associated modules M7 MAPK/metabolism, M24 ubiquitination, M29 glycosylation/ER, and M42 matrisome (**Figure 3B**). Of these modules, the M7 MAPK/metabolism module—the module most highly correlated to cognitive function in the TMT AD network—was the least preserved, with a Z_summary_ score near 0, indicating its highly unique nature to the proteome. We validated these findings in 193 frontal cortex (BA10) tissues analyzed by RNAseq from the Mt. Sinai Brain Bank, which showed similar network preservation results (**Supplementary Figure 5B**). We also analyzed separately the 168 ROSMAP cases that had paired proteomic and transcriptomic data from the DLPFC region (**Supplementary Figure 5C, Supplementary Tables 18 and 21**). Approximately half (55%) of protein modules from this network were preserved in RNA, indicating that significant differences between protein and RNA co-expression exist even in tissue sampled from the same brain region in the same individual. Co-expression analysis indicated a degree of preservation between protein and RNA networks that was higher than what might be expected based on comparison of differential expression between protein and RNA, which was modest, even in paired tissues (**Supplementary Figure 5D, E**). This correlation remained modest when proteins from the M42 matrisome module, which were the most highly differentially expressed proteins in the TMT AD network (**Supplementary Figure 6**), were excluded from the analysis (**Supplementary Figure 5D, E**). Overall, we found that protein network modules were correlated more strongly to cognitive function than RNA network modules, but that in most cases their correlation to pathology was similar to RNA modules (∼0.15 on average in both positive and negative directions). A striking exception was the M42 matrisome module, which was the module with the strongest correlation to any AD trait, with correlation of 0.75 to global pathology, and which was not present in the RNA network (**Supplementary Figure 4B**). A synthetic eigentranscript of M42 in the RNA network showed minimal relationship to disease, and some eigentranscripts—for instance, M7 MAPK/metabolism—demonstrated an opposite relationship to AD compared to protein, perhaps suggesting the presence of compensatory regulatory mechanisms at the RNA level for these module proteins (**Supplementary Figure 5F**). The protein network also demonstrated an overall larger variance in module AD trait correlations than the RNA network. In summary, we observed that approximately half of the TMT AD protein network modules were not present in RNA networks from the same brain region, including the two protein network modules most strongly correlated to AD pathology and cognitive function, highlighting the unique contribution of the proteome to understanding AD pathophysiology.

**Figure 3.**
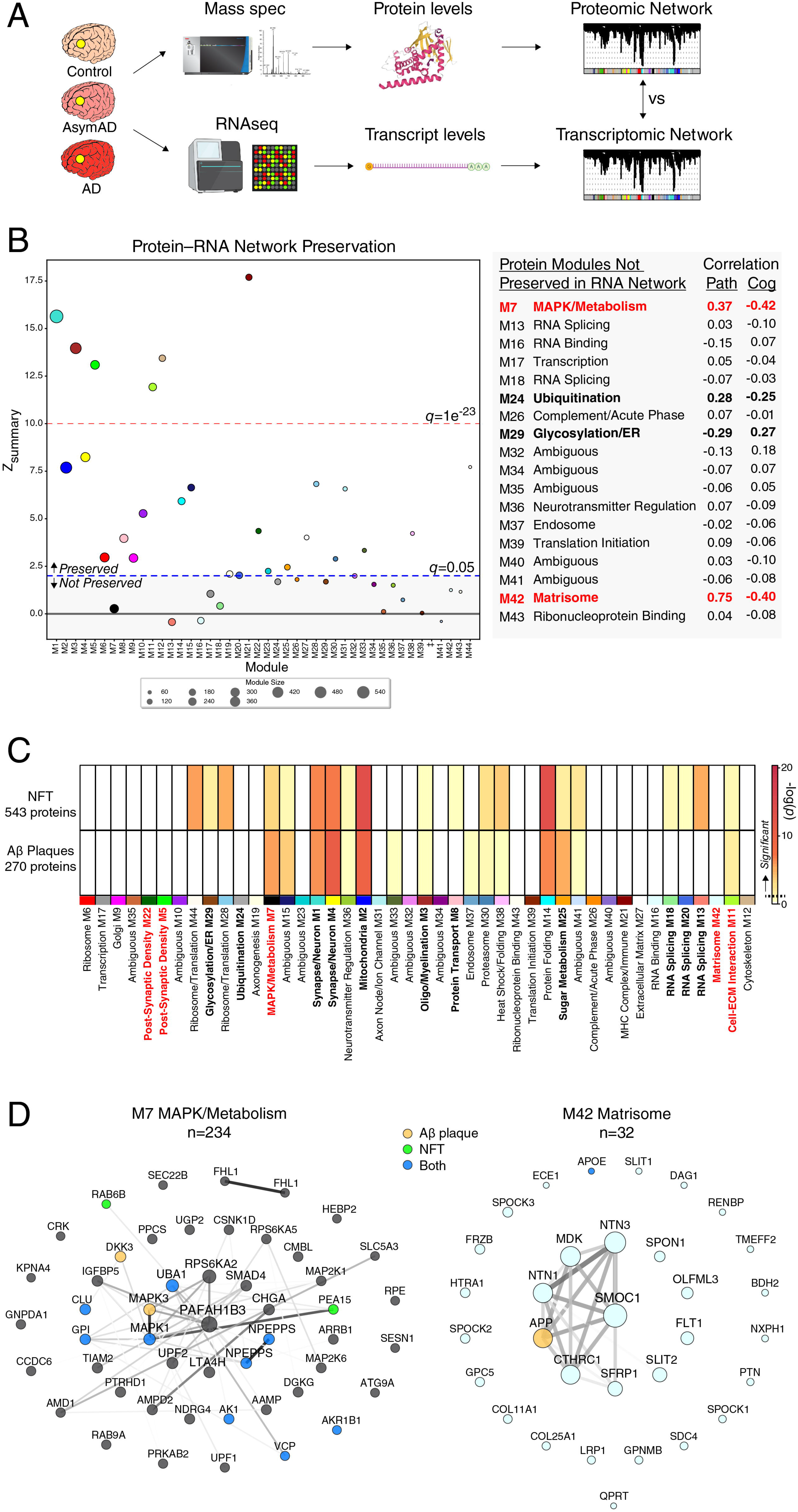
The TMT AD Protein Network Contains Modules Associated with AD That Are Not Present in the Transcriptome. (A-D) Control, AsymAD, and AD frontal cortex tissues from both the ROSMAP cohort (Brodmann area 9; control=125, AsymAD=204, AD=203; 168 overlapping cases with proteomic analysis) and Mt. Sinai Brain Bank (Brodmann area 10; control=54, AsymAD=19, AD=120) were analyzed by RNAseq-based transcriptomics, and co-expression networks generated by WGCNA in similar fashion to the TMT AD protein network (A). (B) Module preservation of the TMT AD protein network into the ROSMAP RNA network. Modules that had a preservation *Z*_summary_ score less than 1.96 (*q*>0.05) were not considered to be preserved. Modules that had a *Z*_summary_ score of greater than or equal to 1.96 (or *q*=0.05, blue dotted line) were considered to be preserved, while modules that had a *Z*_summary_ score greater than or equal to 10 (or *q*=1e^-23^, red dotted line) were considered to be highly preserved. TMT AD network modules that were not preserved in the RNA network, along with their correlation to global pathology and global cognition traits in ROSMAP, are listed on the right. Further information on modules preserved in ROSMAP, as well as preservation analysis with the Mt. Sinai cohort, is provided in **Supplementary Figure 5A, B**. (C) TMT AD network module protein overlap with proteins identified as co-localized with neurofibrillary tangles (NFTs, *n*=543) and amyloid-β (Aβ, *n*=270) plaques as described by Drummond *et al*.^36, 37^. Overlap as shown with a dark yellow hue or darker is considered significant. Overlap with a less stringent set of Aβ plaque associated proteins is provided in **Supplementary Figure 4C**. (D) The top fifty proteins by module eigenprotein correlation value (kME) for the M7 MAPK/metabolism (left, *n*=234 total proteins) and M42 matrisome (right, *n*=32 proteins) modules. Module proteins that were found to be co-localized with NFTs (green), Aβ plaques (orange), or both (blue) are highlighted. Lines between proteins represent correlation matrix adjacency weights. Graphs for all TMT AD network modules are provided in **Extended Data 3**.

### The M7 MAPK/Metabolism Module Is Enriched in Proteins Co-localized with Aβ Plaques and Neurofibrillary Tangles

While co-expression analyses on data obtained from bulk tissue are able to identify modules that are correlated with neuropathological traits such as neurofibrillary tangle (NTF) and Aβ plaque deposition, they do not provide spatial information regarding these relationships. To better understand the potential spatial relationships between the TMT AD protein network modules and the hallmark AD neuropathologies Aβ plaques and NFTs, we performed a module overlap test with proteins that have previously been identified as co-localized with Aβ plaques and NFTs based on laser capture microdissection (LCM) and LFQ proteomic analysis of these structures (**Figure 3C, Supplementary Table 22**)^36, 37^. We found that the M1 and M4 synapse/neuron, M2 mitochondria, and M14 protein folding modules were highly enriched in proteins found in both plaques and tangles. The M7 MAPK/metabolism module was also enriched in both plaques and tangles, but more highly enriched in Aβ plaque-associated proteins. This was also the case for the M25 sugar metabolism module. NFTs were uniquely enriched in proteins from the M28 and M44 ribosome/translation, M29 glycosylation/ER, and M13 RNA splicing modules. Surprisingly, the M42 matrisome module was not significantly enriched with core plaque proteins identified by LCM, even though the amyloid precursor protein (APP) (a proteomic measurement largely driven by Aβ) and apolipoprotein E (ApoE) were members of this module (**Supplementary Table 4**). M42 was highly elevated in AsymAD and AD compared to control, consistent with an association with neuritic plaques. When the analysis was expanded to a less stringent set of plaque-associated proteins identified in at least one LCM experiment rather than proteins identified in multiple experiments, M42 was found to be significantly enriched in plaque-associated proteins (**Supplementary Figure 4C**). This suggested that our TMT proteomic and co-expression analysis was perhaps capturing a significant number of plaque-associated proteins that are less reliably observed by LFQ-MS approaches, even with LCM isolation, such as SPARC-related modular calcium-binding protein 1 (SMOC1), midkine (MDK), and netrin-1 (NTN1). For example, although it was not identified as a core Aβ plaque-associated protein by LCM, MDK demonstrated a pattern of staining on immunohistochemistry consistent with its co-localization with Aβ plaques (**Supplementary Figure 4D**). Other proteins within the M7 MAPK/metabolism and M42 matrisome modules have been shown to co-localize with Aβ plaques and NFTs by immunohistochemistry^28, 38–44^. Many of the proteins within the M42 matrisome module shared heparan sulfate and glycosaminoglycan binding domains likely mediating their interaction with Aβ fibrils^28^ (**Supplementary Figure 4E**). NFT and core Aβ plaque proteins that overlap with the top fifty M7 MAPK/metabolism and M42 matrisome module proteins by module eigenprotein correlation value (kME) are shown in **Figure 3D**. In summary, we found that a number of TMT AD protein network modules were enriched in proteins that are found in NFTs and Aβ plaques, including the M7 MAPK/metabolism module, consistent with a spatial relationship between these biological processes and hallmark AD pathologies.

### M42 Matrisome Module Protein Levels are Influenced by APOE ε4

Because the TMT AD protein network was generated from post-mortem brain tissue, deciphering which biological changes are upstream of the disease process from those that are altered in later stages of disease presents a challenge. One way to separate upstream from downstream changes is to examine the association of genetic variants with the network, on the assumption that genetic modifiers in AD are not influenced by the disease process itself, but rather modify biological processes that are upstream in the disease cascade. To this end, we first assessed which TMT AD network modules were enriched in genetic loci associated with AD as identified by GWAS using a gene-based test of association (**Figure 4A, Supplementary Tables 23-25**)^3, 4, 45^. We found that M42 matrisome, M30 proteasome, and M29 glycosylation/ER modules were significantly enriched for AD risk genes, whereas the M7 MAPK/metabolism module demonstrated a trend towards enrichment. The strong enrichment in M42 was driven by the ApoE protein within the module, given its large effect size on AD risk^1^. When ApoE was removed from the module, M42 was no longer enriched in AD genetic risk loci (data not shown). Interestingly, M42 was also highly enriched for proteins implicated in autism spectrum disorder (ASD), with 28 out of a total of 32 module proteins overlapping with ASD genetic risk factors (**Supplementary Table 25**). In general, AD genetic risk clustered in modules that had glial cell type character or no cell type character, whereas autism and schizophrenia genetic risk clustered in modules with neuronal cell type and synaptic character.

**Figure 4.**
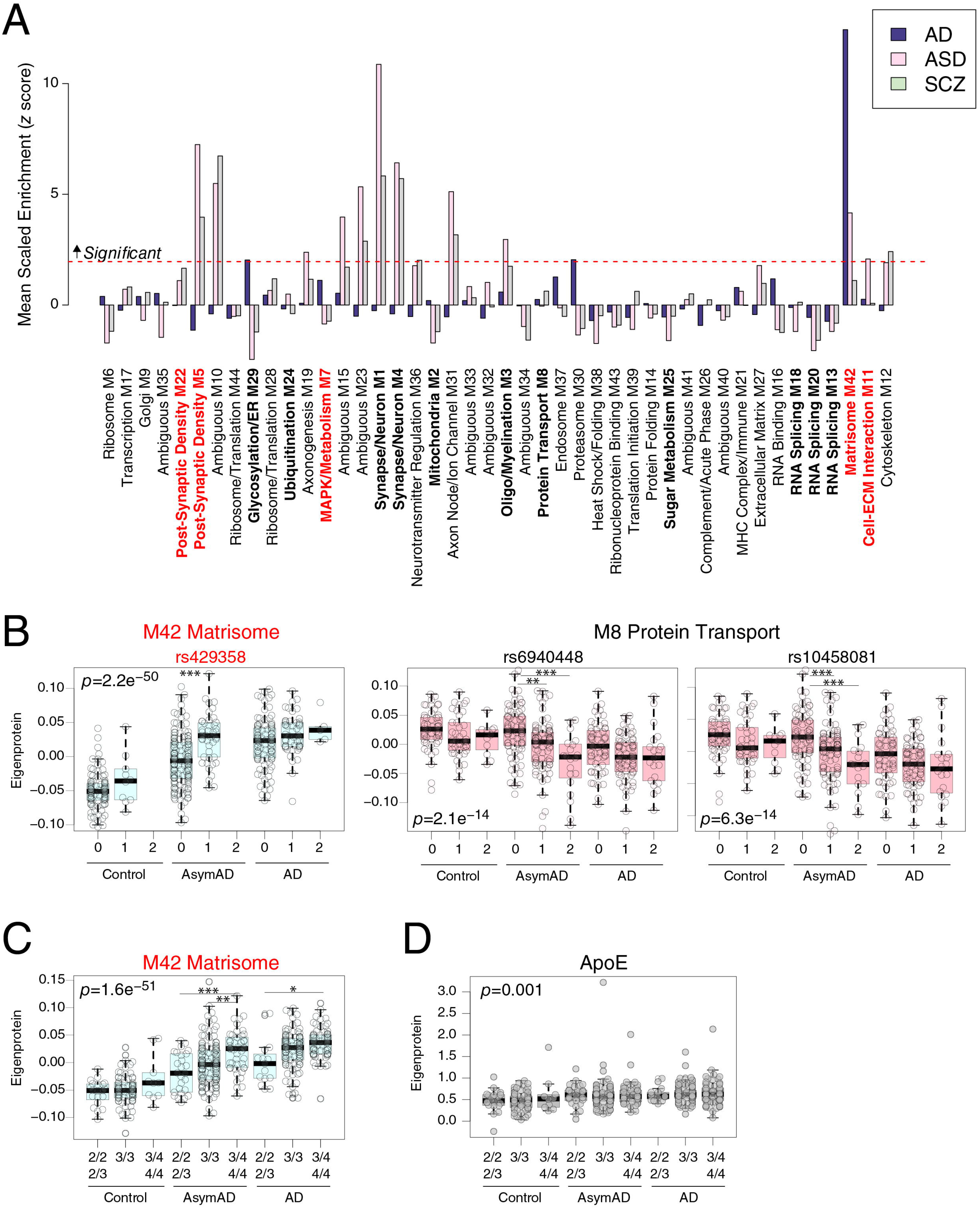
The M42 Matrisome Module is Enriched in AD Genetic Risk and is Increased by *APOE ε4*. (A, B) Enrichment of AD genetic risk factor proteins as identified by GWAS in TMT AD network modules (A). The dashed red line indicates a *Z* score of 1.96 (*p*=0.05), above which enrichment was considered significant. Enrichment in M42 is driven by ApoE. Modules are ordered by relatedness as illustrated in Figure 1B. AD, Alzheimer’s disease; ASD, autism spectrum disorder; SCZ, schizophrenia. (B) Module eigenprotein levels by allele dose (0, 1, 2) for the three SNPs identified as proximal module quantitative trait loci (mod-QTLs), separated by case status. (C) M42 matrisome module eigenprotein levels by *APOE* genotype, separated by case status. (D) ApoE levels by *APOE* genotype, separated by case status. ApoE is increased in AsymAD and AD, but *APOE* genotype does not affect ApoE levels. Full statistics are provided in **Supplementary Table 26**. Differences in eigenprotein levels were assessed by one-way ANOVA with Tukey test. Only significant differences within case status group are shown. \**p*<0.05, \*\**p*<0.01, \*\*\**p*<0.001. Boxplots represent the median, 25^th^, and 75^th^ percentiles, and box hinges represent the interquartile range of the two middle quartiles within a group. Datapoints up to 1.5 times the interquartile range from box hinge define the extent of whiskers (error bars).

While gene-based tests of association provide information on whether network modules are likely to be involved in upstream disease processes, they do not provide information on how variation in the genome influences module levels. To address this question, we performed a module quantitative trait loci (mod-QTL) analysis using genome-wide genotyping available from both ROSMAP and Banner cohorts. At a genome-wide level of significance and after adjusting for diagnosis and sex, among other variables, we found that rs429358 was a proximal mod-QTL (within 1MB, i.e., *cis*) for the M42 matrisome module (**Table 1**). rs429358 is located in the *APOE* locus and defines the *APOE ε4* allele. This mod-QTL was further evident when we plotted the M42 eigenprotein by dose of the rs429358 SNP (**Figure 4B**). Furthermore, M42 was the only module observed to vary by *APOE* genotype (**Figure 4C**). Importantly, rs429358 did not affect ApoE protein levels when tested in a linear regression model adjusting for diagnosis (*p*=0.44). This was confirmed when we analyzed ApoE protein levels by genotype and case status (**Figure 4D, Supplementary Table 26**), indicating that the change in M42 levels caused by rs429358 was independent of ApoE levels. We also observed a proximal mod-QTL for the M8 protein transport module which was near the tubulin beta-2A (TUBB2A) gene (**Table 1, Figure 4B**). There were more distal mod*-*QTLs (>1MB from module proteins, i.e., *trans*) observed than proximal mod*-*QTLs. In summary, we found three TMT AD network modules that were significantly enriched for AD genetic risk factors, and the level of one of these modules—M42 matrisome—was influenced by genetic variation in *APOE* independent of diagnosis or ApoE protein levels, especially in the asymptomatic stage of the disease.

**Table 1.** TMT AD Network Module Quantitative Trait Loci (mod-QTLs) . SNPs associated with the first eigenprotein of a protein module at a genome-wide significant level (p < 5×10^-8^) were referred to as protein co-expression module QTLs (mod-QTLs). mod-QTLs located within 1 megabase of one of the module proteins were defined as proximal (*cis*) mod-QTLs, otherwise, they were categorized as distal (*trans*) mod*-*QTLs. The associations were adjusted for cognitive diagnosis, sex, 10 genetic principal components, and genotyping chip. CHR, chromosome; SNP, single nucleotide polymorphism; BP, base pair; A1, allele; na, not applicable

| SNP | CHR | BP | A1 | Nearest Coding Gene to SNP | BETA | P value | Module | mod-QTL | Module Protein within 1Mb of SNP |
| --- | --- | --- | --- | --- | --- | --- | --- | --- | --- |
| rs28716042 | 8 | 125418451 | G | TMEM65 | 0.04 | 3.7E-09 | M3 Oligo/Myelination | *trans* | na |
| rs112028701 | 13 | 104134726 | T | SLC10A2 | -0.04 | 3.1E-08 | M22 Post-synaptic Density | *trans* | na |
| rs11021075 | 11 | 94940421 | T | SESN3 | 0.02 | 4.9E-08 | M26 Complement/Acute Phase | *trans* | na |
| rs1733609 | 7 | 81536375 | A | CACNA2D1 | 0.02 | 1.6E-08 | M11 Cell-ECM Interaction | *trans* | na |
| rs429358 | 19 | 45411941 | C | APOE | 0.02 | 3.2E-08 | M42 Matrisome | *cis* | APOE |
| rs6940448 | 6 | 3810805 | G | FAM50B | -0.02 | 1.4E-08 | M8 Protein Transport | *cis* | TUBB2A |
| rs1553484 | 6 | 91507934 | A | MAP3K7 | -0.02 | 6.4E-09 | M29 Glycosylation/ER | *trans* | na |
| rs2352535 | 4 | 114127905 | T | ANK2 | 0.02 | 2.4E-08 | M27 Extracellular Matrix | *trans* | na |

### The M7 MAPK/Metabolism Module Is Associated with Cognitive Trajectory Independent of Neuropathology in ROSMAP

The M7 MAPK/metabolism and M42 matrisome modules, among others, were strongly correlated with cognitive function as assessed near time of death. Because subjects in the ROSMAP cohort undergo in-depth and repeated cognitive testing during life, we sought to analyze whether their TMT AD network module levels as assessed post-mortem would be associated with their individual cognitive trajectories during life. We conceptualized our findings into two categories: i) modules with higher expression levels being associated with more rapid cognitive decline (i.e., a negative association between module eigenprotein and cognitive trajectory), and ii) modules with higher expression levels being associated with a slower rate of cognitive decline (i.e., a positive association between module eigenprotein and cognitive trajectory), also referred to as cognitive preservation. We constructed two linear models to analyze module associations with cognitive trajectory: one that adjusted for ten measured neuropathologies in ROSMAP (amyloid-β, tangles, cerebral amyloid angiopathy, cerebral atherosclerosis, arteriolosclerosis, Lewy bodies, TDP-43 deposits, gross infarcts, microinfarcts, and hippocampal sclerosis) to exclude the potential effect of neuropathology on the association, and one that did not adjust for neuropathology. Without adjusting for neuropathology, we found ten modules that were significantly associated with cognitive decline, and eleven modules that were significantly associated with cognitive preservation (**Table 2, Supplementary Table 27**). Modules that were most strongly associated with cognitive decline included M7 MAPK/metabolism, M15 ambiguous, and M42 matrisome, whereas modules that were strongly associated with cognitive preservation included M33 ambiguous, M5 post-synaptic density, and M2 mitochondria. After adjustment for neuropathology, the M42 matrisome, M11 cell-ECM interaction, M20 RNA splicing, and M25 sugar metabolism modules were no longer significantly associated with cognitive decline, and the M44 ribosome/translation, M32 ambiguous, and M9 golgi modules were no longer associated with cognitive preservation. M7 MAPK/metabolism and its related module M15 (ambiguous) remained significantly associated with rate of cognitive decline after adjustment, as well as five other modules, including the M24 ubiquitination module (**Figure 5A, Supplementary Table 27**). Five out of seven modules significantly associated with rate of cognitive decline were unique to the protein network. Modules that remained significantly associated with cognitive preservation after adjustment for neuropathology included the M2 mitochondria and related module M33 (ambiguous), M5 post-synaptic density, and the M29 glycosylation/ER module that was unique to the protein network (**Figure 5B, Supplementary Table 27**). To further examine the association of TMT AD network modules with cognitive trajectory, we assessed which modules were enriched in proteins either positively associated or negatively associated with cognitive resilience after adjustment for neuropathology, as identified by a recent proteome-wide association study (PWAS) of cognitive resilience in ROSMAP^46^ (**Figure 5C, Supplementary Tables 28 and 29**). In our operationalized definition, a positive association with resilience would be equivalent to cognitive preservation, whereas a negative association with cognitive resilience would be equivalent to cognitive decline. Consistent with our module association analysis, we found that M7 MAPK/metabolism and M15 ambiguous were significantly enriched in proteins negatively associated with cognitive resilience, whereas M5 post-synaptic density and M8 protein transport module were significantly enriched in proteins positively associated with cognitive resilience. Other modules, such as M2 mitochondria, M29 glycosylation/ER, and M24 ubiquitination trended towards significance. Proteins identified by PWAS that were enriched in each module are provided in **Supplementary Tables 28 and 29**.

**Figure 5.**
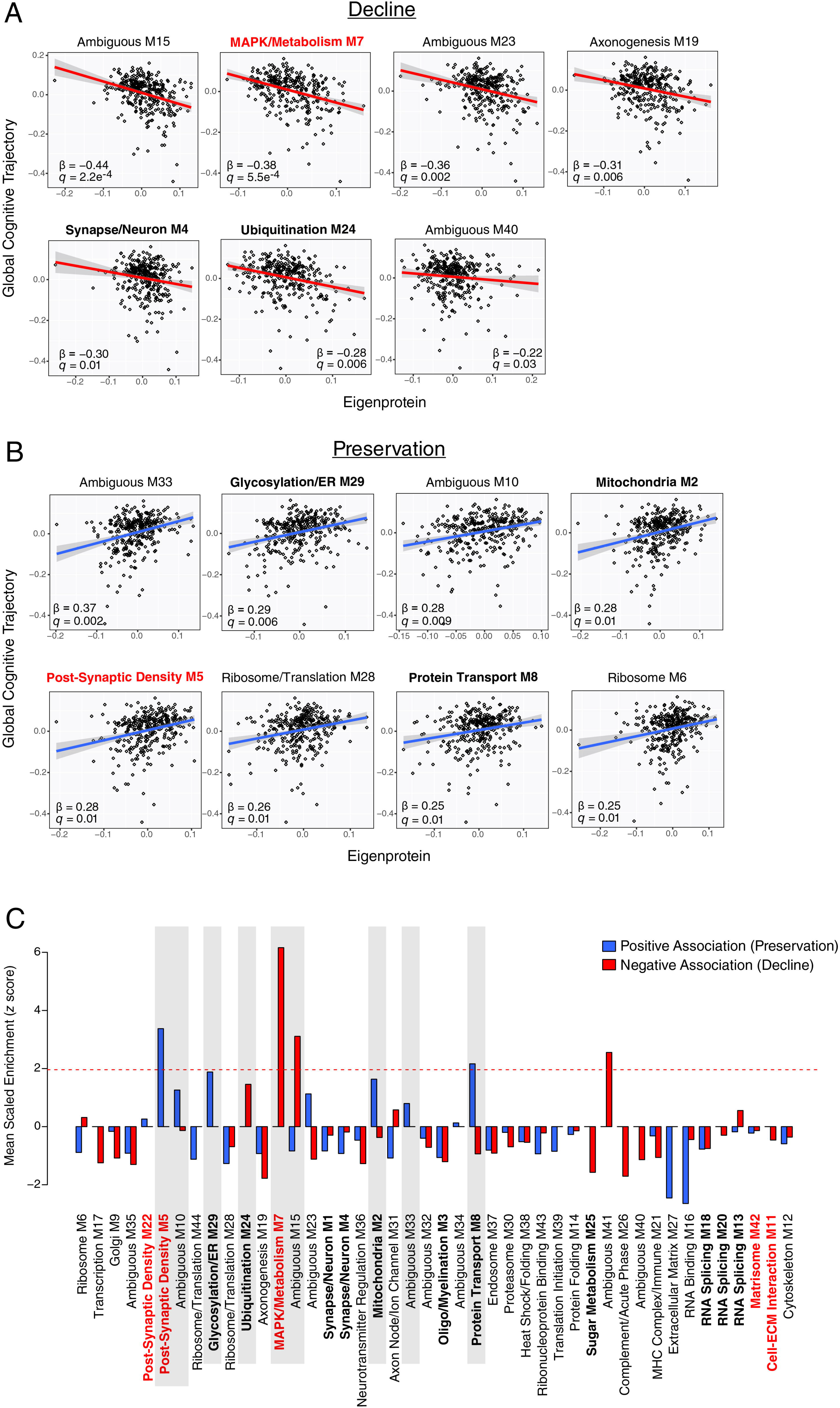
The M7 MAPK/Metabolism Module is Associated with Cognitive Decline. (A-C) TMT AD network modules associated with cognitive decline (A) or cognitive preservation (B) after adjustment for 10 neuropathologies in ROSMAP. Eigenprotein values are plotted against the rate of cognitive change during life for each subject in ROSMAP. Decline is highlighted in red, and preservation is highlighted in blue. β is the effect size of module eigenprotein on cognitive trajectory after adjustment for neuropathology; *q* is the FDR significance level of this effect. Information on the association between all TMT AD network module eigenproteins and cognitive trajectory before and after adjustment for neuropathology is provided in **Supplementary Table 27**. (C) TMT AD network module enrichment of proteins positively associated with cognitive resilience (i.e., preservation) or negatively associated with cognitive resilience (i.e., decline) identified in a prior proteome-wide association study (PWAS) of cognitive resilience in the ROSMAP cohort^46^. The dashed red line indicates a *Z* score of 1.96 (*p*=0.05), above which enrichment was considered significant. Modules that are shaded are consistent with results in (A, B). Modules are ordered by relatedness as illustrated in Figure 1B.

**Table 2.**
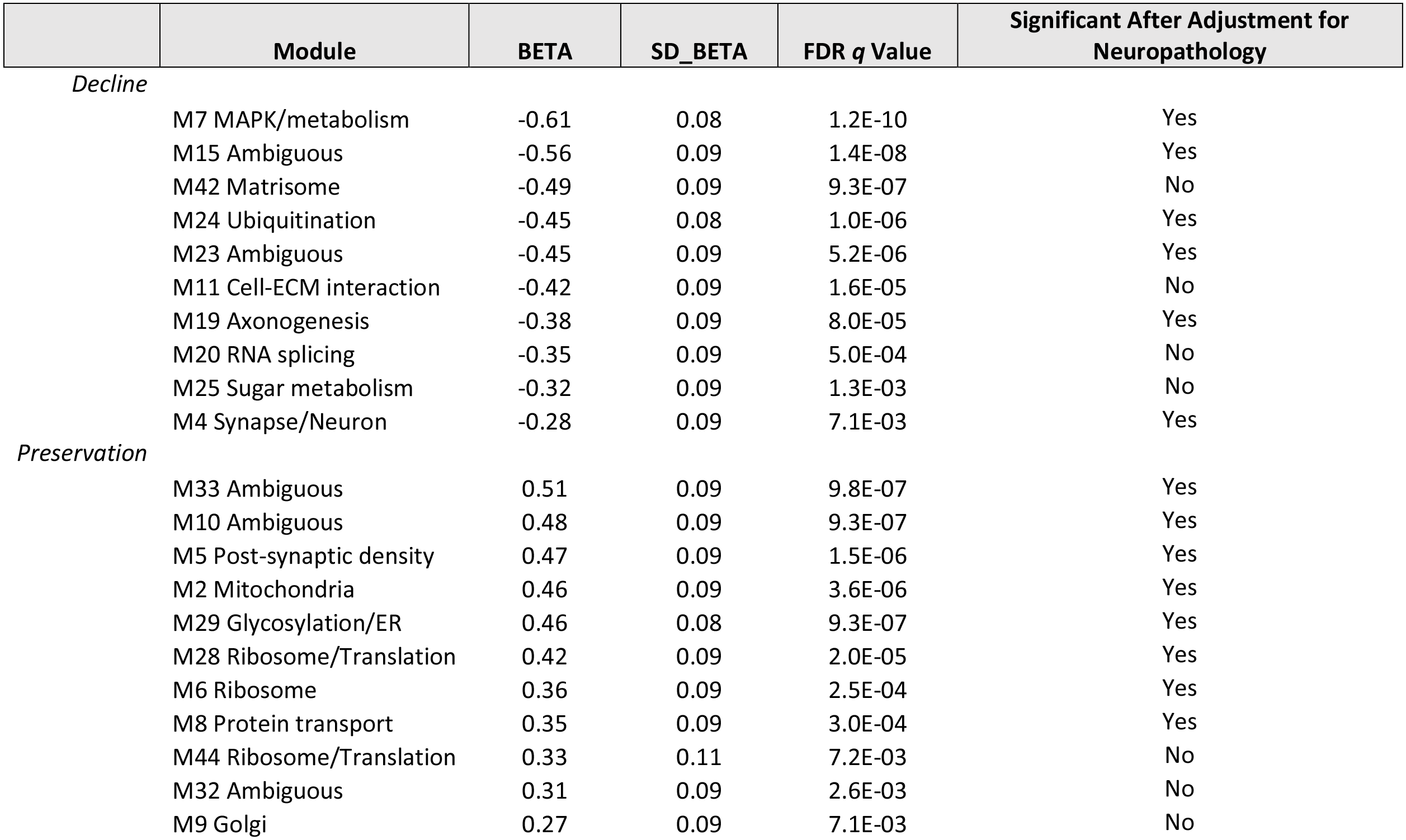
Association of TMT AD Network Modules with Cognitive Trajectory. . The association between a module eigenprotein level for each ROSMAP subject in the TMT AD network and his or her individual cognitive trajectory was modeled with and without adjustment for neuropathology. Modules that remained significantly associated with cognitive trajectory after adjustment for neuropathology are shown in **Figure 5**. Modules that had a negative association with cognitive trajectory were defined as those involved in cognitive decline, whereas modules that had a positive association were defined as those involved in cognitive preservation.

In summary, although the M42 matrisome module was enriched in genetic risk and its levels were influenced by a SNP that defines the *APOE ε4* allele, we found that it was not associated with cognitive trajectory after adjustment for neuropathology. By contrast, the M29 glycosylation/ER module, unique to the protein network, was both enriched for genetic risk and remained associated with cognitive trajectory after adjustment for neuropathology. The M7 MAPK/metabolism module demonstrated a trend towards enrichment for AD genetic risk, and was one of several modules unique to the protein network that significantly correlated with cognitive trajectory even after adjustment for neuropathology, suggesting that although M7 was enriched in Aβ plaque and NFT proteins, at least part of its effect on cognitive function was independent of neuropathology. M7 was associated with cognitive decline, and was also enriched in proteins independently observed to be associated with cognitive decline in a recent PWAS of cognitive resilience.

## Discussion

In this study, we analyzed more than 1000 brain tissues by TMT-MS across multiple centers, cohorts, and brain regions to develop a robust TMT AD protein network that significantly expanded upon our previous consensus LFQ AD network. Using a multilayered –omics approach, we identified new protein network modules strongly associated with AD that were not present in RNA-based networks. Some of these modules, such as the M7 MAPK/metabolism module, were associated with both AD neuropathology and cognitive trajectory in ROSMAP even after adjustment for neuropathology, and one of them—the M42 matrisome module—was influenced by the *APOE* locus. These findings highlight the utility of extending proteomic analytical depth to uncover additional AD-related protein co-expression network changes, as well as the value of analyzing increasingly larger cohorts with comprehensive clinical and pathological phenotyping to provide the statistical power necessary to identify disease-relevant relationships across multi-omic datasets.

The TMT AD network, which was robustly validated by a different algorithm for network generation, was able to provide additional resolution of important disease-related modules in our prior LFQ AD network. The M4 microglia/astrocyte metabolism module in the LFQ network was the most strongly correlated to AD neuropathology and cognitive function (Braak stage *r*=0.49, *p*=4.7e^-27^; mini-mental state examination *r*=–0.67, *p*=8.5e^-23^). In the TMT AD network, this module was split into two separate modules—the M7 MAPK/metabolism module, and the M11 cell-ECM interaction module. M11 contained nearly all LFQ M4 hub proteins such as moesin (MSN), plectin (PLEC), and CD44, whereas M7 contained LFQ M4 proteins with lower module correlation (kME) that now were hubs in M7, such as MAPK1, MAPK3, and leukotriene A-4 hydrolase (LTA4H). While both modules were significantly correlated with AD neuropathology and cognitive function, the M7 MAPK/metabolism module was more strongly correlated with cognitive function and was also associated with cognitive trajectory after adjustment for neuropathology, whereas the M11 cell-ECM interaction module was not associated with cognitive trajectory after adjustment for neuropathology. M11 was highly related to the M42 matrisome module, which was enriched in AD genetic risk primarily due to ApoE being a member of the module, with both modules containing a mod-QTL that influenced their levels. For the M42 matrisome module, this mod-QTL was associated with the *APOE* locus, the strongest genetic risk factor for sporadic AD. Similarly to M11, M42 was not associated with cognitive trajectory after adjustment for its association with neuropathology. Given these observations, one could therefore postulate a model in which the pathophysiology embodied by M11 and M42 is necessary for subsequent downstream AD pathological changes, but that the pathological changes most closely associated with cognitive decline, such as those represented by the M7 MAPK/metabolism module among others, are the prime effectors and/or modulators of cognitive decline in AD. Targeting both types of AD pathophysiology holds promise in the context of AD therapeutic development.

The TMT AD network was able to reveal a module of proteins strongly associated with Aβ—M42. We did not observe a module as strongly associated with tau in the TMT AD network because of the stark differences in peptide levels between N-terminal and microtubule-binding regions of tau in AD brain which, when averaged together into a total microtubule associated protein tau (MAPT) quantitation, dilute significant changes in the measured levels of this protein by mass spectrometry. We addressed this technical issue by quantifying the microtubule binding region (MTBR) and non-MTBR tau domains in a separate database search, and correlated each module to these tau “proteins.” We found that M42 matrisome and M11 cell-ECM correlated most strongly to the MTBR. In contrast to quantitation of MAPT, levels of the amyloid precursor protein (APP) in AD are driven by changes in the level of peptides from the Aβ region of the protein. Therefore, when analyzing AD cases, APP can be considered a surrogate marker for Aβ^4, 47, 48^. APP co-expressed with proteins in the M42 matrisome module. M42, which was not present in RNA networks, contained a number of proteins that have previously been identified by TMT-MS and shown to be correlated with Aβ^10, 28, 29, 49^. These proteins, such as MDK, NTN1, and SMOC1, appear to be less reliably detected by LFQ-MS even when using LCM to isolate plaques from surrounding brain parenchyma for MS analysis, likely reflecting their lower relative abundance to other proteins within Aβ plaques or their potential weak binding affinity to plaques that may be disrupted during tissue fixation. MDK and NTN1 have previously been shown to bind directly to Aβ^28^. Interestingly, many of the M42 proteins contain heparin, heparan sulfate, and glycosaminoglycan binding domains that may mediate their interaction with Aβ plaques. ApoE, a member of the M42 module, has also been shown to interact with heparan sulfate proteoglycans, and loss of this binding interaction has been suggested as a possible mechanism for the remarkable protection afforded by the ApoE Christchurch loss of function mutation recently described in a presenilin-1 autosomal dominant AD mutation carrier^50, 51^. The *APOE ε4* allele has been associated with an increase in matrisome pathways in mixed neuronal-glial induced pluripotent stem cell cultures^52^. Other proteins in M42 may influence Aβ plaque pathophysiology through different mechanisms, such as secreted frizzled-related protein 1 (SFRP1), which modulates Wnt signaling, and has been shown to inhibit the disintegrin and metalloproteinase domain-containing protein 10 (ADAM10) that is important for regulation of Notch signaling and Aβ metabolism^39, 53, 54^. To what extent modulation of M42 protein levels, enzymatic activity, or protein-Aβ/protein-proteoglycan interactions may affect Aβ plaque deposition or its downstream consequences remains to be determined, but such proteins represent promising therapeutic targets for AD. They may also represent promising biofluid AD biomarkers. Indeed, we have recently shown that SMOC1 is strongly elevated in AD cerebrospinal fluid (CSF)^28, 49, 55^. Consistent with our mod-QTL findings, levels of SMOC1 in CSF were recently identified to be influenced by a pQTL associated with the *APOE ε4* allele^56^.

Interestingly, while the M42 matrisome module was enriched in genes that fall within AD risk loci, it was also enriched in genetic risk for autism spectrum disorder (ASD), suggesting a key role for the matrisome in both diseases. Proper function of the matrisome is known to be important for cortical patterning during development^57, 58^, and remodeling of the extracellular matrix by microglia is important for synaptic plasticity and experiential learning^59^. Disruption of the matrisome by Aβ plaques and/or the response of microglia to such disruption may be a potential mechanism by which cognitive dysfunction occurs in AD. A module similar to M11 cell-ECM interaction that contains MSN as a hub protein has been shown to be increased in ASD^60^. The interplay between proper matrisome structure and microglia function in the context of neural development, learning, and memory is an area that deserves further study.

The M7 MAPK/metabolism module was most strongly correlated to cognitive function in the TMT AD network, and along with the highly related M15 module of ambiguous ontology, was also the most strongly correlated to cognitive decline before and after adjustment for AD neuropathology. Like M42, M7 was unique to the protein network with no cognate module in RNA networks, and had stronger correlation to cognitive function than any module in the ROSMAP RNA network. This module represented the stress-activated MAPK cascade through hub proteins such as MAPK1/ERK2, MAPK3/ERK1, and ribosomal protein S6 kinase alpha-2 (RPS6KA2), but it also contained the hub proteins platelet-activating factor acetylhydrolase IB subunit alpha1 (PAFAH1B3) and LTA4H—proteins both involved in cellular lipid metabolism^61, 62^; ubiquitin-like modifier-activating enzyme 1 (UBA1), which catalyzes the first step in ubiquitin conjugation to mark cellular proteins for degradation through the ubiquitin-proteasome system^63^; mothers against decapentaplegic homolog 4 (SMAD4), part of nuclear SMAD complex required for transforming growth factor-β (TGF-β) signaling^64^; puromycin-sensitive aminopeptidase (NPEPPS), which degrades tau^65^; regulator of nonsense transcripts (UPF2) important for the nonsense-mediated decay (NMD) of mRNA transcripts that contain a premature stop codon^66^; and chromogranin-A (CHGA), which is important for immune and host defense responses^67^. It also contained many proteins involved in glycolytic sugar metabolism. Therefore, this module could be viewed as representing activation of a “stress-response” program, likely at least in part to pathologic Aβ and tau deposits as suggested by its spatial overlap with Aβ plaques and NFTs, with consequent shift in cellular metabolism from oxidative phosphorylation to glycolysis. This metabolic shift is clearly evident in CSF, as many of the most strongly elevated proteins in AD CSF are involved in glycolysis^3, 49, 68, 69^. The brain cell types that drive this shift are presumably microglia and/or astrocytes. Indeed, upregulation of microglial glycolytic metabolism is a key event in response to plaques, and is disrupted by mutations in the triggering receptor expressed on myeloid cells 2 (TREM2) protein that are strong risk factors for development of AD^70, 71^. Furthermore, single-cell RNA sequencing of astrocytes in humanized ApoE4 mice shows increased levels of proteins involved in glycolysis compared to ApoE3 mice, suggesting that astrocyte metabolism may be altered in AD. However, in contrast to our previous findings from LFQ network in which the sugar metabolism module was strongly enriched in microglia and astrocyte protein markers, in the TMT network the M7 and M25 sugar metabolism module did not have strong cell type character. Further elucidation of cell type metabolic shifts in AD will be facilitated by single-cell and single cell-type proteomics.

As part of the stress-response program, M7 contains a number of proteins that have been shown to play important roles in the metabolism of misfolded proteins, including NPEPPS, UBA1, and valosin-containing protein (VCP). VCP was recently identified as a tau disaggregase, with a hypomorph mutation leading to dementia with vacuolar changes and neurofibrillary tangle deposition^72^. UPF2, as noted above, plays an important role in mRNA surveillance and the degradation of transcripts by NMD that contain premature stop codons through mutations or mis-splicing events. We have previously shown that aggregation of RNA binding proteins that are involved in mRNA splicing is an early event in AD pathogenesis, and likely leads to aberrant mRNA splicing events that may give rise to premature stop codons or alternative exon-exon junctions^10, 47, 48, 73–75^. These proteins, such as U1 small nuclear ribonucleoprotein 70 kDa (SNRNP70, or U1-70K) and others that are part of the U1 spliceosome complex, have been shown to interact with NFTs through their low-complexity basic-acidic domains, and were strongly enriched in the M13 RNA splicing module^76^. Module M20 RNA splicing, of which putative RNA-binding protein luc7-like 1 (LUC7L) was the top member by kME, was highly related to M13 and was more strongly associated with AD neuropathological traits and cognitive function. The effect of tau misfolding on mRNA splicing continues to be an area of active study^77^. Interestingly, we did not observe a well-defined M7 module in the 168-case ROSMAP network built for direct protein-RNA comparisons, perhaps due to the likely association of M7 with tau-related pathophysiology and the fact that there were only two cases with advanced tau pathology (Braak VI) in this cohort. Indeed, only three cases with Braak VI pathology were present in the full ROSMAP cohort used for construction of the TMT AD network, whereas there were 28 cases with Braak VI pathology in the smaller Banner cohort. This demonstrates the value of analyzing multiple cohorts together in a consensus network, which helps to de-emphasize changes that are unique to any one cohort in the overall analysis.

In our analysis of the association between module levels and cognitive trajectory, we observed that M7 remained associated with cognitive decline after adjustment for neuropathology, suggesting that the changes in this module in AD may not be due solely to a response to plaques and tangles. Indeed, while M7 trended towards an increase in AsymAD, the change was not significant from control. This observation, along with its association with cognitive decline, and the fact that many of the M7 hub proteins such as UBA1 were independently associated with cognitive decline in a PWAS study of cognitive resilience^46^, would suggest that increased M7 MAPK/metabolism levels would be detrimental to cognitive function. However, as noted previously, robust microglial metabolic function is important for a beneficial stress response to amyloid plaques^71^, and also appears to be important for maintenance of cognitive function during aging^78^. Furthermore, M7 trended toward enrichment in AD genetic risk in this study, suggesting that perhaps loss of function of M7 is detrimental to cognitive function. This apparent paradox was also observed in our previous LFQ study, in which increased levels of the M4 astrocyte/microglia metabolism module—the parent module of M7 and M11—were strongly associated with reduced cognitive function, yet many key M4 proteins were noted to be associated with beneficial inflammatory responses in mouse models and decreased in cases of rapidly progressive AD. M4 was also enriched in AD genetic risk. Therefore, the stress response embodied by M7 may serve both beneficial and detrimental roles in AD, and determining which aspects of a potential beneficial response to augment, or detrimental response to inhibit, will likely require direct modulation experiments in appropriate animals models or human clinical trials, as well as biomarkers to measure such a response. Further study and modulation of the biological response represented by M7 represents a key goal in AD therapeutic development.

In addition to modules that were associated with cognitive decline, we identified modules that were associated with cognitive preservation. We operationally defined cognitive preservation as the positive correlation between module eigenprotein and cognitive trajectory. Modules associated with cognitive preservation included M5 post-synaptic density and M2 mitochondrial, as well as M29 glycosylation/ER that was unique to the protein network and enriched in AD genetic risk, suggesting that perhaps loss of M29 function may be detrimental to cognitive function. Neuritin (NRN1), rabphilin-3A (RPH3A), and neurosecretory protein VGF were previously identified by PWAS to be key proteins associated with cognitive resilience^46, 79^, and all are members of the M5 post-synaptic density module. Mitochondrial homeostasis has been associated with cognitive preservation^3, 79^. Interestingly, the module most strongly associated with cognitive preservation was M33, which did not have a coherent ontology, but was enriched for AD genetic risk to nearly the same level as M7. The hub protein with strongest kME to M33 was parvalbumin (PVALB), which is a marker for a class of GABA-ergic inhibitory interneurons that are important for generating gamma oscillatory frequencies in the brain and normal neuronal network function^80^. Loss or dysfunction of PVALB-positive interneurons has been suggested as an important driver of AD cognitive dysfunction in human and animal models^81, 82^, and it is therefore possible that M33 represents protein co-expression related to this brain cell type. Further development of cell type specific marker lists derived from proteomic experiments would be useful to dissect potential module overlap with additional cellular subtypes beyond the primary types assessed in this study, as well as eventual direct interrogation through single-cell proteomics to define such potential cell subtype changes at the proteomic level^83, 84^.

We were able to leverage both genomic and transcriptomic data in this study to better understand how proteomics data converge or diverge from these other –omics approaches. We observed AD genetic risk enrichment in M42 matrisome, M30 proteasome, and M29 glycosylation/ER modules, and a mod-QTL for M42 that showed strong genetic association with the *APOE* locus.

In our LFQ consensus analysis, we observed enrichment of AD genetic risk in the LFQ M2 myelin/oligodendrocyte and M4 astrocyte/microglia metabolism modules. In the TMT network, the cognate M3 oligo/myelination module did not reach statistical significance for enrichment of AD genetic risk, likely due to its slightly different module protein membership such as including a splice variant of the AD risk factor protein Myc box-dependent-interacting protein 1 (BIN1) rather than the canonical BIN1 protein. We did, however, observe a distal mod-QTL for M3 that mapped most closely to the transmembrane 65 (TMEM65) protein. TMEM65 is involved in proper mitochondrial respiration, and mutations in this protein have been associated with mitochondrial encephalomyopathy^85^. Although M3 was not as strongly correlated to AD traits as some other modules and was not elevated in AsymAD—as also observed in the LFQ network—alterations in this module may nonetheless play an important role in setting the stage for subsequent cognitive decline^86, 87^, similar to the association of the *APOE ε4* allele with the M42 matrisome module identified by mod-QTL analysis.

Our study demonstrates the importance of analyzing proteins directly in addition to their coding transcripts. We found only a modest correlation between mRNA and protein level changes in AD, consistent with previous findings on the relationship of mRNA to protein levels, even at the single cell level^83, 84, 88^. Such differences are likely caused by many factors, including both translational and post-translational regulation, as well as perhaps differences in sampling due to cell structure or polarization. The concordance between mRNA and protein co-expression was better than differential expression, yet nearly half of the modules in the TMT AD protein network were not preserved in RNA networks from the same brain region. Modules that were preserved tended to reflect cell and organelle types, such as the M1 synapse/neuron, M2 mitochondria, M3 oligo/myelination, and M5 post-synaptic density modules. Modules that were less well preserved tended to include processes related to abnormal protein processing and metabolism, including M7 MAPK/metabolism, M13 RNA splicing, M24 ubiquitination, and M42 matrisome modules. As might be expected for a disease defined by cognitive decline in the presence of characteristic protein dysmetabolism, this observation indicates that a significant proportion of biological changes relevant to AD pathophysiology are occurring through mechanisms that are not reflected through changes in mRNA abundance or co-expression, and highlights the importance of integrating multiple levels of –omics data to further our understanding of the disease.

## Supporting information

Johnson et al 2021 Extended Data

Johnson et al 2021 Supplementary Tables Rev1

## Acknowledgements

We are grateful to those who agreed to donate their brains for research and who participated in the described observational studies. This study was supported by the following National Institutes of Health funding mechanisms: RF1AG057471, RF1AG057470, R01AG01581, U54AG065187, U01AG061357, R01AG057911, R01AG061800, R01AG053960, RF1AG062181, P30AG10161, R01AG15819, R01AG17917, U01AG61356, RF1AG057440, K08AG068604, U24NS072026, P30AG19610, and R01AG056533. The study was also supported by VA Administration BX005219, Arizona Department of Health Services (contract 211002, Arizona Alzheimer’s Research Center), the Arizona Biomedical Research Commission (contracts 4001, 0011, 05-901 and 1001 to the Arizona Parkinson’s Disease Consortium) and the Michael J. Fox Foundation for Parkinson’s Research. We thank Kaiwen Yu for his contributions to this study. We also thank the Neuropathology Core of the Emory Center for Neurodegenerative Disease Core Facilities for their assistance.

## Data and Code Availability

Raw data, case traits, and analyses related to this manuscript are available at https://www.synapse.org/DeepConsensus. The results published here are in whole or in part based on data obtained from the AMP-AD Knowledge Portal (https://adknowledgeportal.synapse.org). The AMP-AD Knowledge Portal is a platform for accessing data, analyses and tools generated by the AMP-AD Target Discovery Program and other programs supported by the National Institute on Aging to enable open-science practices and accelerate translational learning. The data, analyses and tools are shared early in the research cycle without a publication embargo on secondary use. Data are available for general research use according to the following requirements for data access and data attribution (https://adknowledgeportal.synapse.org/#/DataAccess/Instructions). ROS/MAP resources can be requested at www.radc.rush.edu. The algorithm used for batch correction is fully documented and available as an R function, which can be downloaded from https://github.com/edammer/TAMPOR.

## Author Contributions

ECBJ, EKC, EBD, DMD, RB, LP, TSW, APW, JP, and NTS designed experiments; DMD, LP, RB, LP, and LY carried out experiments; ECBJ, EKC, EBD, ESG, YL, JL, TSW, APW, and NTS analyzed data; DMD, RB, and LP provided advice on the interpretation of data; ECBJ wrote the manuscript with input from coauthors; TGB, CG, DAB, PLD, VH, BZ, and MG provided tissue samples; AIL, JJL, and NTS supervised the study. All authors approved the final manuscript.

## Methods

### Brain Tissue Samples and Case Classification

Brain tissue used in this study was obtained from the autopsy collections of the Banner Sun Health Research Institute^15^, Mount Sinai School of Medicine Brain Bank, Religious Orders Study and Rush Memory and Aging Project^89^, and Emory Alzheimer’s Disease Research Center. Tissue was from the dorsolateral prefrontal cortex (Brodmann area 9), frontal cortex (Brodmann area 6 and 10), anterior cingulate (Brodmann area 24), temporal cortex (Brodmann area 37), or parahippocampal gyrus (Brodmann area 36), as indicated. Human postmortem tissues were acquired under proper Institutional Review Board (IRB) protocols at each respective institution. Postmortem neuropathological evaluation of neuritic plaque distribution was performed according to the Consortium to Establish a Registry for Alzheimer’s Disease (CERAD) criteria^17^, while extent of spread of neurofibrillary tangle pathology was assessed with the Braak staging system^16^. Other neuropathologic diagnoses were made in accordance with established criteria and guidelines^90^. All case metadata are provided in **Supplementary Tables 1, 13-16, and 30**. Case classification harmonization across cohorts was performed using the following rubric: cases with CERAD 0-1 and Braak 0-3 without dementia at last evaluation were defined as control (if Braak equals 3, then CERAD must equal 0); cases with CERAD 1-3 and Braak 3-6 without dementia at last evaluation were defined as AsymAD; cases with CERAD 2-3 and Braak 3-6 with dementia at last evaluation were defined as AD. Dementia was defined as MMSE <24, or CDR ≥1, based on prior comparative study^91^.

### Brain Tissue Homogenization and Protein Digestion

For ROSMAP and Banner tissues, procedures were performed essentially as described^4, 29^. Approximately 100 mg (wet tissue weight) of brain tissue was homogenized in 8 M urea lysis buffer (8 M urea, 10 mM Tris, 100 mM NaH2PO4, pH 8.5) with HALT protease and phosphatase inhibitor cocktail (ThermoFisher) using a Bullet Blender (NextAdvance). Each Rino sample tube (NextAdvance) was supplemented with ∼100 μL of stainless steel beads (0.9 to 2.0 mm blend, NextAdvance) and 500 μL of lysis buffer. Tissues were added immediately after excision and homogenized with bullet blender at 4 °C with 2 full 5 min cycles. The lysates were transferred to new Eppendorf Lobind tubes and sonicated for 3 cycles consisting of 5 s of active sonication at 30% amplitude, followed by 15 s on ice. Samples were then centrifuged for 5 min at 15,000 x *g* and the supernatant transferred to a new tube. Protein concentration was determined by bicinchoninic acid (BCA) assay (Pierce). For protein digestion, 100 μg of each sample was aliquoted and volumes normalized with additional lysis buffer. Samples were reduced with 1 mM dithiothreitol (DTT) at room temperature for 30 min, followed by 5 mM iodoacetamide (IAA) alkylation in the dark for another 30 min. Lysyl endopeptidase (Wako) at 1:100 (w/w) was added and digestion allowed to proceed overnight. Samples were then 7-fold diluted with 50 mM ammonium bicarbonate. Trypsin (Promega) was then added at 1:50 (w/w) and digestion was carried out for another 16 h. The peptide solutions were acidified to a final concentration of 1% (vol/vol) formic acid (FA) and 0.1% (vol/vol) trifluoroacetic acid (TFA), and desalted with a 30 mg HLB column (Oasis). Each HLB column was first rinsed with 1 mL of methanol, washed with 1 mL 50% (vol/vol) acetonitrile (ACN), and equilibrated with 2×1 mL 0.1% (vol/vol) TFA. The samples were then loaded onto the column and washed with 2×1 mL 0.1% (vol/vol) TFA. Elution was performed with 2 volumes of 0.5 mL 50% (vol/vol) ACN. An equal amount of peptide from each sample was aliquoted and pooled as the global pooled internal standard (GIS), which was split and labeled in each TMT batch as described below. This was performed separately for each cohort except for the ROSMAP BA6 and BA37 tissues, which were batched together and shared a GIS at the protein level before digestion. Procedures for tissue homogenization of the Mt. Sinai and Emory cohorts were performed as previously described^9, 28^.

### Isobaric Tandem Mass Tag (TMT) Peptide Labeling

Prior to TMT labeling, cases were randomized by covariates (age, sex, PMI, diagnosis, etc.), into the appropriate number of batches. Peptides from each individual case and the GIS pooled standard or bridging sample (at least 1 per batch) were labeled using the TMT 10-plex kit (ThermoFisher 90406) for ROSMAP BA9 tissues, and TMT 10-plex kit plus channel 11 (131C, lot #SJ258847) for ROSMAP BA6/BA37 and Banner tissues. In each batch, up to 2 TMT channels were used to label GIS standards, while the remaining TMT channels were reserved for individual samples following randomization. Labeling was performed as previously described^9, 10, 29^. Briefly, each sample (containing 100 μg of peptides) was re-suspended in 100 mM TEAB buffer (100 μL). The TMT labeling reagents (5mg) were equilibrated to room temperature, and anhydrous ACN (256 μL) was added to each reagent channel. Each channel was gently vortexed for 5 min, and then 41 μL from each TMT channel was transferred to the peptide solutions and allowed to incubate for 1 h at room temperature. The reaction was quenched with 5% (vol/vol) hydroxylamine (8 μl) (Pierce). All channels were then combined and dried by SpeedVac (LabConco) to approximately 150 μL and diluted with 1 mL of 0.1% (vol/vol) TFA, then acidified to a final concentration of 1% (vol/vol) FA and 0.1% (vol/vol) TFA. Labeled peptides were desalted with a 200 mg C18 Sep-Pak column (Waters). Each Sep-Pak column was activated with 3 mL of methanol, washed with 3 mL of 50% (vol/vol) ACN, and equilibrated with 2×3 mL of 0.1% TFA. The samples were then loaded and each column was washed with 2×3 mL 0.1% (vol/vol) TFA, followed by 2 mL of 1% (vol/vol) FA. Elution was performed with 2 volumes of 1.5 mL 50% (vol/vol) ACN. The eluates were then dried to completeness using a SpeedVac.

For the Emory cohort, an aliquot equivalent to 20 μg was taken from each sample and combined to make one GIS per brain region. All peptide mixtures were dried under vacuum. For each tissue region, TMT 10-plex kits (ThermoFisher) were used to label the 40 samples and 10 GIS mixtures. In each batch, TMT channels 126 and 131 were used to label GIS standards, while the 8 middle TMT channels were used to label 2 samples from each disease state, as previously described^29^. Labeling was performed according to the manufacturer’s protocol. Briefly, each sample (containing 80 μg of peptides) was resuspended in 100 mM TEAB buffer (100 μL). The TMT labeling reagents were equilibrated to room temperature, and anhydrous acetonitrile (41 μL) was added to each reagent channel and softly vortexed for 5 min. Peptide suspensions were transferred to the corresponding TMT channels and incubated for 1 h at room temperature. The reaction was quenched with 5% (vol/vol) hydroxylamine (8 μl). To ensure complete labeling, select channels from each batch were analyzed by LC-MS/MS according to previously published methods^92^. All 10 channels were then combined and dried by vacuum to ∼500 μL. Sep-Pak desalting was performed, and the eluate then dried to completeness using a SpeedVac.

For the Mt. Sinai cohort, prior to TMT labeling, the 198 samples were randomized by co-variates (protein quality, sample concentration, diagnosis, age and sex) into 20 batches (10 cases per batch)^28^. The digested peptides were resuspended in 50 mM HEPES (pH 8.5), and labeled with the TMT 11-plex kit (ThermoFisher) according to the manufacturer’s protocol. In each batch, TMT channel 126 was used to label GIS samples. All 11 channels were mixed equally and desalted with a 100 mg C18 Sep-Pak column (Waters) for subsequent fractionation.

### High-pH Off-line Fractionation

High pH fractionation was performed essentially as described^9, 93^ with slight modification. Dried samples were re-suspended in high pH loading buffer (0.07% vol/vol NH_4_OH, 0.045% vol/vol FA, 2% vol/vol ACN) and loaded onto an Agilent ZORBAX 300 Extend-C18 column (2.1mm x 150 mm with 3.5 µm beads). An Agilent 1100 HPLC system was used to carry out the fractionation. Solvent A consisted of 0.0175% (vol/vol) NH_4_OH, 0.01125% (vol/vol) FA, and 2% (vol/vol) ACN; solvent B consisted of 0.0175% (vol/vol) NH_4_OH, 0.01125% (vol/vol) FA, and 90% (vol/vol) ACN. The sample elution was performed over a 58.6 min gradient with a flow rate of 0.4 mL/min. The gradient consisted of 100% solvent A for 2 min, then 0% to 12% solvent B over 6 min, then 12% to 40 % over 28 min, then 40% to 44% over 4 min, then 44% to 60% over 5 min, and then held constant at 60% solvent B for 13.6 min. A total of 96 individual equal volume fractions were collected across the gradient and subsequently pooled by concatenation^93^ into 24 fractions and dried to completeness using a SpeedVac. Off-line fractionation of the Mt. Sinai and Emory cohorts was performed as previously described^9, 28^.

### TMT Mass Spectrometry

All fractions were resuspended in an equal volume of loading buffer (0.1% FA, 0.03% TFA, 1% ACN) and analyzed by liquid chromatography coupled to tandem mass spectrometry essentially as described^94^, with slight modifications. Peptide eluents were separated on a self-packed C18 (1.9 μm, Dr. Maisch, Germany) fused silica column (25 cm × 75 μM internal diameter (ID); New Objective, Woburn, MA) by a Dionex UltiMate 3000 RSLCnano liquid chromatography system (ThermoFisher Scientific) for the ROSMAP samples, and an Easy-nanoLC system (ThermoFisher Scientific) for the Banner samples. ROSMAP peptides were monitored on an Orbitrap Fusion mass spectrometer (ThermoFisher Scientific), and Banner peptides were monitored on an Orbitrap HF-X mass spectrometer (ThermoFisher Scientific). For ROSMAP BA9 samples, elution was performed over a 180 min gradient with flow rate at 225 nL/min. The gradient was from 3% to 7% buffer B over 5 min, then 7% to 30% over 140 min, then 30% to 60% over 5 min, then 60% to 99% over 2 min, then held constant at 99% solvent B for 8 min, and then back to 1% B for an additional 20 min to equilibrate the column. Buffer A was water with 0.1% (vol/vol) formic acid, and buffer B was 80% (vol/vol) acetonitrile in water with 0.1% (vol/vol) formic acid. For ROSMAP BA6/BA37 samples, sample elution was performed over a 120 min gradient with flow rate of 300 nL/min with buffer B ranging from 1% to 50% (buffer A: 0.1% formic acid in water, buffer B: 0.1% formic acid in 80% ACN). The mass spectrometer was set to acquire in data dependent mode using the top speed workflow with a cycle time of 3 seconds. Each cycle consisted of 1 full scan followed by as many MS/MS (MS2) scans that could fit within the time window. For ROSMAP BA9 tissues, the full scan (MS1) was performed with an m/z range of 350-1500 at 120,000 resolution (at 200 m/z) with AGC set at 4×10^5^ and maximum injection time 50 ms. The most intense ions were selected for higher energy collision-induced dissociation (HCD) at 38% collision energy with an isolation of 0.7 m/z, a resolution of 30,000, an AGC setting of 5×10^4^, and a maximum injection time of 100 ms. Five of the 50 TMT batches were run on the Orbitrap Fusion mass spectrometer using the SPS-MS3 method as previously described^29^. For ROSMAP BA6/BA37 tissues, full MS scans were collected at a resolution of 120,000 (400-1400 m/z range, 4×10^5 AGC, 50 ms maximum ion injection time). All higher energy collision-induced dissociation (HCD) MS/MS spectra were acquired at a resolution of 60,000 (1.6 m/z isolation width, 35% collision energy, 5×10^4^ AGC target, 50 ms maximum ion time). Dynamic exclusion was set to exclude previously sequenced peaks for 20 seconds within a 10-ppm isolation window. For Banner samples, elution was performed over a 120 min gradient at a flow rate of 300 nL/min with buffer B ranging from 1% to 40% (buffer A: 0.1% formic acid in water, buffer B: 0.1 % formic acid in ACN). The mass spectrometer was set to acquire data in positive ion mode using data-dependent acquisition. Each cycle consisted of one full MS scan followed by a maximum of 10 MS/MS scans. Full MS scans were collected at a resolution of 120,000 (350-1500 m/z range, 3×10^6^ AGC, 50 ms maximum ion injection time). All higher energy collision-induced dissociation (HCD) MS/MS spectra were acquired at a resolution of 45,000 (0.7 m/z isolation width, 35% collision energy, 1×10^5^ AGC target, 96 ms maximum ion time). Dynamic exclusion was set to exclude previously sequenced peaks for 20 seconds within a 10-ppm isolation window. TMT mass spectrometry of the Mt. Sinai and Emory cohorts was performed as previously described^9, 28^.

### Database Searches and Protein Quantification

All RAW files (1,200 RAW files generated from 50 TMT 10-plexes for ROSMAP BA9 tissues; 624 RAW files generated from 26 TMT 11-plexes for ROSMAP BA6/BA37 tissues; 528 RAW files generated from 22 TMT 11-plexes for Banner tissues; 760 RAW files generated from 19 TMT 11-plexes for Mt. Sinai tissues; 210 RAW files generated from 10 TMT 10-plexes for Emory tissues) were analyzed using the Proteome Discoverer suite (version 2.3, ThermoFisher Scientific). MS2 spectra were searched against the UniProtKB human proteome database containing both Swiss-Prot and TrEMBL human reference protein sequences (90,411 target sequences downloaded April 21, 2015), plus 245 contaminant proteins. The Sequest HT search engine was used and parameters were specified as follows: fully tryptic specificity, maximum of two missed cleavages, minimum peptide length of 6, fixed modifications for TMT tags on lysine residues and peptide N-termini (+229.162932 Da) and carbamidomethylation of cysteine residues (+57.02146 Da), variable modifications for oxidation of methionine residues (+15.99492 Da) and deamidation of asparagine and glutamine (+0.984 Da), precursor mass tolerance of 20 ppm, and a fragment mass tolerance of 0.05 Da for MS2 spectra collected in the Orbitrap (0.5 Da for the MS2 from the SPS-MS3 batches). Percolator was used to filter peptide spectral matches (PSMs) and peptides to a false discovery rate (FDR) of less than 1%. Following spectral assignment, peptides were assembled into proteins and were further filtered based on the combined probabilities of their constituent peptides to a final FDR of 1%. Multi-consensus was performed to achieve parsimony across individual batches. In cases of redundancy, shared peptides were assigned to the protein sequence in adherence with the principles of parsimony. As default, the top matching protein or “master protein” is the protein with the largest number of unique peptides and with the smallest value in the percent peptide coverage (i.e., the longest protein). In cases where more than one isoforms were scored equally, the additional “candidate master proteins” can be queried from the .pdresult file provided on the https://www.synapse.org/DeepConsensus. Reporter ions were quantified from MS2 or MS3 scans using an integration tolerance of 20 ppm with the most confident centroid setting. Only unique and razor (i.e., parsimonious) peptides were considered for quantification.

For the multi-consensus of Banner plus ROSMAP BA9 cases, peptide-specific TMT reporter abundance was first corrected within TMT batches using the “purityCorrect” function of the MSnbase R package prior to summing of reporter abundance of parsimonious groups of peptides. The “purity matrix” listing the fraction of specific reporter signal was assembled using TMT labeling reagent lot-specific information for the following batches (ROSMAP, batches 1-11: 10-plex kit lot RF234620; batches 12-50: channel-specific lots SG253447 (126), SG253458 (127N), SG255461 (127C), SF253450 (128N), SG253451 (128C), SH255464 (129N), SH255465 (129C), SF253465 (130N), SH253466 (130C), and SG253467 (131N)); all Banner batches used the same 10 channel-specific lots as ROSMAP batches 12-50, plus channel 11 (131C) lot #SJ258847. After correction, peptide quantitation was summed for razor plus unique peptides, thereby assembling protein abundances. Protein abundances were normalized by scaling sums of protein signal within a channel for each specific case protein sample to the maximum channel-specific protein abundance sum, as is typically calculated in the ‘normalized abundance’ columns in Proteome Discoverer output.

### Controlling for Batch-specific Variance Across Proteomics Datasets

We used a tunable median polish approach termed TAMPOR to remove technical batch variance in the proteomic data, as previously described^3^. TAMPOR is a variation on the standard median polish approach^95^ to remove intrabatch, interbatch, and intercohort technical variance while preserving meaningful biological variance in protein expression values, normalizing to the central tendency (median) of selected intrabatch or intracohort samples. This approach removes batchwise technical variance in a manner preserving other variance, and is robust to outliers and up to 50% missing values. The algorithm is fully documented and available as an R function, which can be downloaded from https://github.com/edammer/TAMPOR. If a protein had greater than 50% missing values, it was removed from the matrix. No imputation of missing values was performed for any cohort.

Prior to merging the protein abundances in the consensus ROSMAP and Banner cohorts, the multi-consensus normalized protein abundance data matrix was first segregated by cohort. Intra-cohort batch effects in the Banner cohort (22 TMT 11-plex batches) and ROSMAP cohort (50 TMT-10 plex batches) were then normalized separately in two steps, which were iterated until convergence. The first step transforms TMT reporter abundance into a ratio according to **Equation 1**, followed by log_2_ transformation. The second step scales all sample median protein log_2_ abundance ratios to zero, then unlogs the ratios and multiplies the ratios by median protein relative abundance factors recorded before step 1. The process is iterated until convergence, with output available as log_2_(normalized ratio), or unlogged normalized relative TMT reporter intensities.

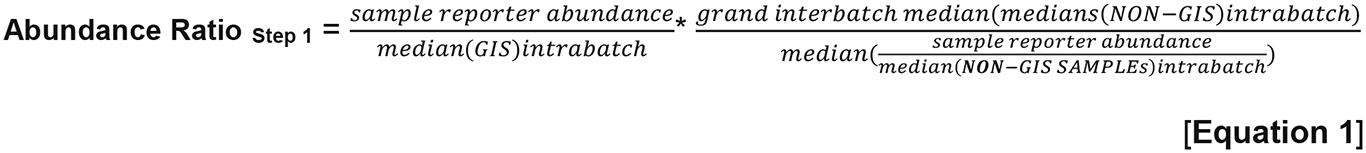

For the Banner cohort, **Equation 1** (representing TAMPOR step 1) leveraged the median protein abundance from the pooled global internal standard (GIS) TMT channels as the denominators in both of the factors to normalize sample specific protein abundances across the Banner cohort batches (**Supplementary Figure 1C**). For the ROSMAP cohort, equation 1 was employed exactly as specified, using the median of non-GIS samples balanced for diagnosis and other traits across the 50 ROSMAP TMT batches, and for the second factor numerator, the grand median of all batch-specific second-term denominators. The use of non-GIS sample intrabatch medians was needed to completely adjust for batch effect and allow GIS samples to exist as outliers in the final normalized data (**Supplementary Figure 1B**). Following removal of intra-cohort batch effects in Banner and ROSMAP cohorts separately, all samples except cohort-specific GIS samples were processed jointly with TAMPOR into a single reassembled multi-consensus sample-protein matrix using the median of within-cohort control cases as the central tendency, enforcing that the population of all log2(ratio) output for control samples within the final 598 Banner plus ROSMAP samples would tend towards zero. All other cohorts were normalized essentially as described for the Banner cohort using pooled cohort GIS channels.

### Outlier Removal and Regression of Unwanted Covariates

In the consensus ROSMAP plus Banner BA9 TMT protein abundance matrix, there were 516 of 598 individual case samples which could be classified as AD, AsymAD, or control according to our classification scheme. The other 82 cases were excluded (diagnosis labeled “Exclude” in traits) at this point. We removed outliers detected by network connectivity *Z*-transformed metric for a sample, |Z.k|>3 standard deviations from the mean Z.k, iteratively until no further detection, as previously described ^3, 4, 96^. We then ran a pioneer round of bootstrap regression (described below) before repeating the outlier check procedure. All outliers (15 before and 13 following pioneer regression) were removed from the unregressed data, and the remaining 488 case samples were regressed the same as in the pioneer round of bootstrap regression.

The consensus ROSMAP plus Banner BA9 TMT matrix, and each of the other cohorts’ protein abundance matrices, were subjected to nonparametric bootstrap regression by subtracting the trait of interest (age at death, sex, or postmortem interval (PMI)) times the median estimated coefficient from 1000 iterations of fitting for each protein in the cohort-specific log_2_(abundance) matrix. Ages at death used for regression were uncensored. Case status/diagnosis was also explicitly modeled (i.e., protected) in each regression. Following regression of each individual cohort, we assessed whether any cohort-specific tissue dissection bias was present by performing a Spearman rank correlation of traits including age, sex, PMI, and white matter markers to the top five principle components (PC) of log_2_(abundance). Any new outliers introduced by regression were not considered in the PCs. No gross difference in percent variance explained by any of the top five PCs with white matter correlation was observed.

### ROSMAP and Mt. Sinai RNA Batch Correction and Preprocessing

Regression of the RNAseq data was modeled on Sieberts et al.^97, 98^. Raw RNA counts were loaded, and variance partitioning was determined. Only genes that were expressed at a level of more than 1 count per million (CPM) total reads in at least 50% of the samples were retained for analysis. Genes were further filtered to include those with available gene length and percentage GC content from the BioMart December 2016 archive. This left 15582 genes and 633 samples after filtering. Samples with no RIN, PMI, sex or age at time of death were removed (*n*=2, leaving 631 total samples). Using our diagnostic criteria, cases were again filtered to include only those in the AD (*n*=203), AsymAD (*n*=205), and control (*n*=125) categories (total *n*=533).

The raw counts were normalized in two steps. First, to account for variations in percent GC and gene length, conditional quantile normalization (CQN) was used^99^. Secondly, a weighted linear model was applied to the raw CPM counts using the voom-limma package in Bioconductor to estimate the confidence of sampling abundance^99,100^.

Before normalization with the voom-limma package, sample outliers were detected using principal component analysis (PCA) and the aberrant distribution of the log(CPM)^101, 102^. Based on the expression pattern and the first two prinicipal components, one sample was determined to be an outlier and removed from the data. No genes were determined to be outliers. Genes that were above and below 3 standard deviations of the aberrant distribution of the log(CPM) counts were assigned NA values. The final raw counts matrix before voom-limma normalization was *n*=15582 genes by *n*=532 samples.

Using PCA analysis, the significant covariates in the data were determined (FDR<0.1). Due to the correlated nature of the covariates it is advantageous to normalize and adjust the expression matrix using an iterative approach. This was accomplished using the voom-limma package. The primary variable of interest (diagnosis) was excluded from the pool of available covariates for selection, thereby protecting it from normalization. In each round of iteration, the residual covariates were determined from the PCA analysis and were used to construct a design matrix. Voom weights were estimated for dispersion control. A linear model was then fit to the CQN expression using the voom weights and design matrix. Using the new matrix, the PCs of the residual gene expression and a new set of significant covariates were determined. If any significant residual covariates remained with FDR<0.1, the normalization was repeated.

### Differential Expression Analysis

Differentially expressed proteins were identified using one-way ANOVA followed by Holm post-hoc correction of all pairwise comparisons. Significantly altered proteins with corresponding adjusted *p* value are provided in **Supplementary Table 2**. Differential expression is presented as volcano plots, which were generated with the ggplot2 package in R v3.5.2 or the matplotlib package v3.3.2 in Python v3.8.5.

### Weighted Correlation Network Analysis (WGCNA)

We used the WeiGhted Correlation Network Analysis (WGCNA) algorithm for our network analysis pipeline, as previously described^3, 4, 49^. A weighted protein co-expression network for the Banner plus ROSMAP BA9 consensus zero-centered log2(ratio) data was generated using the *n*=8,826 log_2_ protein abundance x *n*=488 case-sample matrix that had undergone reporter purity, batch effect, and other covariate correction, as well as network connectivity outlier (*n*=28) removal as described above. Soft threshold power was determined for the data set as a data set-specific scale free topology power based on the following two guidelines: 1) The power in a plot of power (x) vs R² (y) should be where the R² has approached an asymptote, usually near or above 0.80, and 2) the mean and median connectivity at that power should not be exceedingly high, preferably around 100 or less. The power at which these criteria are met is a tradeoff between cleaning up spurious correlations due to chance (particularly important when total samples in the network are low), and maintaining sensitivity of the clustering to still be able to pick up correlations in as much of the data as possible.

An initial network was built as described below with power=7.0. Upon so doing, a single module of *n*=64 proteins was found to harbor proteins assembled from mis-cleaved tryptic peptides with higher variance in the Banner cohort driving module membership. To remove this data artifact, the clean abundance matrix values for the 64 proteins specific to measurement in Banner case-samples were set to missing values, and then enforcement of the 50% missing value threshold resulted in final input for the consensus network of *n*=8,619 proteins across *n*=488 case-samples. We confirmed that the 57 surviving proteins from the aberrant module were dispersed into diverse modules in the final network, indicating resolution of the data artifact due to this minor differential protein digestion in the Banner cohort.

The WGCNA::blockwiseModules() function was used with the following settings for the consensus network: soft threshold power=7.0, deepSplit=4, minimum module size of 20, merge cut height of 0.07, mean TOM denominator, a signed network with partitioning about medioids (PAM) respecting the dendrogram, and a reassignment threshold of *p*<0.05, with clustering completed within a single block. Specifically, this approach calculates pair-wise biweight mid-correlations (bicor, a robust correlation metric) between each protein pair, and transforms this correlation matrix into a signed adjacency matrix. The connection strength of components within this matrix is used to calculate a topological overlap matrix (TOM), which represents measurements of protein expression pattern similarity across cohort samples constructed on the pairwise correlations for all proteins within the network. Hierarchical protein correlation clustering analysis by this approach was conducted using 1-TOM, and initial module identifications were established using dynamic tree cutting as implemented in the WGCNA::blockwiseModules() function. Module eigenproteins were defined, which are the most representative abundance value for a module equivalent to the module’s first principle component, and which explain covariance of all proteins within each module^103^. Using the signedKME function in WGCNA, a table of bicor correlations between each protein and each module eigenprotein was obtained; this module membership measure is defined as k_ME_. After blockwiseModules network construction, 44 modules consisting of 18 or more proteins were detected. To enforce a kME table with no aberrant assignments to modules, a *post hoc* cleanup procedure was applied in which proteins with an intramodular kME less than 0.30 were removed, then reassignment of (a) any grey proteins (unassigned to a module) with a maximum kME to any module of >0.30 and (b) proteins with intramodular kME more than 0.10 below the maximum kME of the protein’s correlation to any other module, was done to reassign each such protein to the module corresponding to the protein’s maximum kME. Then, MEs and the signed kME table were recalculated with the WGCNA::moduleEigengenes() and WGCNA::signedKME() functions, respectively. Finally, the kME table individual protein reassignment process was repeated if additional corrections could be made, up to a total of 30 iterations. For the consensus network, this required 11 iterations until resolution, which increased the module size of the smallest module (M44) in the network to 28, and decreased grey (unassigned) protein count for the network from 3,156 (35.8%) to 2,282 (25.9%).

The WGCNA::blockwiseModules() fucntion was also used to generate the Mount Sinai RNA network, ROSMAP RNA network, and the ROSMAP RNA-protein overlap networks. The parameters used to build these networks were the same as those used in the consensus network build with the exception of the soft threshold power, which was 10.0, 12.5, 10.0, and 8.0 for the Mount Sinai RNA network, ROSMAP RNA network, ROSMAP RNA overlap network, and ROSMAP protein overlap network, respectively. As in the consensus network, a post-hoc kME table clean-up was applied to each network. The Mount Sinai RNA network contained 93 modules with minimum module size of 45 genes. The ROSMAP RNA networks were similar in size, with 88 modules and minimum module size 49 for the *n*=532 network, and 91 modules with minimum module size 44 for the *n*=168 network. The ROSMAP RNA-protein overlap network contained 69 modules with minimum module size 13.

### MONET M1 Analysis

The three top-performing methods from the DMI DREAM Challenge were compiled in the MONET toolbox and released to the public for use (https://github.com/BergmannLab/MONET.git)23. We selected the M1 method from this toolbox as a complimentary network analysis method to explore the AD TMT network. Unlike WGNCA’s hierarchical clustering method, the M1 method determines modules and communities by optimizing the well-known modularity function from Newman and Girvan^104^. However, unlike traditional modularity optimization methods, it searches the network at multiple topological scales resulting in a multiresolution approach. The authors have added the resistance parameter, *r*, which averts genes from joining modules. If *r* = 0 the method returns to Newman and Girvan’s original modularity optimization; *r* > 0 produces smaller modules (or reveals network substructure); and *r* < 0 produces larger modules (or results in network superstructure)^105^. Instead of manually choosing the parameter *r*, users are allowed to optimize their network by tuning four hyperparameters: minimum module size, maximum module size, desired average degree, and desired average degree tolerance. The MONET M1 algorithm will then fit a resistance value to the data to produce a network described by the user’s parameters. Input for M1 was an edge list, obtained from a cleaned abundance matrix as follows: power for scale-free toplogy was determined as described in the above WGCNA methods section for each M1 input network, and the adjacency was calculated for the clean abundance data matrix raised to this power using the WGCNA adjacency function with additional parameters type=“signed”, corFnc=“bicor”, and the corOptions parameter set to use pairwise complete correlation. As M1 takes an edge list as input, the adjacency upper triangle correlation values were used to populate the weights of unique pairwise correlations in the edge list. No sparsification of the edge list was applied prior to running M1 and neither TOM nor 1-TOM (dissimilarity) were considered.

We optimized the hyperparameters using a grid search by varying minimum module size, *i* ∈ {3, 10, 15, 20}, maximum module size, *j* ∈ {100, 200, 300, 400, 500}, and desired average degree, *k* ∈ {25, 50, 75}. The desired average degree tolerance was left at the default value of 0.2. Here, the optimal model was defined as the set of parameters that minimized the percentage of proteins not assigned to a module. The final parameters selected were *i* = 3, *j* = 100, *k* = 75, which built a network with 373 modules and 26.91% proteins not assigned to a module. After the network was built, the smallest modules were pruned from the graph so that the smallest module contained no less than 20 members in concordance with the WGCNA network. This increased the percent of proteins not assigned to a module to 30.22% and decreased the number of modules to 87. This final network was used in module preservation studies with the network built using the WGCNA algorithm.

### Network Preservation

We used the WGCNA::modulePreservation() function to assess network module preservation across cohorts. We also used this function to assess the effect of missing values on the consensus network. *Z*_summary_ composite preservation scores were obtained using the consensus network as the template versus each other cohort or missing value threshold tested, with 500 permutations. Random seed was set to 1 for reproducibility, and the quickCor option was set to 0. We also assessed network module preservation using synthetic eigenproteins. Briefly, protein module members in the consensus network template with a k_ME.intramodule_ among the top 20th percentile were assembled into a synthetic module in each target cohort, and synthetic modules with at least 4 members were used to calculate synthetic weighted eigengenes representing the variance of all members in the target network across case samples via the WGCNA::moduleEigengenes() function. Statistics and correlation scatterplots involving target cohort traits were then calculated and visualized.

### Network Module Overlap and Percent Novelty Analyses

Module membership by gene symbol was overlapped for all pairwise comparisons of modules in the current TMT consensus network (44 modules, this study) to those of the LFQ consensus network (13 modules) previously published^3^. A one-tailed Fisher exact test looking for significant overrepresentation or overlap was employed, and *p* values were corrected for multiple testing using the Benjamini-Hochberg method. In addition, novel gene products in the TMT network were identified and checked for significant overrepresentation (one-tailed) in the TMT consensus 44 modules not including grey, and in a second analysis, only considering the top 20 percent of gene product members of modules as ranked by kME_intramodule_. Finally, one-tailed Fisher exact tests were also employed to determine module-wise overrepresentation of amyloid plaque-^36^ and neurofibrillary tangle-^37^ associated proteins identified in previous studies. R functions fisher.test() and p.adjust() were used to obtain the above statistics.

### Gene Ontology and Cell Type Marker Enrichment Analyses

To characterize differentially expressed proteins and co-expressed proteins based on gene ontology annotation, we used GO Elite v1.2.5 as previously published^4, 86^, with pruned output visualized using an in-house R script. Cell type enrichment was also investigated as previously published^4, 86^. For the cell type enrichment analyses, we generated an in-house marker list combining previously published cell type marker lists from Sharma et al. and Zhang et al.^106, 107^ (**Supplementary Table 7**). For each of the five cell types of interest (endothelia, microglia, astrocyte, neuron, and oligodendrocyte), genes from the Sharma *et al*. list and genes from the Zhang *et al*. list were joined into one list per cell type. If, after the lists were merged, a gene symbol was assigned to two cell types, we defaulted to the cell type defined by the Zhang et al. list such that each gene symbol was affiliated with only one cell type. The gene symbols were then processed through MyGene to update them to the most current nomenclature, and converted to human symbols using homology look up. Fisher’s Exact tests were performed using the human cell type marker lists to determine cell type enrichment.

### GWAS Module Association

To determine if any protein products of GWAS targets were enriched in a particular module, we used the single nucleotide polymorphism (SNP) summary statistics from Kunkle *et al*. ^108^ to calculate the gene level association value using MAGMA v1.08b^45^, as previously described^4^. To remove SNPs in linkage disequilibrium with the *APOE* locus from consideration in the analysis, we excluded SNPs within a 2 megabase window centered on *APOE. APOE* was manually added to the gene list and assigned a –log *p* value of 50, given its known strong association with AD. SNPs associated with non-protein coding genes based on information in the current version of Ensembl bioMart were also removed from consideration (*n*=1151). A total of 31 genes with MAGMA P_MULTI<0.05 were excluded from the analysis. A final list of 1822 genes with gene-based GWAS *p*<0.05, including *APOE*, was used for enrichment analysis. Similar analyses were performed with GWAS candidates for Schizophrenia (SCZ) and Autism Spectrum Disorders (ASD). These GWAS datasets were provided and downloaded from the Psychiatric Genomics Consortium (http://www.med.unc.edu/pgc/downloads).

### PWAS Module Association

Proteins (*N*=8,356) tested in the PWAS study by Yu *et al*. ^46^ for correlation to cognitive resilience (or decline, when negatively correlated) were split into lists of unique gene symbols representing protein gene products positively correlated (*n*=645) and negatively correlated (*n*=575) to cognitive resilience, and then these lists with corresponding *p* values were separately checked for enrichment in consensus TMT network modules using a permutation-based test (10,000 permutations) implemented in R with exact *p* values for the permutation tests calculated using the permp function of the statmod package. Module-specific mean *p* values for risk enrichment were determined as a *Z* score, specifically as the difference in mean *p* value of gene product proteins hitting a module at the level of gene symbol minus the mean *p* value of genes hit in the 10,000 random replacement permutations, divided by the standard deviation of *p* value means also determined in the random permutations. This method is identical to that used for determining module-wise enrichment of risk in GWAS results summarized as gene-level *p* values using MAGMA (see GWAS Module Association methods section).

### Network Module Quantitative Trait Loci Analysis

DNA from ROS/MAP participants underwent whole genome sequencing (WGS) or genome-wide genotyping using either the Affymetrix GeneChip 6.0 or Illumina OmniQuad Express chip as previously described^109^. We used WGS when multiple data sources were available. Participants from Banner were genotyped using the Affymetrix Precision Medicine Array. Quality control of WGS and array-based genotypes were performed separately using Plink as described previously^110^. Briefly, variants with Hardy Weinberg equilibrium p-value < 10^-7^, with missing genotype rate >5%, with minor allele frequency <1%, and are not single nucleotide polymorphisms (SNPs) were removed. KING was used to remove individuals estimated to be closer than second degree kinship^111^. Genotypes were imputed to the 1000 Genome Project Phase 3^112^ using the Michigan Imputation Server^113^. SNPs with imputation *R*^2^ > 0.3 were retained for analysis. Genetic variants associated with a protein co-expression module were identified using linear regression to model the first eigenprotein of the protein module as a function of genotype, adjusting for sex, cognitive diagnosis, 10 principal components, and genotyping chip. Among genetic variants with genome-wide level of significant association with a module (p <5×10^-08^), we categorized them as either proximal or distal protein module quantitative trait locus (mod-QTL). Proximal mod-QTL was defined as SNPs within 1 megabase of any of the genes in the corresponding module, otherwise, they were categorized as distal mod-QTLs. Mod-pQTLs were clumped by Plink using default parameters so that SNPs within 250kb of one another and in linkage disequilibrium (LD r^2^ > 0.5) were represented by the lead SNP (i.e., the most statistically significant SNP in the clumped locus). Association between ApoE protein levels and the rs429358 genotype was tested in a linear regression model adjusting for diagnosis, 10 population principal components, and cohort.

### Cognitive Trajectory Analysis

ROS/MAP participants underwent cognitive testing annually in the domains of episodic memory, perceptual orientation, perceptual speed, semantic memory, and working memory as described in detail previously^114^. The raw score from each of these 17 cognitive tests was converted to a Z score using the mean and standard deviation of the cohorts at the baseline visit. Then the Z scores were averaged to create a composite annual global cognitive score. The rate of cognitive change over time for each participant was represented by the random slope of a linear mixed model where the annual global cognitive score was the longitudinal outcome, follow-up year was the independent variable, adjusting for age at recruitment, sex, and years of education as previously described^109^. We used the person-specific random slope to represent the rate of change of cognitive performance over time for each subject. To examine associations between protein co-expression modules and cognitive trajectory, we performed linear regression with cognitive trajectory as the outcome and the first module eigenprotein as the predictor with or without adjusting for the 10 measured pathologies. The 10 age-related pathologies measured in ROS/MAP included amyloid-β, tangles, cerebral amyloid angiopathy, cerebral atherosclerosis, arteriolosclerosis, Lewy body, TDP-43, gross infarct, microinfarct, and hippocampal sclerosis as described in detail before^115^. Multiple testing adjustment (for multiple modules) was addressed with Benjamini-Hochberg false discovery rate (FDR)^116^.

### Immunohistochemistry

Human forebrain 8µm thick sections were deparaffinized and processed for immunohistochemical labeling with antibodies on a ThermoFisher autostainer. Primary antibody was rabbit anti-Midkine EP1143Y (Abcam). Secondary antibody was biotinylated goat anti-mouse/rabbit (Jackson Immunoresearch Labs). Sections were blocked with normal serum and incubated with primary antibody (1:1000), then exposed to secondary antibody (1:200) followed by avidin-biotin complex (Vector ABC Elite kit) and developed with diaminobenzidine (DAB). After sections were mounted and coverslipped, images were captured using an Olympus bright-field microscope and camera (OlympusBX51). For final output, images were processed using Adobe Photoshop software.

### Other Statistics

All statistical analyses were performed in R (v3.5.2). Boxplots represent the median, 25^th^, and 75^th^ percentile extremes; thus, hinges of a box represent the interquartile range of the two middle quartiles of data within a group. The farthest data points up to 1.5 times the interquartile range away from box hinges define the extent of whiskers (error bars). Correlations were performed using the biweight midcorrelation function as implemented in the WGCNA R package. Comparisons between two groups were performed by *t* test. Comparisons among three or more groups were performed with one-way ANOVA with Tukey or *post hoc* pairwise comparison of significance. Comparison of variance was performed using *F* test. *P* values were adjusted for multiple comparisons by false discovery rate (FDR) correction where indicated. Module membership graphs were generated using the network package v2.5 in Python v3.8.5 and in-house graphing scripts.

## Supplementary Figures

**Supplementary Figure 1.**
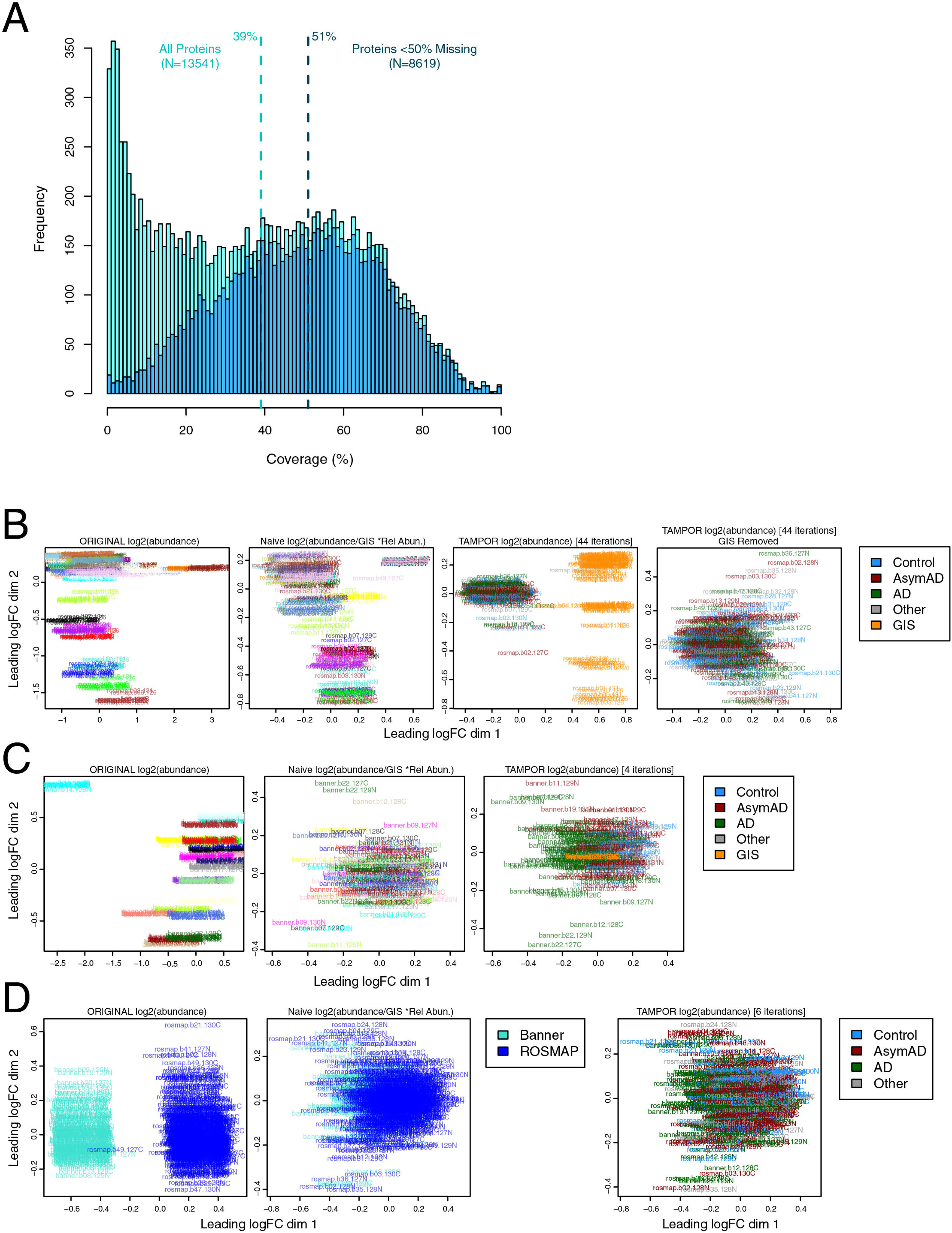
Percent Protein Coverage by TMT-MS and TMT-MS Batch Correction. (A) Percent protein coverage for all proteins measured (*n*=13,541) and those measured in at least 50% of cases (*n*=8,619), which was the threshold for inclusion in the co-expression network. The vertical dashed lines represent median percent coverage for each distribution. (B-D) TMT-MS batch correction as illustrated by multidimensional scaling (MDS). Starting log2 abundance, log2 abundance divided by the global internal standard (GIS), and additional batch correction by the TAMPOR algorithm in the ROSMAP cohort (B) and the Banner cohort (C) separately, followed by final cohort correction (D). Each batch is designated an arbitrary color in the first two panels in (B) and (C). dim, dimension; logFC, log fold change.

**Supplementary Figure 2.**
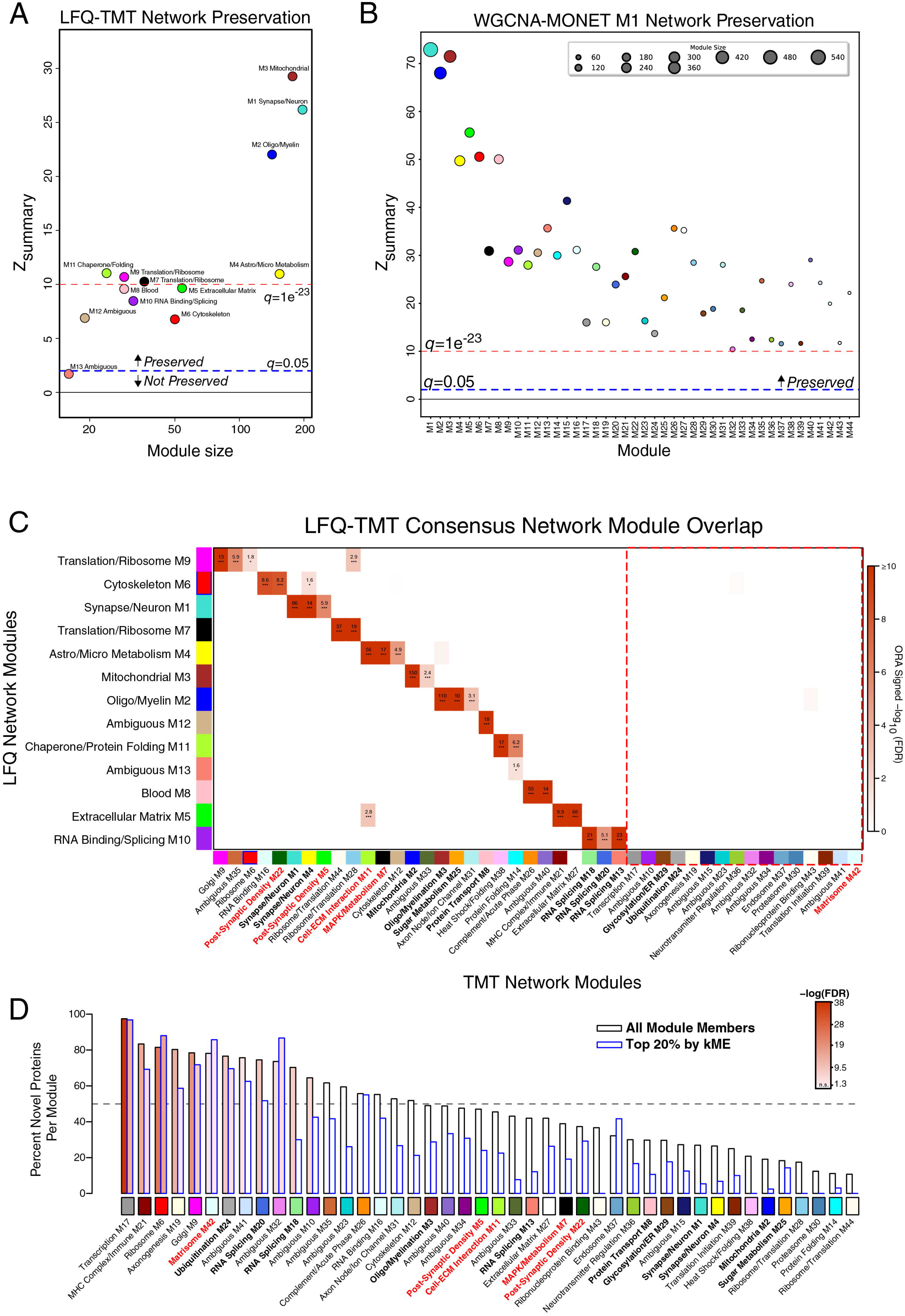
LFQ and TMT AD Network Comparison. (A-D) LFQ AD network module preservation in the TMT AD network (A). Modules that had a *Z*_summary_ score of greater than or equal to 1.96 (or *q*=0.05, blue dotted line) were considered to be preserved, while modules that had a *Z*_summary_ score greater than or equal to 10 (or *q*=1e^-23^, red dotted line) were considered to be highly preserved. (B) Preservation of the TMT AD network built using the weighted correlational network algorithm (WGCNA) into the network built on the same matrix using the MONET M1 algorithm. (C) Module member overrepresentation analysis (ORA) of the LFQ and TMT AD networks. The dashed red box highlights modules that are unique to the TMT network. The numbers in each box represent the –log_10_(FDR) value for the overlap. The heatmap is thresholded at a minimum of FDR(0.1) for clarity. (D) Percent novelty of TMT network module protein members compared to LFQ network proteins for all module members (black) or the top 20% of module proteins by strength of correlation to the module eigenprotein (kME) (blue). The dashed line indicates 50% novel protein members. Bars are shaded according to *P* value significance. ORA and percent novelty *P* values were corrected by the Benjamini-Hochberg procedure. * <0.05, ** <0.01, *** <0.005.

**Supplementary Figure 3.**
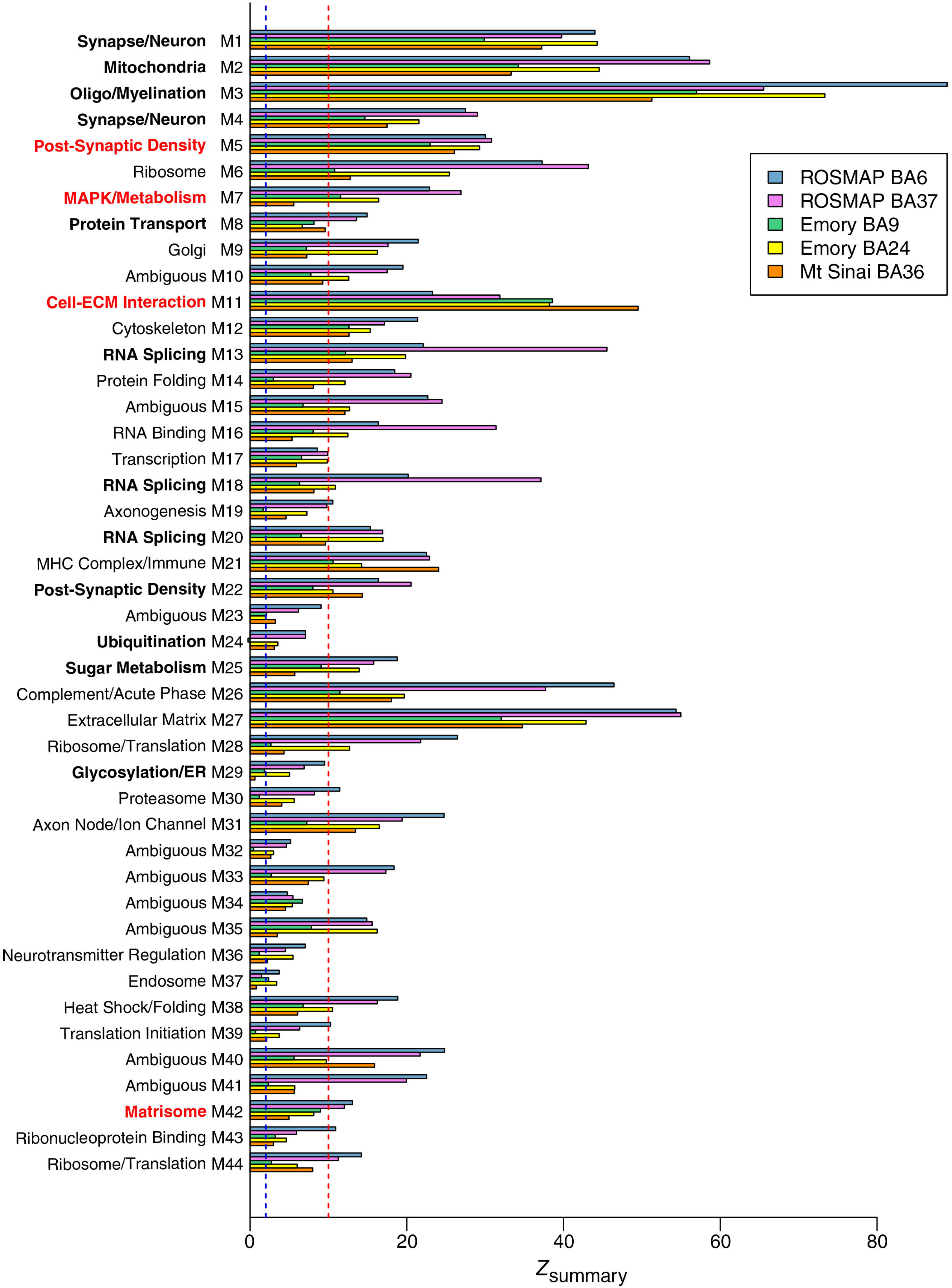
TMT AD Network Module Preservation. Modules that had a *Z*_summary_ score of greater than or equal to 1.96 (or *q*=0.05, blue dotted line) were considered to be preserved, while modules that had a *Z*_summary_ score greater than or equal to 10 (or *q*=1e^-23^, red dotted line) were considered to be highly preserved. AD, Alzheimer’s disease; Aβ, amyloid-β; AsymAD, asymptomatic Alzheimer’s disease; BA, Brodmann area; ECM, extracellular matrix; ER, endoplasmic reticulum; MAPK, mitogen-activated protein kinase.

**Supplementary Figure 4.**
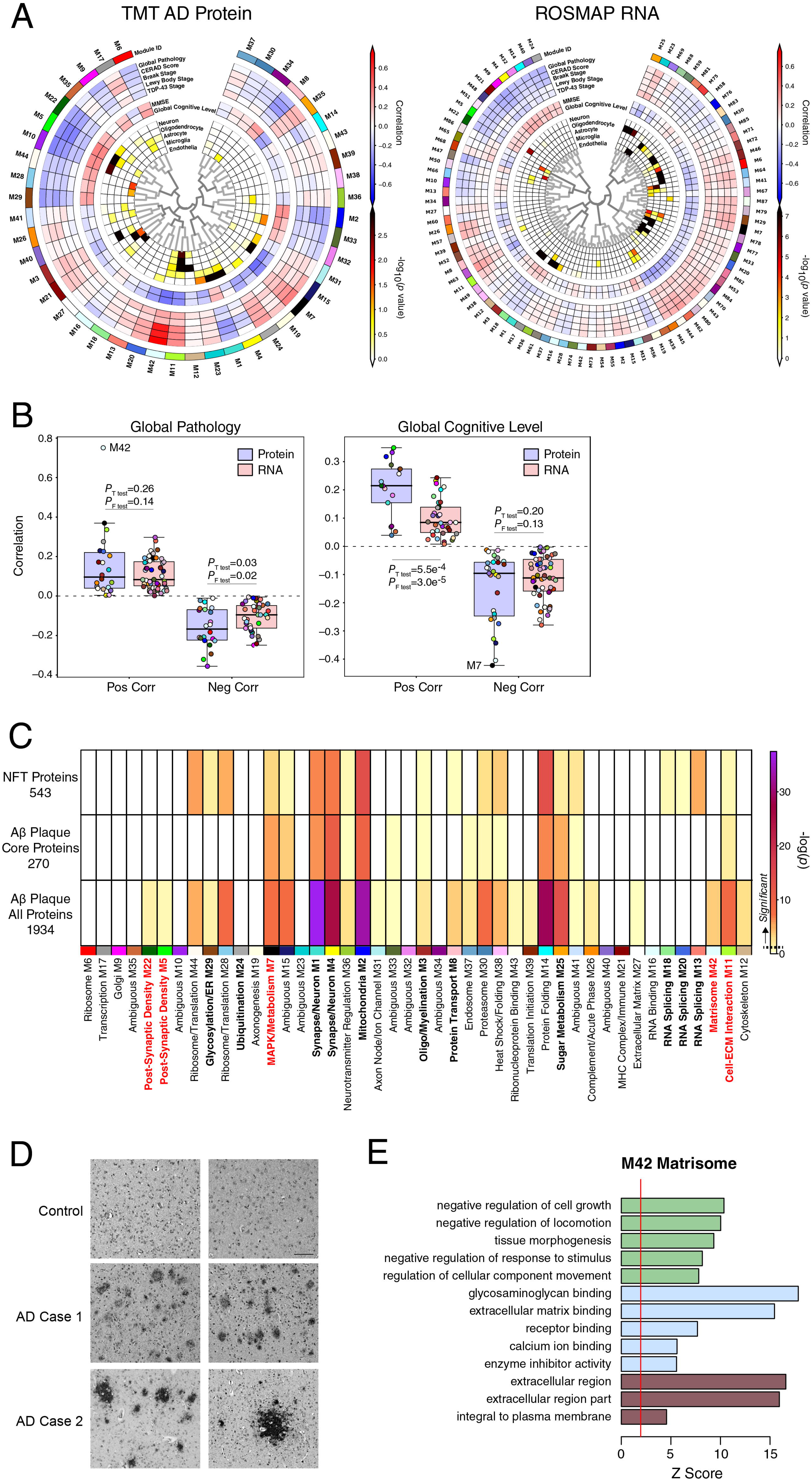
Protein and RNA AD Network Trait Correlations and TMT AD Network Module Overlap with Neurofibrillary Tangle and Aβ Plaque Proteins. (A-B) Module trait correlation analysis between protein and RNA networks. (A) WGCNA networks of TMT AD protein (left) and ROSMAP RNA (right). Protein and RNA data were obtained from the same brain region (dorsolateral prefrontal cortex, Brodmann area 9), with 168 ROSMAP cases shared between networks. Module eigenprotein to trait correlations are shown by red and blue heatmap. (B) Protein and RNA network module correlations with global pathology (left) and global cognitive level (right) as measured in ROSMAP. Differences in overall positive and negative correlations between protein and RNA modules were assessed by Welch’s *t* test, whereas differences in overall variation in correlation were measured by *F* test. *P* values for each test are provided. Boxplots represent the median, 25^th^, and 75^th^ percentiles, and box hinges represent the interquartile range of the two middle quartiles within a group. Datapoints up to 1.5 times the interquartile range from box hinge define the extent of whiskers (error bars). (C) TMT AD network module protein overlap with proteins identified as co-localized with neurofibrillary tangles (NFTs, *n*=543) and amyloid-β (Aβ) plaques as described by Drummond *et al*.^36, 37^. Overlap with Aβ plaques was performed with a set of proteins consistently observed in Aβ plaques across multiple experiments (Aβ plaque core proteins, *n*=270), as well as with a set of proteins that included proteins observed only once across multiple experiments (Aβ plaque all proteins, *n*=1934). Overlap was performed with Fisher’s exact test, and corrected by the Benjamini-Hochberg procedure. (D) Immunohistochemistry of midkine (MDK), a hub protein of the M42 matrisome module, in control and AD brain. Scale bar represents 500 µM. (E) Gene ontology analysis of the M42 matrisome module, including biological process (green), molecular function (blue), and cellular component (brown) ontologies. The red line indicates a *z* score of 1.96, or *p*=0.05.

**Supplementary Figure 5.**
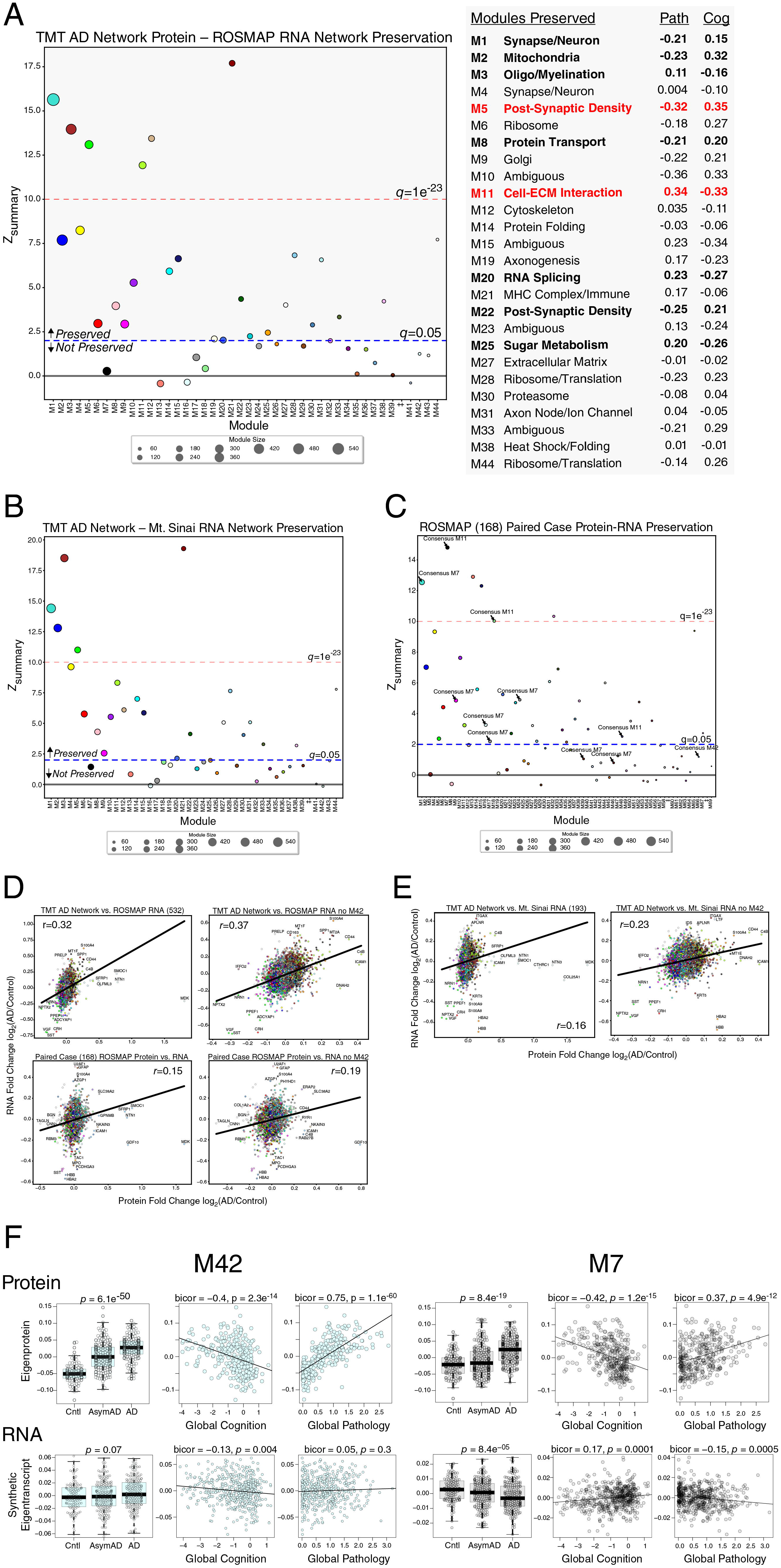
TMT AD Network Module Preservation in RNA Networks. (A-F) Module preservation of the TMT AD protein network into the ROSMAP RNA network (A). Modules that had a preservation *Z*_summary_ score less than 1.96 (*q*>0.05) were not considered to be preserved. Modules that had a *Z*_summary_ score of greater than or equal to 1.96 (or *q*=0.05, blue dotted line) were considered to be preserved, while modules that had a *Z*_summary_ score greater than or equal to 10 (or *q*=1e^-23^, red dotted line) were considered to be highly preserved. TMT AD network modules that were preserved in the RNA network, along with their correlation to global pathology and global cognition traits in ROSMAP, are listed on the right. (B) Module preservation of the TMT AD protein network into the Mt. Sinai RNA network. (C) Module preservation of the ROSMAP 168 protein network into the paired case RNA network. AD protein network module assignments are provided for M7, M11, and M42. Additional module assignments are provided in **Supplementary Table 21**. (D) Correlation of AD *versus* control RNA and protein levels between the TMT protein and ROSMAP RNA (*n*=532 cases) networks (top), as well as between cases paired between protein and RNA in ROSMAP (*n*=168), including (left) or excluding (right) M42 proteins. (E) Correlation of AD *versus* control RNA and protein levels between the TMT protein and Mt. Sinai RNA (*n*=193 cases) networks, including (left) or excluding (right) M42 proteins. Correlations were performed using Pearson correlation. (F) Comparison of M42 matrisome (left) and M7 MAPK/metabolism (right) eigenprotein (top) and synthetic eigentranscript (bottom, *n*=532 ROSMAP cases) levels by case status, and correlation with global pathology and global cognitive function in ROSMAP. Differences were assessed by one-way ANOVA. Correlations were performed using bicor. Boxplots represent the median, 25^th^, and 75^th^ percentiles, and box hinges represent the interquartile range of the two middle quartiles within a group. Datapoints up to 1.5 times the interquartile range from box hinge define the extent of whiskers (error bars).

**Supplementary Figure 6.**
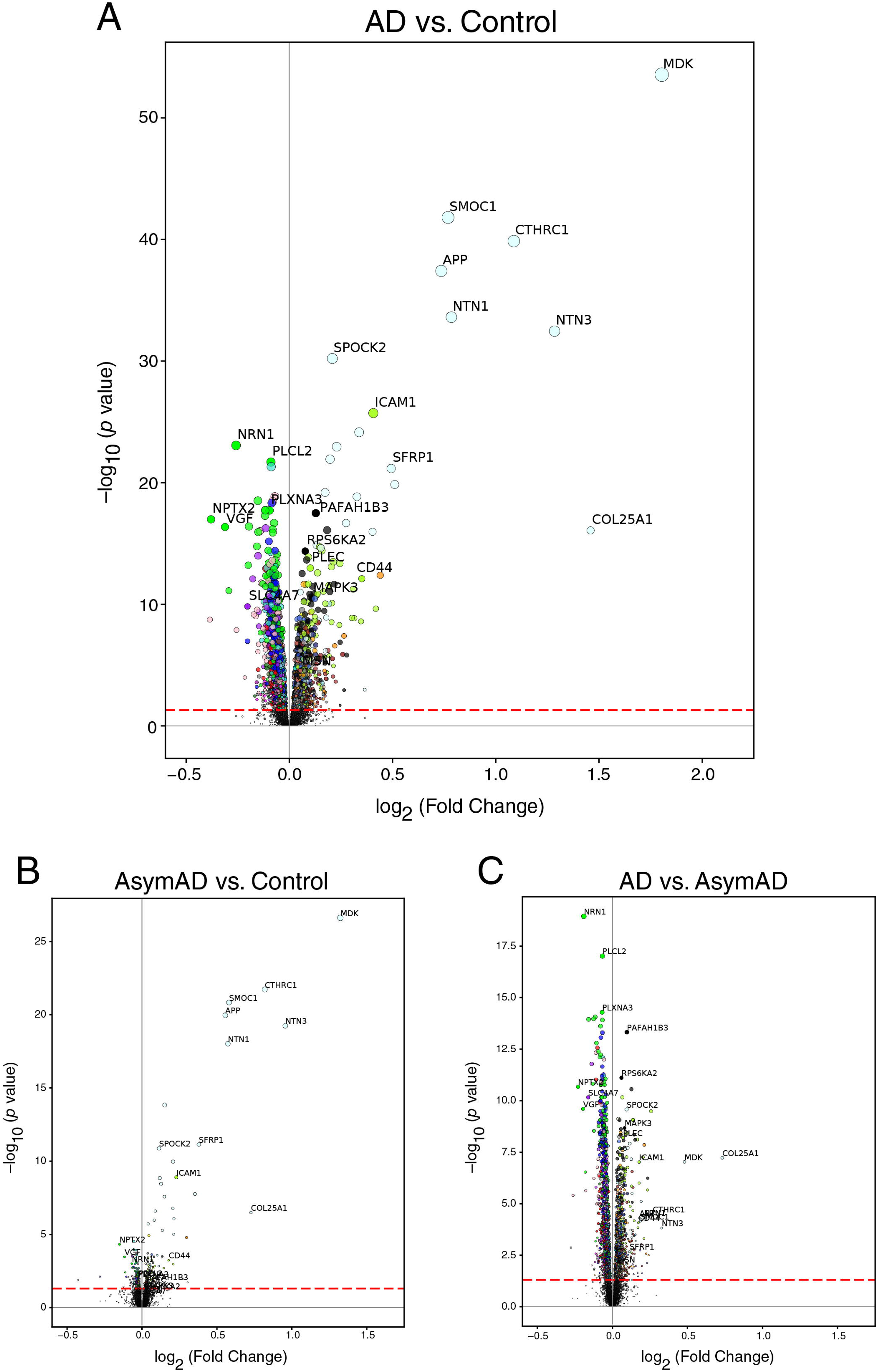
TMT AD Network Protein Differential Expression. (A-C) Differential expression between AD *versus* control (A), AsymAD *versus* control (B), and AsymAD *versus* AD (C). The dashed red line indicates a *P* value of 0.05, above which proteins are considered significantly differentially expressed. Proteins are colored by the network module in which they reside, according to the module color scheme provided in **Figure 1B**. Proteins in the M42 matrisome module are colored lightcyan. Significance was adjusted by the Holm procedure. Fold change and statistics for all proteins are provided in **Supplementary Table 2.**

## Extended Data

**Extended Data 1. GO Analysis on TMT AD Network Modules.** Gene ontology (GO) analysis was performed to gain insight into the biological meaning of each AD protein network module. Enrichment for a given ontology is shown by *z* score.

**Extended Data 2. TMT AD Network Module Eigenprotein Levels.** Differences among case groups were assessed by one-way ANOVA with Tukey test. Boxplots represent the median, 25^th^, and 75^th^ percentiles, and box hinges represent the interquartile range of the two middle quartiles within a group. Datapoints up to 1.5 times the interquartile range from box hinge define the extent of whiskers (error bars).

**Extended Data 3. TMT AD Network Module Protein Graphs and Overlap with Amyloid-β Plaque and Tau Neurofibrillary Tangle Proteins.** The size of each circle indicates the relative eigenprotein correlation value (kME) in each network module. Those proteins with the largest kME are considered “hub” proteins within the module, and explain the largest variance in module expression. Lines represent weighted adjacency values between proteins. Proteins colored orange are consistently found associated with amyloid-β plaques. Proteins colored dark blue are found to be associated with tau neurofibrillary tangles (NFTs). Proteins colored green are found to be associated with both plaques and tangles.

