## Supplementary material for "Large-Scale Deep Multi-Layer Analysis of Alzheimer’s Disease Brain Reveals Strong Proteomic Disease-Related Changes Not Observed at the RNA Level": Johnson et al 2021 Extended Data

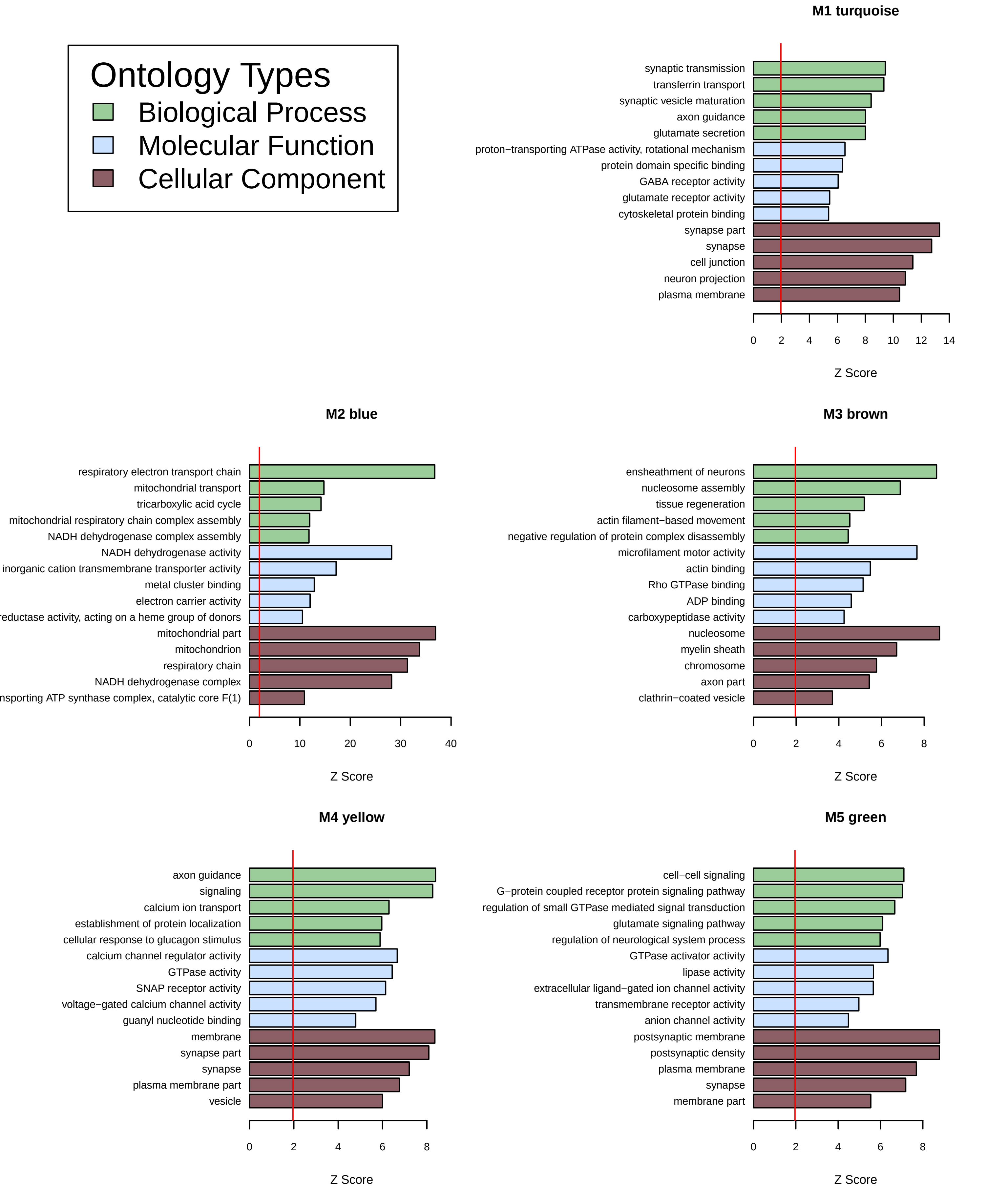
Ontology Types
Biological Process
Molecular Function
Cellular Component
M1 turquoise
synaptic transmission
transferrin transport
synaptic vesicle maturation
axon guidance
glutamate secretion
proton−transporting ATPase activity, rotational mechanism
protein domain specific binding
GABA receptor activity
glutamate receptor activity
cytoskeletal protein binding
synapse part
synapse
cell junction
neuron projection
plasma membrane
0 2 4 6 8 10 12 14
Z Score
M2 blue
respiratory electron transport chain
mitochondrial transport
tricarboxylic acid cycle
mitochondrial respiratory chain complex assembly
NADH dehydrogenase complex assembly
NADH dehydrogenase activity
inorganic cation transmembrane transporter activity
metal cluster binding
electron carrier activity
reductase activity, acting on a heme group of donors
mitochondrial part
mitochondrion
respiratory chain
NADH dehydrogenase complex
nsporting ATP synthase complex, catalytic core F(1)
0 10 20 30 40
Z Score
M3 brown
ensheathment of neurons
nucleosome assembly
tissue regeneration
actin filament−based movement
negative regulation of protein complex disassembly
microfilament motor activity
actin binding
Rho GTPase binding
ADP binding
carboxypeptidase activity
nucleosome
myelin sheath
chromosome
axon part
clathrin−coated vesicle
0 2 4 6 8
Z Score
M4 yellow
axon guidance
signaling
calcium ion transport
establishment of protein localization
cellular response to glucagon stimulus
calcium channel regulator activity
GTPase activity
SNAP receptor activity
voltage−gated calcium channel activity
guanyl nucleotide binding
membrane
synapse part
synapse
plasma membrane part
vesicle
0 2 4 6 8
Z Score
M5 green
cell−cell signaling
G−protein coupled receptor protein signaling pathway
regulation of small GTPase mediated signal transduction
glutamate signaling pathway
regulation of neurological system process
GTPase activator activity
lipase activity
extracellular ligand−gated ion channel activity
transmembrane receptor activity
anion channel activity
postsynaptic membrane
postsynaptic density
plasma membrane
synapse
membrane part
0 2 4 6 8
Z Score

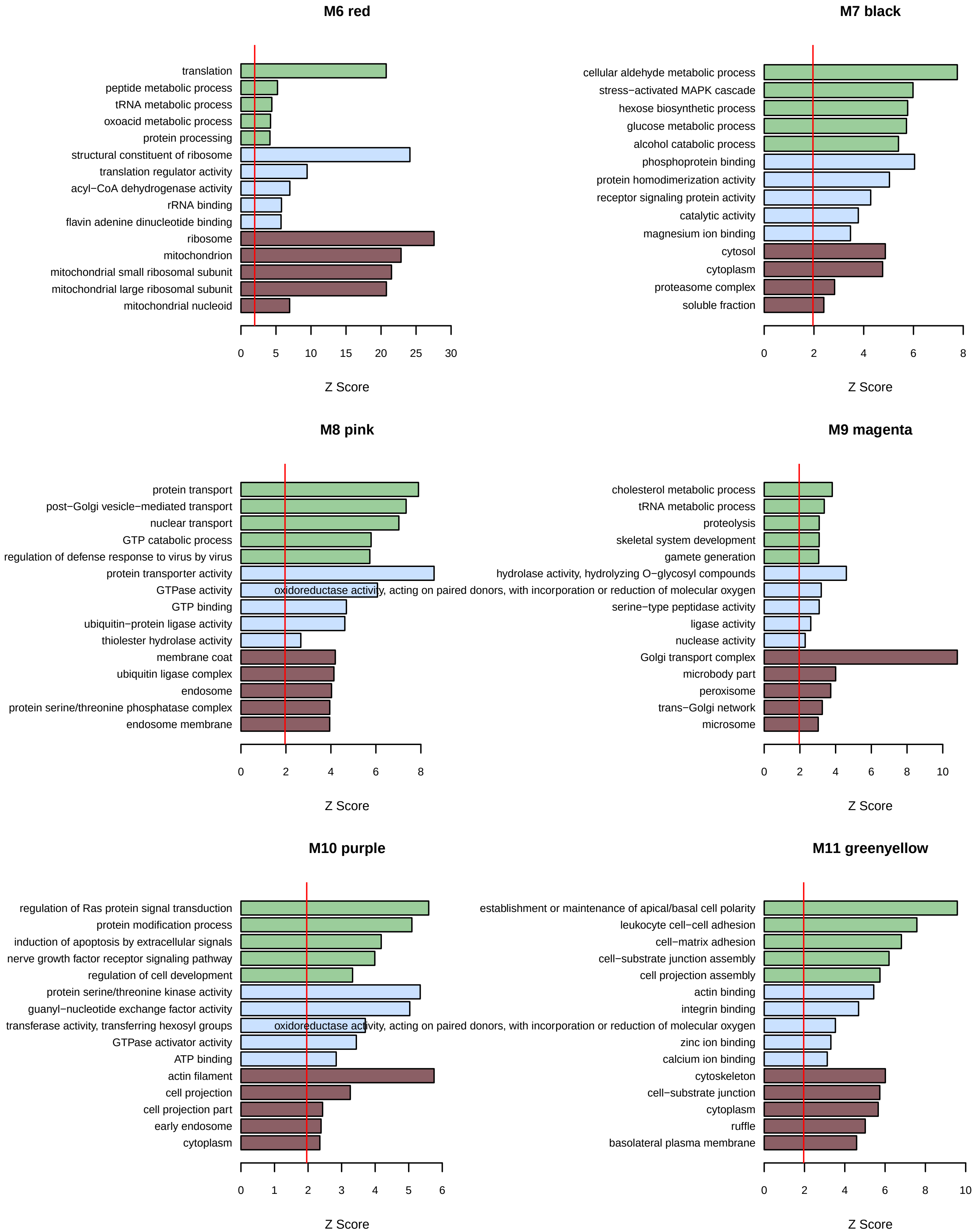
M6 red
translation
peptide metabolic process
tRNA metabolic process
oxoacid metabolic process
protein processing
structural constituent of ribosome
translation regulator activity
acyl−CoA dehydrogenase activity
rRNA binding
flavin adenine dinucleotide binding
ribosome
mitochondrion
mitochondrial small ribosomal subunit
mitochondrial large ribosomal subunit
mitochondrial nucleoid
0 5 10 15 20 25 30
Z Score
M7 black
cellular aldehyde metabolic process
stress−activated MAPK cascade
hexose biosynthetic process
glucose metabolic process
alcohol catabolic process
phosphoprotein binding
protein homodimerization activity
receptor signaling protein activity
catalytic activity
magnesium ion binding
cytosol
cytoplasm
proteasome complex
soluble fraction
0 2 4 6 8
Z Score
M8 pink
protein transport
post−Golgi vesicle−mediated transport
nuclear transport
GTP catabolic process
regulation of defense response to virus by virus
protein transporter activity
GTPase activity
GTP binding
ubiquitin−protein ligase activity
thiolester hydrolase activity
membrane coat
ubiquitin ligase complex
endosome
protein serine/threonine phosphatase complex
endosome membrane
0 2 4 6 8
Z Score
M9 magenta
cholesterol metabolic process
tRNA metabolic process
proteolysis
skeletal system development
gamete generation
hydrolase activity, hydrolyzing O−glycosyl compounds
oxidoreductase activity, acting on paired donors, with incorporation or reduction of molecular oxygen
serine−type peptidase activity
ligase activity
nuclease activity
Golgi transport complex
microbody part
peroxisome
trans−Golgi network
microsome
0 2 4 6 8 10
Z Score
M10 purple
regulation of Ras protein signal transduction
protein modification process
induction of apoptosis by extracellular signals
nerve growth factor receptor signaling pathway
regulation of cell development
protein serine/threonine kinase activity
guanyl−nucleotide exchange factor activity
transferase activity, transferring hexosyl groups
GTPase activator activity
ATP binding
actin filament
cell projection
cell projection part
early endosome
cytoplasm
0 1 2 3 4 5 6
Z Score
M11 greenyellow
establishment or maintenance of apical/basal cell polarity
leukocyte cell−cell adhesion
cell−matrix adhesion
cell−substrate junction assembly
cell projection assembly
actin binding
integrin binding
oxidoreductase activity, acting on paired donors, with incorporation or reduction of molecular oxygen
zinc ion binding
calcium ion binding
cytoskeleton
cell−substrate junction
cytoplasm
ruffle
basolateral plasma membrane
0 2 4 6 8 10
Z Score

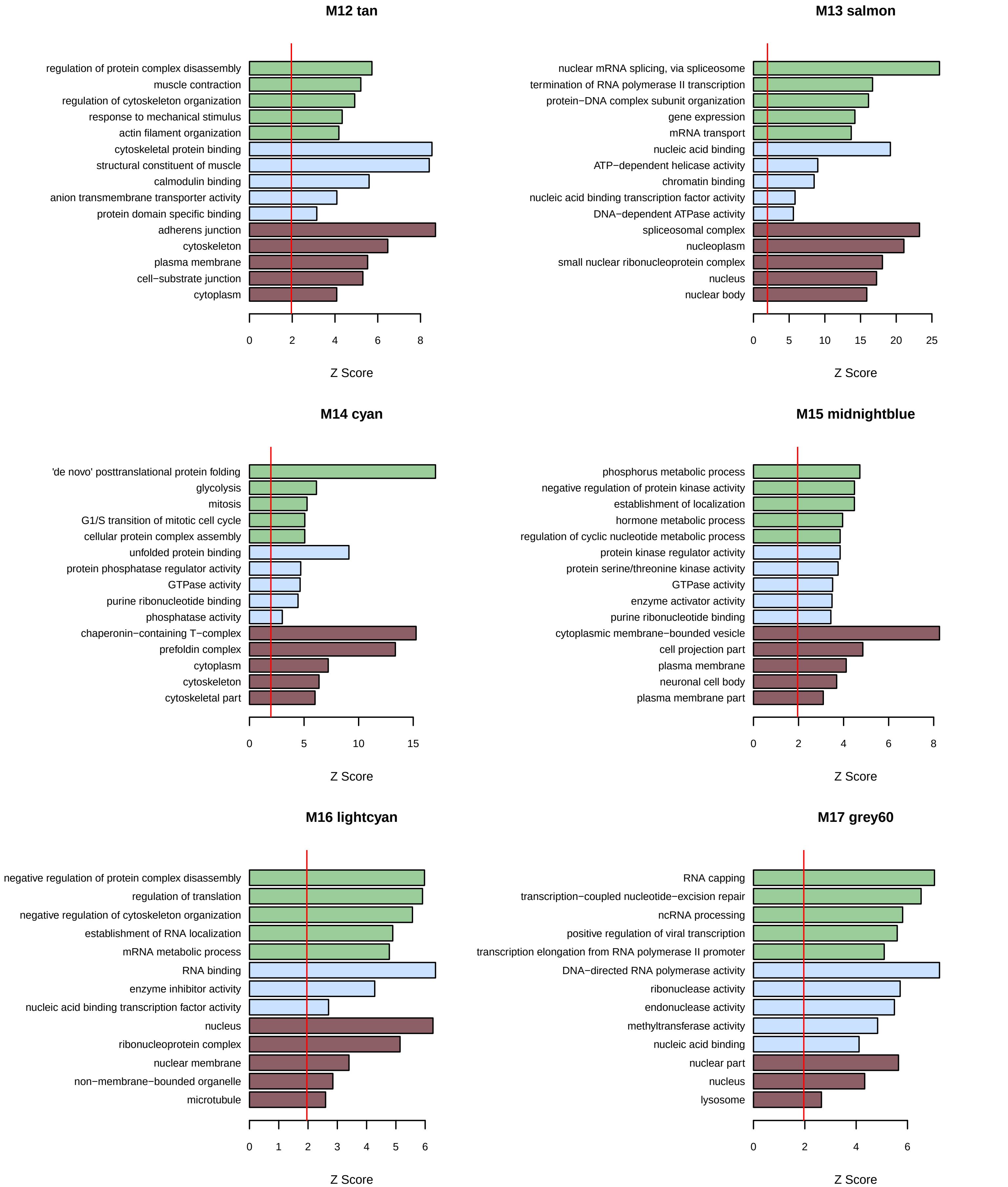

M12 tan
regulation of protein complex disassembly
muscle contraction
regulation of cytoskeleton organization
response to mechanical stimulus
actin filament organization
cytoskeletal protein binding
structural constituent of muscle
calmodulin binding
anion transmembrane transporter activity
protein domain specific binding
adherens junction
cytoskeleton
plasma membrane
cell−substrate junction
cytoplasm
0 2 4 6 8
Z Score
M13 salmon
nuclear mRNA splicing, via spliceosome
termination of RNA polymerase II transcription
protein−DNA complex subunit organization
gene expression
mRNA transport
nucleic acid binding
ATP−dependent helicase activity
chromatin binding
nucleic acid binding transcription factor activity
DNA−dependent ATPase activity
spliceosomal complex
nucleoplasm
small nuclear ribonucleoprotein complex
nucleus
nuclear body
0 5 10 15 20 25
Z Score
M14 cyan
'de novo' posttranslational protein folding
glycolysis
mitosis
G1/S transition of mitotic cell cycle
cellular protein complex assembly
unfolded protein binding
protein phosphatase regulator activity
GTPase activity
purine ribonucleotide binding
phosphatase activity
chaperonin−containing T−complex
prefoldin complex
cytoplasm
cytoskeleton
cytoskeletal part
0 5 10 15
Z Score
M15 midnightblue
phosphorus metabolic process
negative regulation of protein kinase activity
establishment of localization
hormone metabolic process
regulation of cyclic nucleotide metabolic process
protein kinase regulator activity
protein serine/threonine kinase activity
GTPase activity
enzyme activator activity
purine ribonucleotide binding
cytoplasmic membrane−bounded vesicle
cell projection part
plasma membrane
neuronal cell body
plasma membrane part
0 2 4 6 8
Z Score
M16 lightcyan
negative regulation of protein complex disassembly
regulation of translation
negative regulation of cytoskeleton organization
establishment of RNA localization
mRNA metabolic process
RNA binding
enzyme inhibitor activity
nucleic acid binding transcription factor activity
nucleus
ribonucleoprotein complex
nuclear membrane
non−membrane−bounded organelle
microtubule
0 1 2 3 4 5 6
Z Score
M17 grey60
RNA capping
transcription−coupled nucleotide−excision repair
ncRNA processing
positive regulation of viral transcription
transcription elongation from RNA polymerase II promoter
DNA−directed RNA polymerase activity
ribonuclease activity
endonuclease activity
methyltransferase activity
nucleic acid binding
nuclear part
nucleus
lysosome
0 2 4 6
Z Score

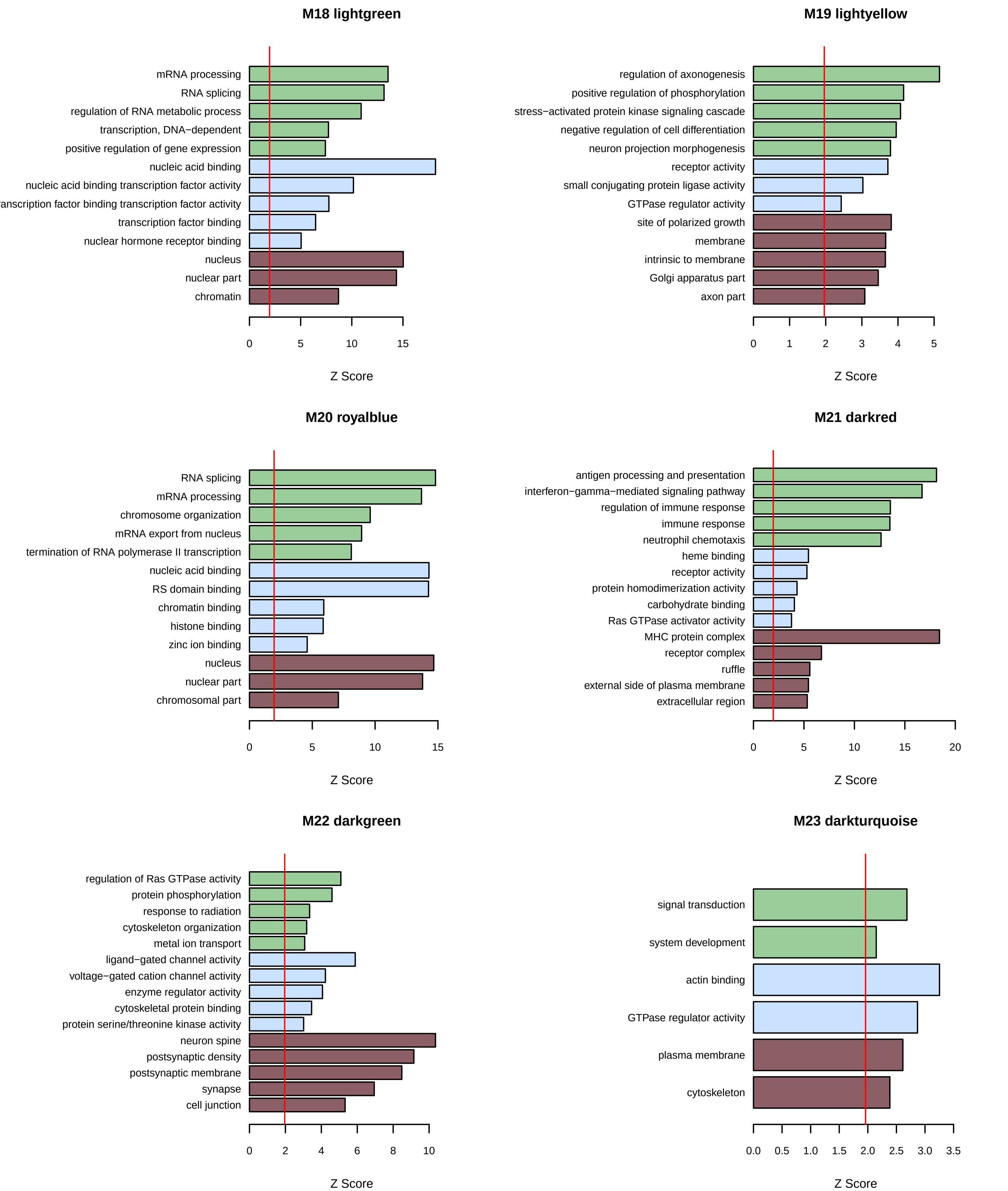

M18 lightgreen
mRNA processing
RNA splicing
regulation of RNA metabolic process
transcription, DNA−dependent
positive regulation of gene expression
nucleic acid binding
nucleic acid binding transcription factor activity
anscription factor binding transcription factor activity
transcription factor binding
nuclear hormone receptor binding
nucleus
nuclear part
chromatin
0 5 10 15
Z Score
M19 lightyellow
regulation of axonogenesis
positive regulation of phosphorylation
stress−activated protein kinase signaling cascade
negative regulation of cell differentiation
neuron projection morphogenesis
receptor activity
small conjugating protein ligase activity
GTPase regulator activity
site of polarized growth
membrane
intrinsic to membrane
Golgi apparatus part
axon part
0 1 2 3 4 5
Z Score
M20 royalblue
RNA splicing
mRNA processing
chromosome organization
mRNA export from nucleus
termination of RNA polymerase II transcription
nucleic acid binding
RS domain binding
chromatin binding
histone binding
zinc ion binding
nucleus
nuclear part
chromosomal part
0 5 10 15
Z Score
M21 darkred
antigen processing and presentation
interferon−gamma−mediated signaling pathway
regulation of immune response
immune response
neutrophil chemotaxis
heme binding
receptor activity
protein homodimerization activity
carbohydrate binding
Ras GTPase activator activity
MHC protein complex
receptor complex
ruffle
external side of plasma membrane
extracellular region
0 5 10 15 20
Z Score
M22 darkgreen
regulation of Ras GTPase activity
protein phosphorylation
response to radiation
cytoskeleton organization
metal ion transport
ligand−gated channel activity
voltage−gated cation channel activity
enzyme regulator activity
cytoskeletal protein binding
protein serine/threonine kinase activity
neuron spine
postsynaptic density
postsynaptic membrane
synapse
cell junction
0 2 4 6 8 10
Z Score
M23 darkturquoise
signal transduction
system development
actin binding
GTPase regulator activity
plasma membrane
cytoskeleton
0.0 0.5 1.0 1.5 2.0 2.5 3.0 3.5
Z Score

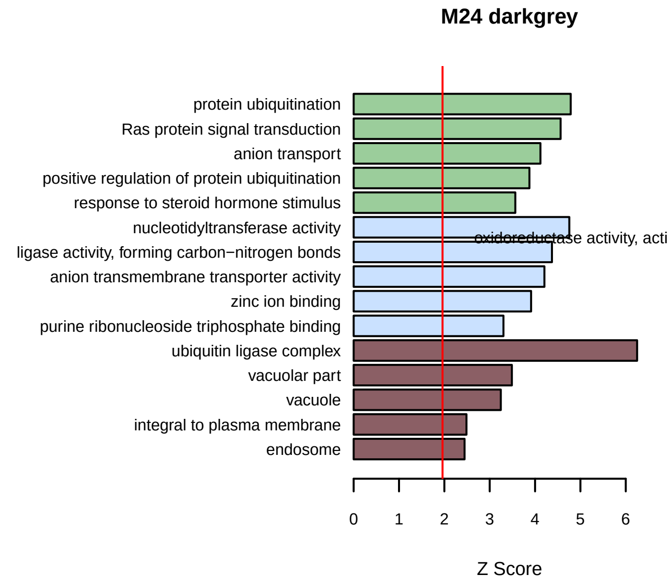
M24 darkgrey
protein ubiquitination
Ras protein signal transduction
anion transport
positive regulation of protein ubiquitination
response to steroid hormone stimulus
nucleotidyltransferase activity
ligase activity, forming carbon−nitrogen bonds
anion transmembrane transporter activity
zinc ion binding
purine ribonucleoside triphosphate binding
ubiquitin ligase complex
vacuolar part
vacuole
integral to plasma membrane
endosome
0 1 2 3 4 5 6
Z Score

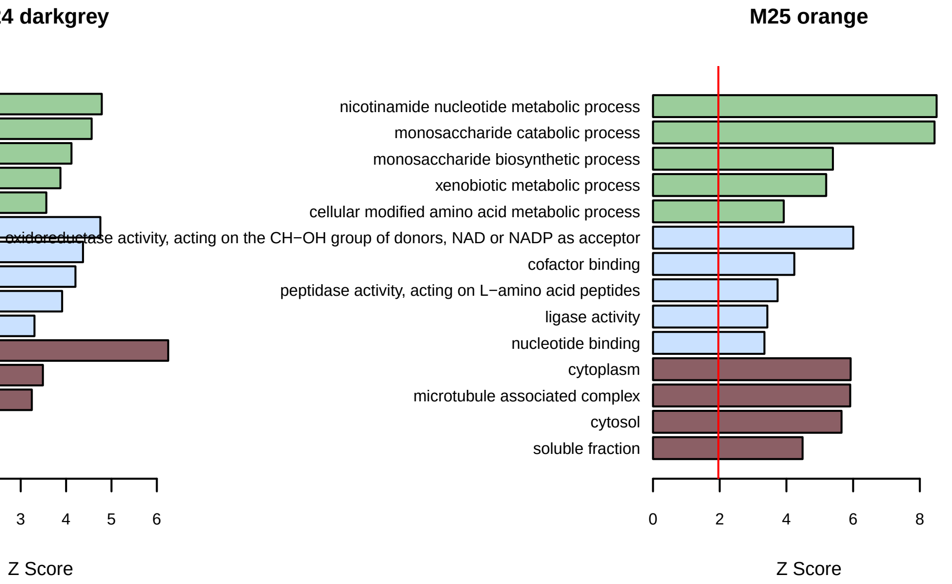
M25 orange
nicotinamide nucleotide metabolic process
monosaccharide catabolic process
monosaccharide biosynthetic process
xenobiotic metabolic process
cellular modified amino acid metabolic process
oxidoreductase activity, acting on the CH−OH group of donors, NAD or NADP as acceptor
cofactor binding
peptidase activity, acting on L−amino acid peptides
ligase activity
nucleotide binding
cytoplasm
microtubule associated complex
cytosol
soluble fraction
0 2 4 6 8
Z Score

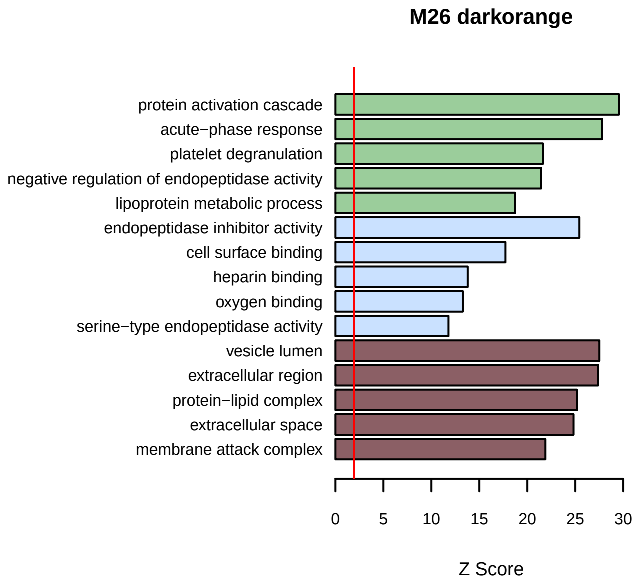
M26 darkorange
protein activation cascade
acute−phase response
platelet degranulation
negative regulation of endopeptidase activity
lipoprotein metabolic process
endopeptidase inhibitor activity
cell surface binding
heparin binding
oxygen binding
serine−type endopeptidase activity
vesicle lumen
extracellular region
protein−lipid complex
extracellular space
membrane attack complex
0 5 10 15 20 25 30
Z Score

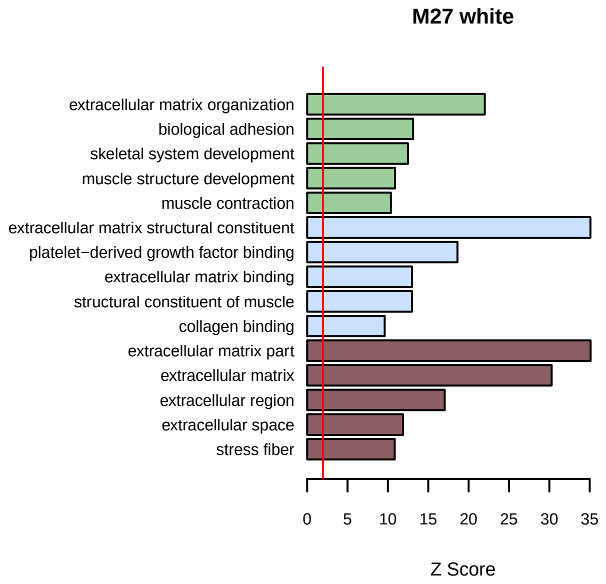
M27 white
extracellular matrix organization
biological adhesion
skeletal system development
muscle structure development
muscle contraction
extracellular matrix structural constituent
platelet−derived growth factor binding
extracellular matrix binding
structural constituent of muscle
collagen binding
extracellular matrix part
extracellular matrix
extracellular region
extracellular space
stress fiber
0 5 10 15 20 25 30 35
Z Score

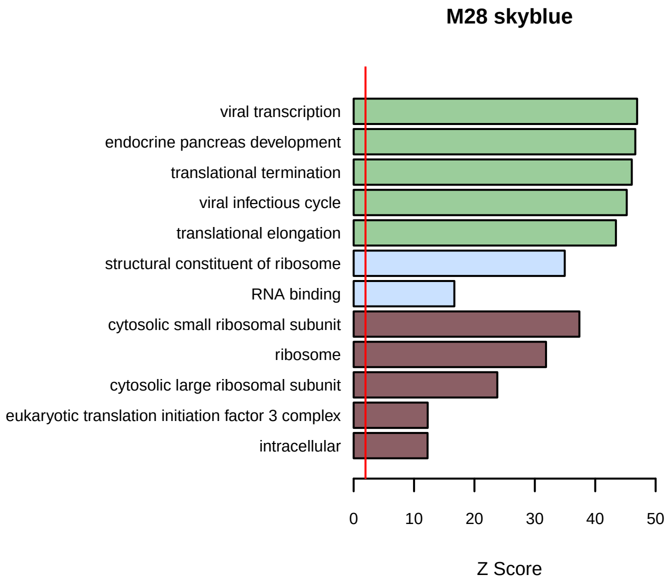
M28 skyblue
viral transcription
endocrine pancreas development
translational termination
viral infectious cycle
translational elongation
structural constituent of ribosome
RNA binding
cytosolic small ribosomal subunit
ribosome
cytosolic large ribosomal subunit
eukaryotic translation initiation factor 3 complex
intracellular
0 10 20 30 40 50
Z Score

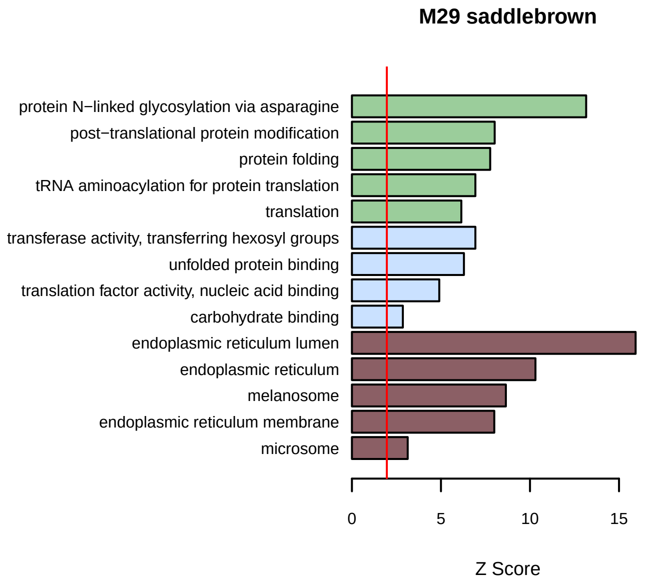
M29 saddlebrown
protein N−linked glycosylation via asparagine
post−translational protein modification
protein folding
tRNA aminoacylation for protein translation
translation
transferase activity, transferring hexosyl groups
unfolded protein binding
translation factor activity, nucleic acid binding
carbohydrate binding
endoplasmic reticulum lumen
endoplasmic reticulum
melanosome
endoplasmic reticulum membrane
microsome
0 5 10 15
Z Score

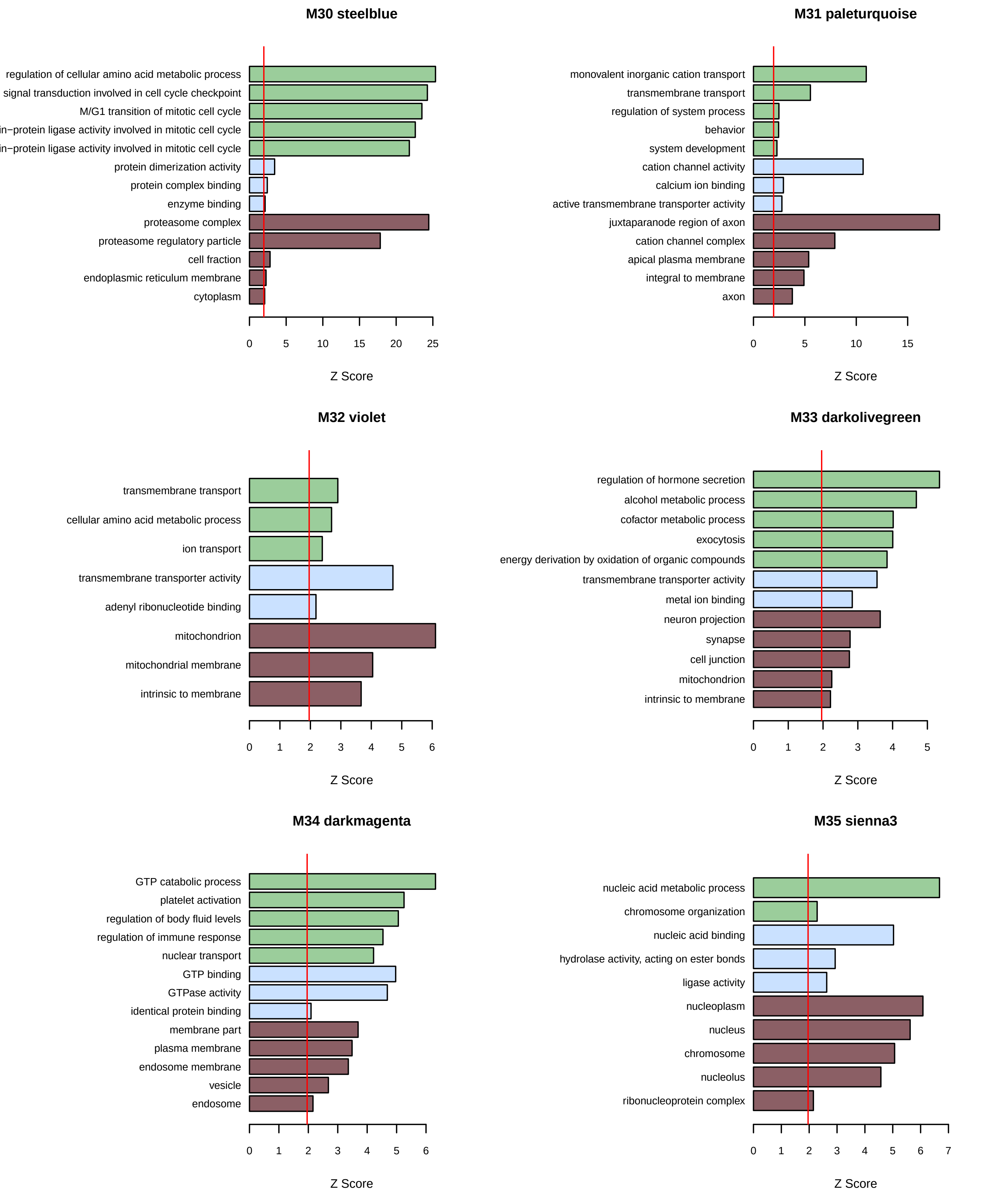

M30 steelblue
regulation of cellular amino acid metabolic process
signal transduction involved in cell cycle checkpoint
M/G1 transition of mitotic cell cycle
in−protein ligase activity involved in mitotic cell cycle
in−protein ligase activity involved in mitotic cell cycle
protein dimerization activity
protein complex binding
enzyme binding
proteasome complex
proteasome regulatory particle
cell fraction
endoplasmic reticulum membrane
cytoplasm
0 5 10 15 20 25
Z Score
M31 paleturquoise
monovalent inorganic cation transport
transmembrane transport
regulation of system process
behavior
system development
cation channel activity
calcium ion binding
active transmembrane transporter activity
juxtaparanode region of axon
cation channel complex
apical plasma membrane
integral to membrane
axon
0 5 10 15
Z Score
M32 violet
transmembrane transport
cellular amino acid metabolic process
ion transport
transmembrane transporter activity
adenyl ribonucleotide binding
mitochondrion
mitochondrial membrane
intrinsic to membrane
0 1 2 3 4 5 6
Z Score
M33 darkolivegreen
regulation of hormone secretion
alcohol metabolic process
cofactor metabolic process
exocytosis
energy derivation by oxidation of organic compounds
transmembrane transporter activity
metal ion binding
neuron projection
synapse
cell junction
mitochondrion
intrinsic to membrane
0 1 2 3 4 5
Z Score
M34 darkmagenta
GTP catabolic process
platelet activation
regulation of body fluid levels
regulation of immune response
nuclear transport
GTP binding
GTPase activity
identical protein binding
membrane part
plasma membrane
endosome membrane
vesicle
endosome
0 1 2 3 4 5 6
Z Score
M35 sienna3
nucleic acid metabolic process
chromosome organization
nucleic acid binding
hydrolase activity, acting on ester bonds
ligase activity
nucleoplasm
nucleus
chromosome
nucleolus
ribonucleoprotein complex
0 1 2 3 4 5 6 7
Z Score

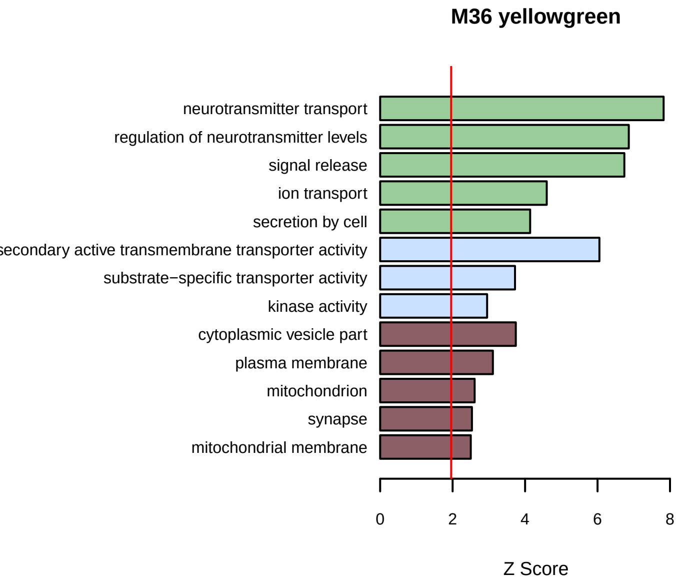
M36 yellowgreen
neurotransmitter transport
regulation of neurotransmitter levels
signal release
ion transport
secretion by cell
secondary active transmembrane transporter activity
substrate−specific transporter activity
kinase activity
cytoplasmic vesicle part
plasma membrane
mitochondrion
synapse
mitochondrial membrane
0
2
4
6
8
Z Score

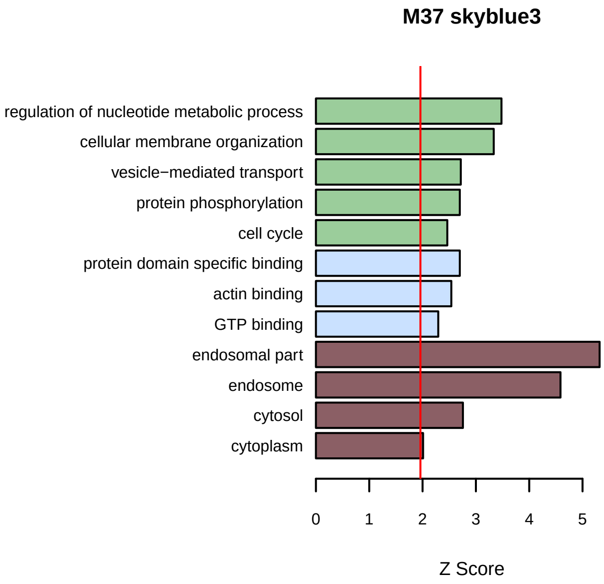
M37 skyblue3
regulation of nucleotide metabolic process
cellular membrane organization
vesicle−mediated transport
protein phosphorylation
cell cycle
protein domain specific binding
actin binding
GTP binding
endosomal part
endosome
cytosol
cytoplasm
0
1
2
3
4
5
Z Score

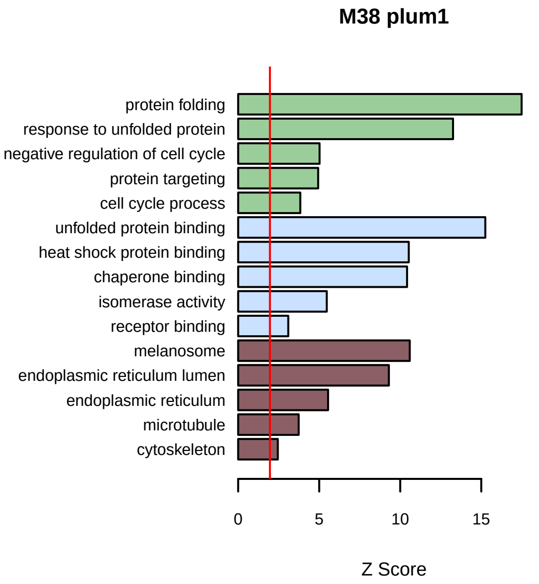
M38 plum1
protein folding
response to unfolded protein
negative regulation of cell cycle
protein targeting
cell cycle process
unfolded protein binding
heat shock protein binding
chaperone binding
isomerase activity
receptor binding
melanosome
endoplasmic reticulum lumen
endoplasmic reticulum
microtubule
cytoskeleton
0
5
10
15
Z Score

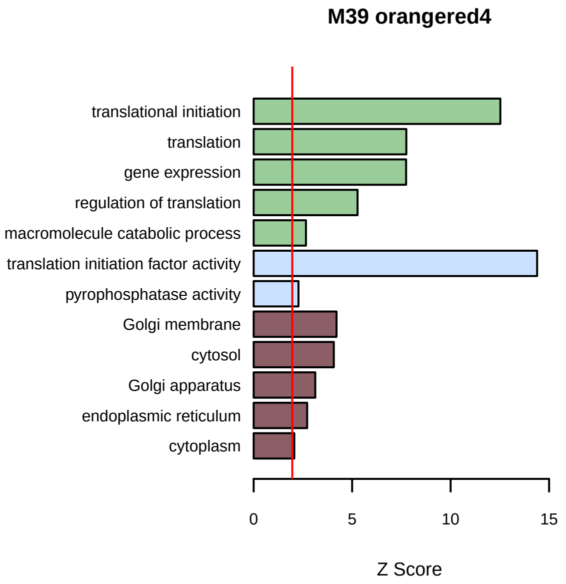
M39 orangered4
translational initiation
translation
gene expression
regulation of translation
macromolecule catabolic process
translation initiation factor activity
pyrophosphatase activity
Golgi membrane
cytosol
Golgi apparatus
endoplasmic reticulum
cytoplasm
0
5
10
15
Z Score

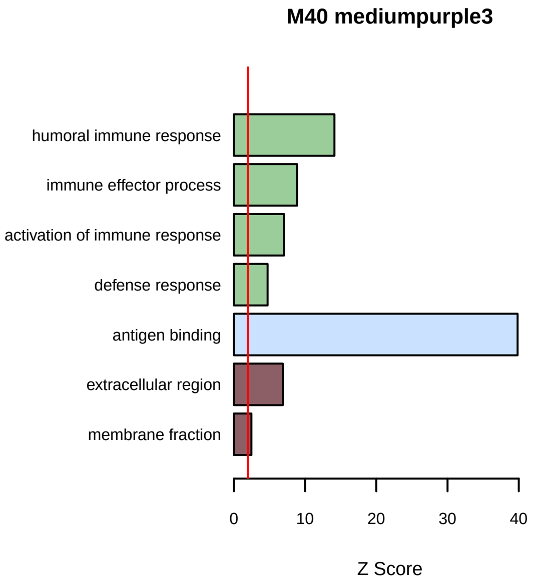
M40 mediumpurple3
humoral immune response
immune effector process
activation of immune response
defense response
antigen binding
extracellular region
membrane fraction
0
10
20
30
40
Z Score

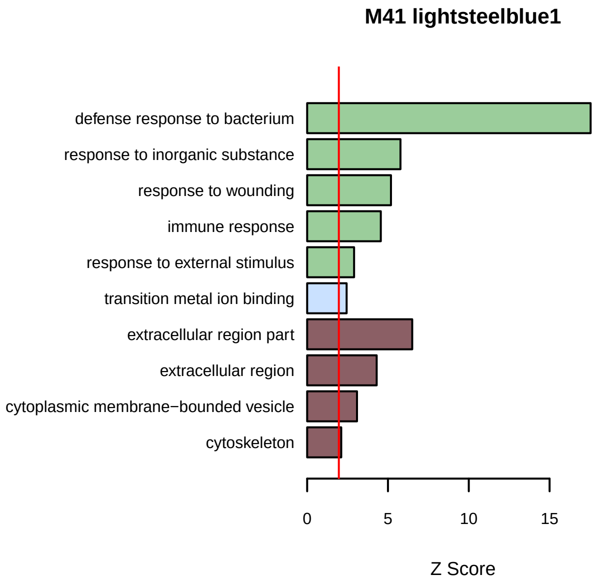
M41 lightsteelblue1
defense response to bacterium
response to inorganic substance
response to wounding
immune response
response to external stimulus
transition metal ion binding
extracellular region part
extracellular region
cytoplasmic membrane−bounded vesicle
cytoskeleton
0
5
10
15
Z Score

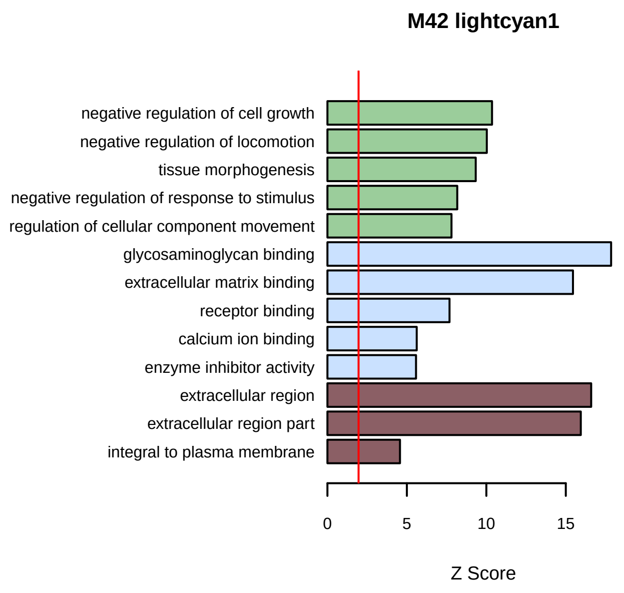
M42 lightcyan1
negative regulation of cell growth
negative regulation of locomotion
tissue morphogenesis
negative regulation of response to stimulus
regulation of cellular component movement
glycosaminoglycan binding
extracellular matrix binding
receptor binding
calcium ion binding
enzyme inhibitor activity
extracellular region
extracellular region part
integral to plasma membrane
0
5
10
15
Z Score

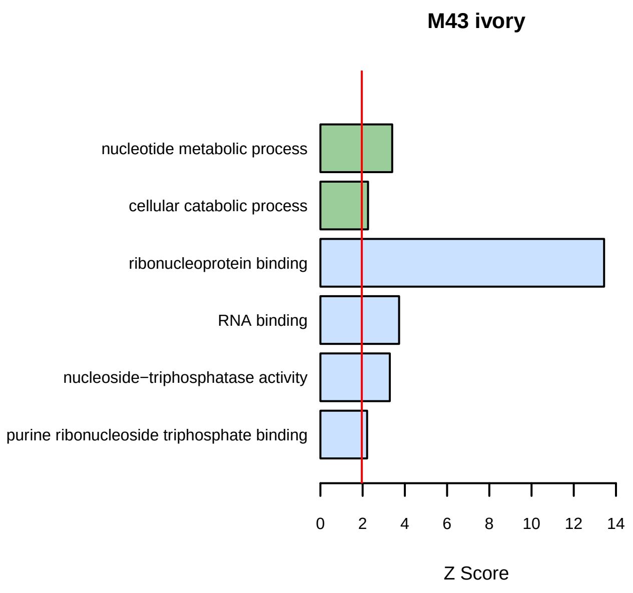
M43 ivory
nucleotide metabolic process
cellular catabolic process
ribonucleoprotein binding
RNA binding
nucleoside−triphosphatase activity
purine ribonucleoside triphosphate binding
0
2
4
6
8
10
12
14
Z Score

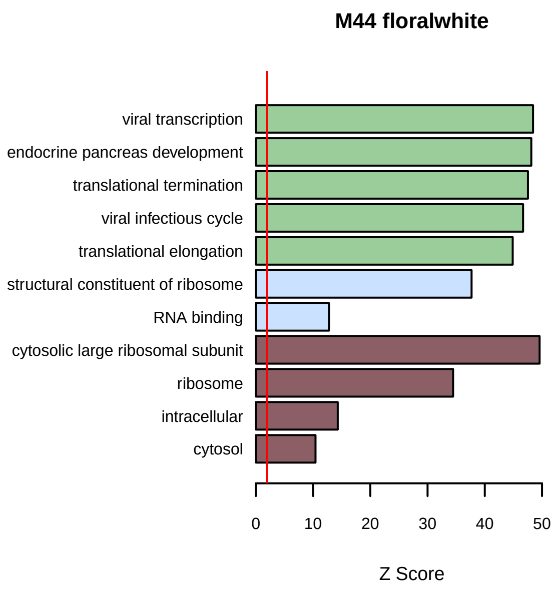
M44 floralwhite
viral transcription
endocrine pancreas development
translational termination
viral infectious cycle
translational elongation
structural constituent of ribosome
RNA binding
cytosolic large ribosomal subunit
ribosome
intracellular
cytosol
0
10
20
30
40
50
Z Score

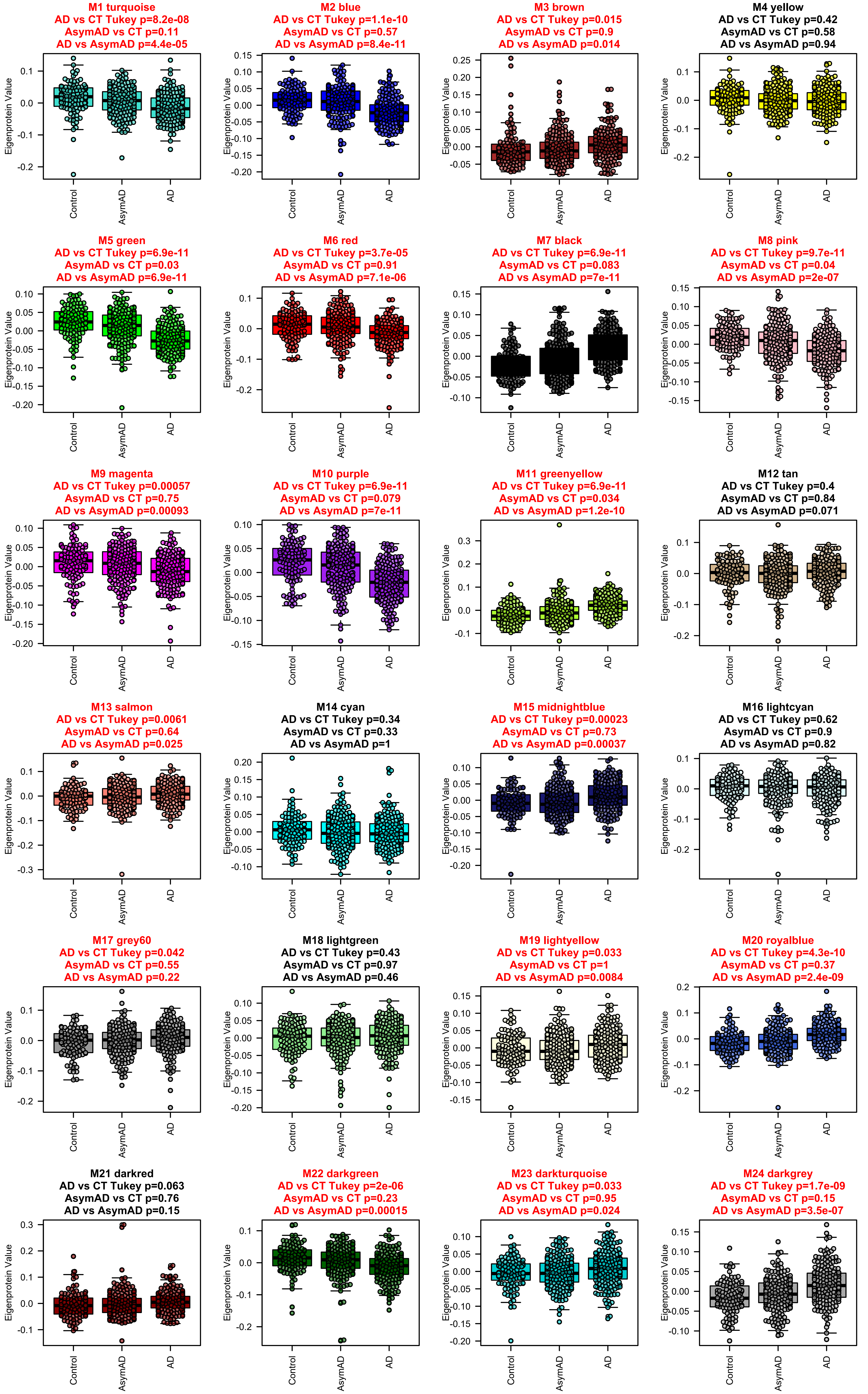
M1 turquoise
AD vs CT Tukey p=8.2e-08
AsymAD vs CT p=0.11
AD vs AsymAD p=4.4e-05
M2 blue
AD vs CT Tukey p=1.1e-10
AsymAD vs CT p=0.57
AD vs AsymAD p=8.4e-11
M3 brown
AD vs CT Tukey p=0.015
AsymAD vs CT p=0.9
AD vs AsymAD p=0.014
M4 yellow
AD vs CT Tukey p=0.42
AsymAD vs CT p=0.58
AD vs AsymAD p=0.94
M5 green
AD vs CT Tukey p=6.9e-11
AsymAD vs CT p=0.03
AD vs AsymAD p=6.9e-11
M6 red
AD vs CT Tukey p=3.7e-05
AsymAD vs CT p=0.91
AD vs AsymAD p=7.1e-06
M7 black
AD vs CT Tukey p=6.9e-11
AsymAD vs CT p=0.083
AD vs AsymAD p=7e-11
M8 pink
AD vs CT Tukey p=9.7e-11
AsymAD vs CT p=0.04
AD vs AsymAD p=2e-07
M9 magenta
AD vs CT Tukey p=0.00057
AsymAD vs CT p=0.75
AD vs AsymAD p=0.00093
M10 purple
AD vs CT Tukey p=6.9e-11
AsymAD vs CT p=0.079
AD vs AsymAD p=7e-11
M11 greenyellow
AD vs CT Tukey p=6.9e-11
AsymAD vs CT p=0.034
AD vs AsymAD p=1.2e-10
M12 tan
AD vs CT Tukey p=0.4
AsymAD vs CT p=0.84
AD vs AsymAD p=0.071
M13 salmon
AD vs CT Tukey p=0.0061
AsymAD vs CT p=0.64
AD vs AsymAD p=0.025
M14 cyan
AD vs CT Tukey p=0.34
AsymAD vs CT p=0.33
AD vs AsymAD p=1
M15 midnightblue
AD vs CT Tukey p=0.00023
AsymAD vs CT p=0.73
AD vs AsymAD p=0.00037
M16 lightcyan
AD vs CT Tukey p=0.62
AsymAD vs CT p=0.9
AD vs AsymAD p=0.82
M17 grey60
AD vs CT Tukey p=0.042
AsymAD vs CT p=0.55
AD vs AsymAD p=0.22
M18 lightgreen
AD vs CT Tukey p=0.43
AsymAD vs CT p=0.97
AD vs AsymAD p=0.46
M19 lightyellow
AD vs CT Tukey p=0.033
AsymAD vs CT p=1
AD vs AsymAD p=0.0084
M20 royalblue
AD vs CT Tukey p=4.3e-10
AsymAD vs CT p=0.37
AD vs AsymAD p=2.4e-09
M21 darkred
AD vs CT Tukey p=0.063
AsymAD vs CT p=0.76
AD vs AsymAD p=0.15
M22 darkgreen
AD vs CT Tukey p=2e-06
AsymAD vs CT p=0.23
AD vs AsymAD p=0.00015
M23 darkturquoise
AD vs CT Tukey p=0.033
AsymAD vs CT p=0.95
AD vs AsymAD p=0.024
M24 darkgrey
AD vs CT Tukey p=1.7e-09
AsymAD vs CT p=0.15
AD vs AsymAD p=3.5e-07
Eigenprotein Value
Control
AsymAD
AD

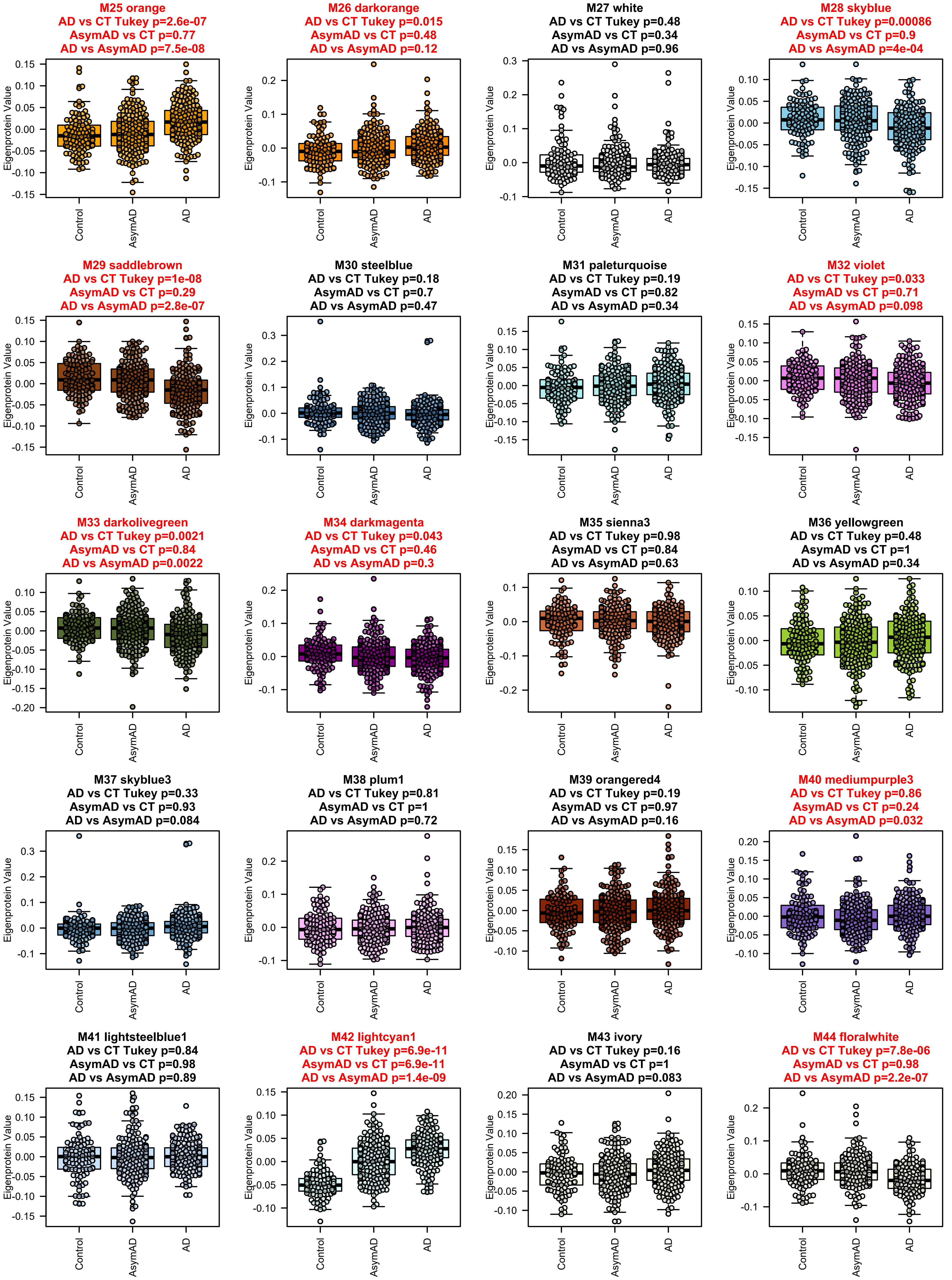
M25 orange
AD vs CT Tukey p=2.6e-07
AsymAD vs CT p=0.77
AD vs AsymAD p=7.5e-08
M26 darkorange
AD vs CT Tukey p=0.015
AsymAD vs CT p=0.48
AD vs AsymAD p=0.12
M27 white
AD vs CT Tukey p=0.48
AsymAD vs CT p=0.34
AD vs AsymAD p=0.96
M28 skyblue
AD vs CT Tukey p=0.00086
AsymAD vs CT p=0.9
AD vs AsymAD p=4e-04
M29 saddlebrown
AD vs CT Tukey p=1e-08
AsymAD vs CT p=0.29
AD vs AsymAD p=2.8e-07
M30 steelblue
AD vs CT Tukey p=0.18
AsymAD vs CT p=0.7
AD vs AsymAD p=0.47
M31 paleturquoise
AD vs CT Tukey p=0.19
AsymAD vs CT p=0.82
AD vs AsymAD p=0.34
M32 violet
AD vs CT Tukey p=0.033
AsymAD vs CT p=0.71
AD vs AsymAD p=0.098
M33 darkolivegreen
AD vs CT Tukey p=0.0021
AsymAD vs CT p=0.84
AD vs AsymAD p=0.0022
M34 darkmagenta
AD vs CT Tukey p=0.043
AsymAD vs CT p=0.46
AD vs AsymAD p=0.3
M35 sienna3
AD vs CT Tukey p=0.98
AsymAD vs CT p=0.84
AD vs AsymAD p=0.63
M36 yellowgreen
AD vs CT Tukey p=0.48
AsymAD vs CT p=1
AD vs AsymAD p=0.34
M37 skyblue3
AD vs CT Tukey p=0.33
AsymAD vs CT p=0.93
AD vs AsymAD p=0.084
M38 plum1
AD vs CT Tukey p=0.81
AsymAD vs CT p=1
AD vs AsymAD p=0.72
M39 orangered4
AD vs CT Tukey p=0.19
AsymAD vs CT p=0.97
AD vs AsymAD p=0.16
M40 mediumpurple3
AD vs CT Tukey p=0.86
AsymAD vs CT p=0.24
AD vs AsymAD p=0.032
M41 lightsteelblue1
AD vs CT Tukey p=0.84
AsymAD vs CT p=0.98
AD vs AsymAD p=0.89
M42 lightcyan1
AD vs CT Tukey p=6.9e-11
AsymAD vs CT p=6.9e-11
AD vs AsymAD p=1.4e-09
M43 ivory
AD vs CT Tukey p=0.16
AsymAD vs CT p=1
AD vs AsymAD p=0.083
M44 floralwhite
AD vs CT Tukey p=7.8e-06
AsymAD vs CT p=0.98
AD vs AsymAD p=2.2e-07
Eigenprotein Value
Control
AsymAD
AD

**Module 1, n = 558,**
**FDR Corrected pvalue 3.6797e-07**

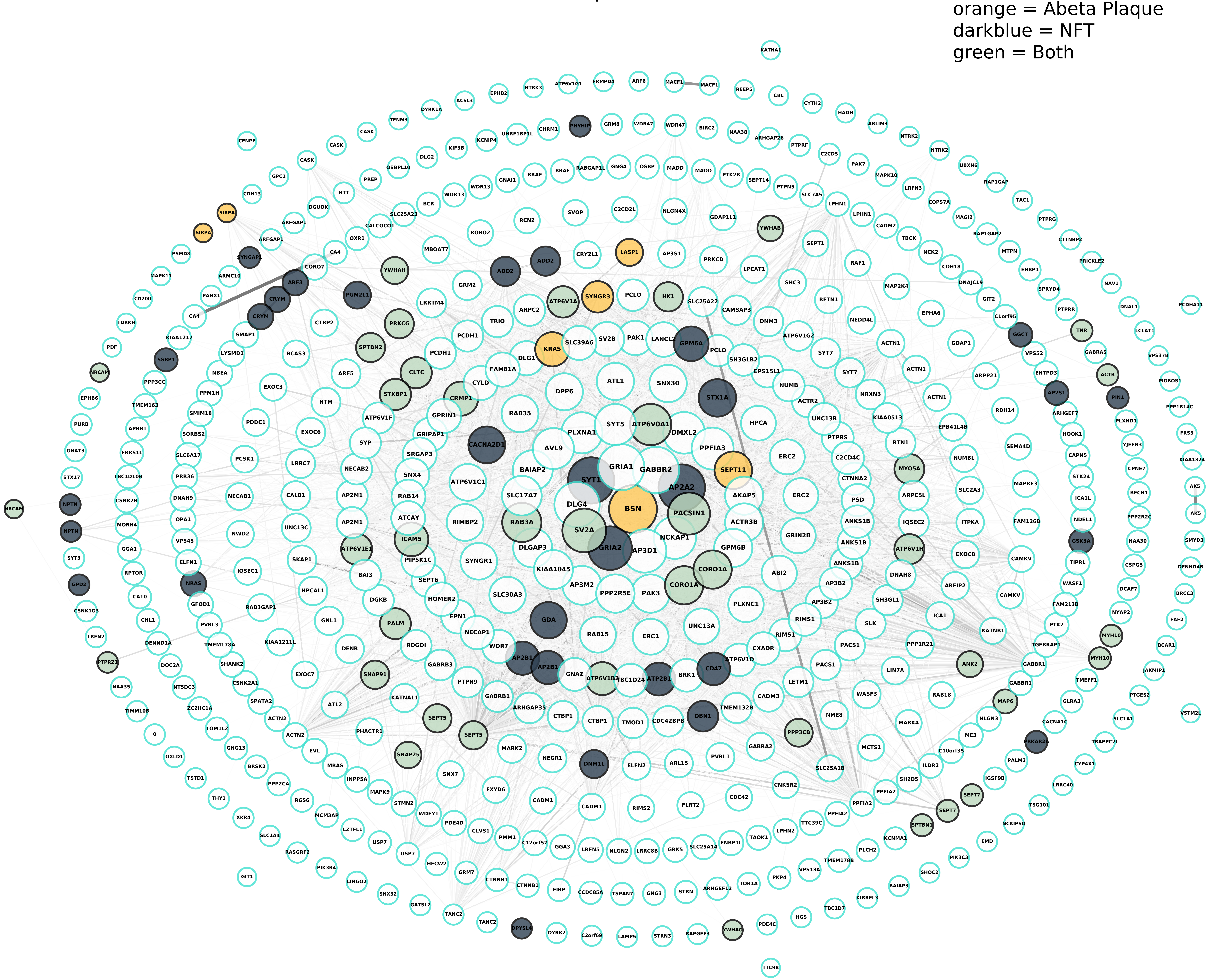

**Module 2, n = 403,**
**FDR Corrected pvalue 1.6412e-14**

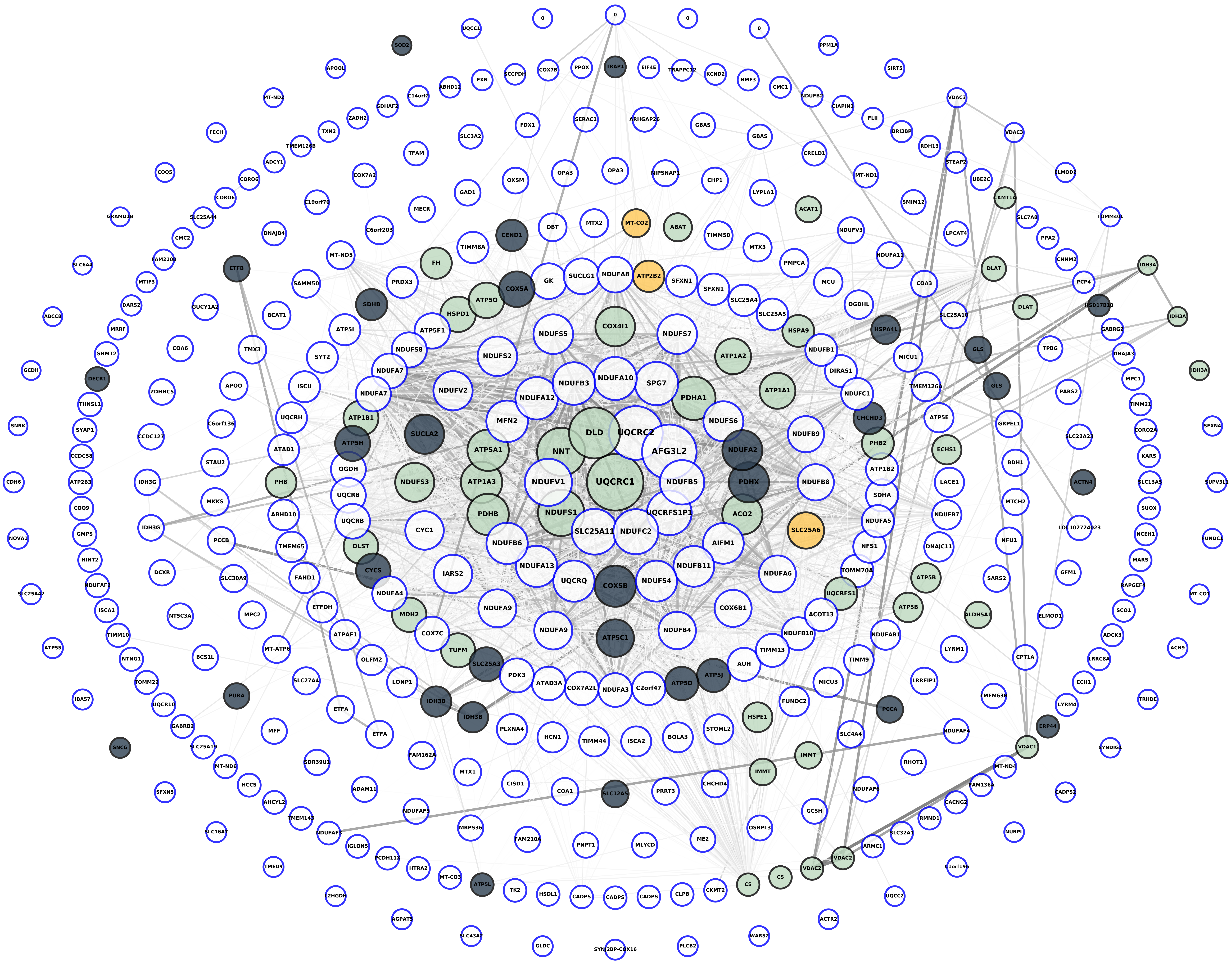

**Module 3, n = 398,**
**FDR Corrected pvalue 0.18123**

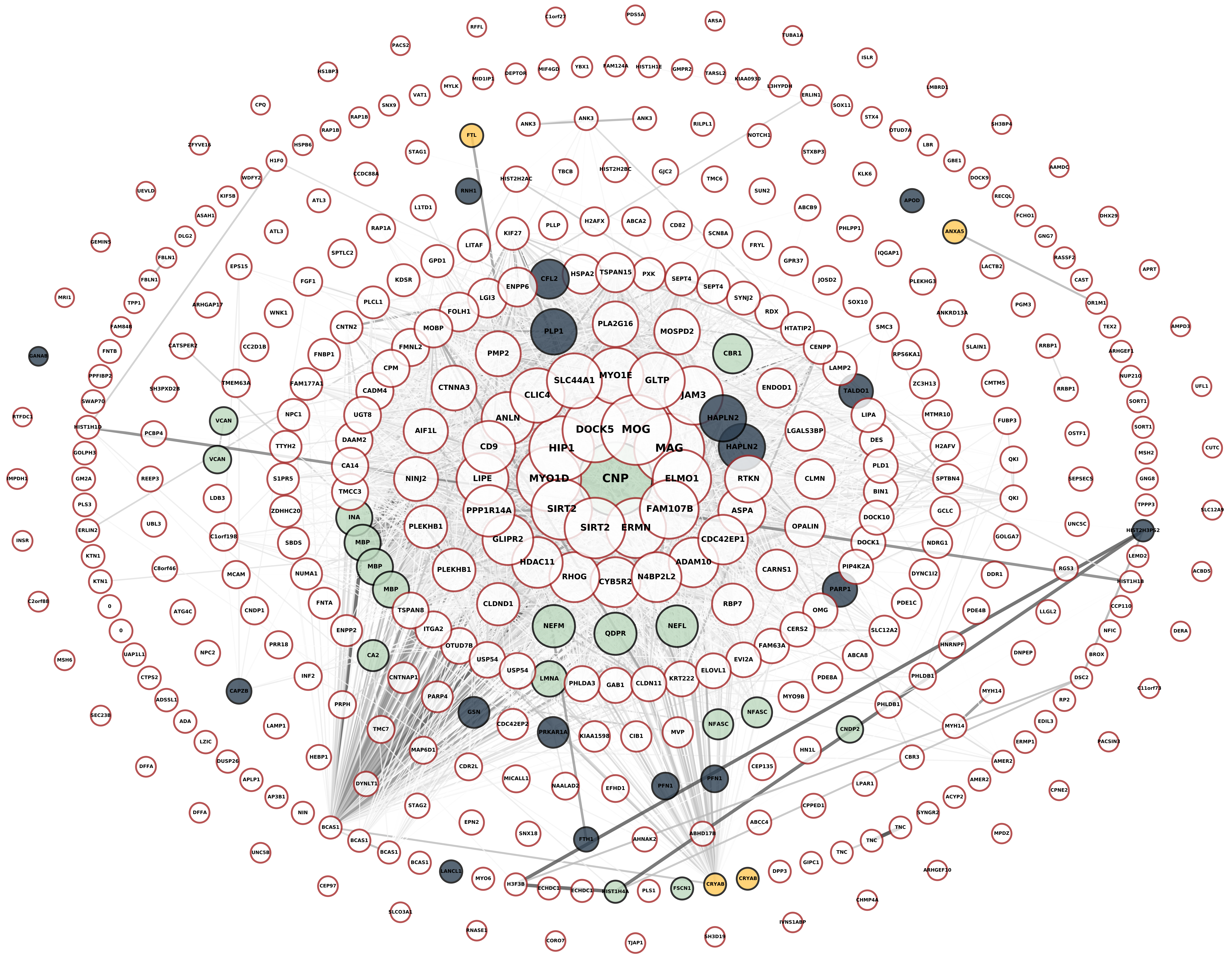

**Module 4, n = 291,**
**FDR Corrected pvalue 1.6684e-07**

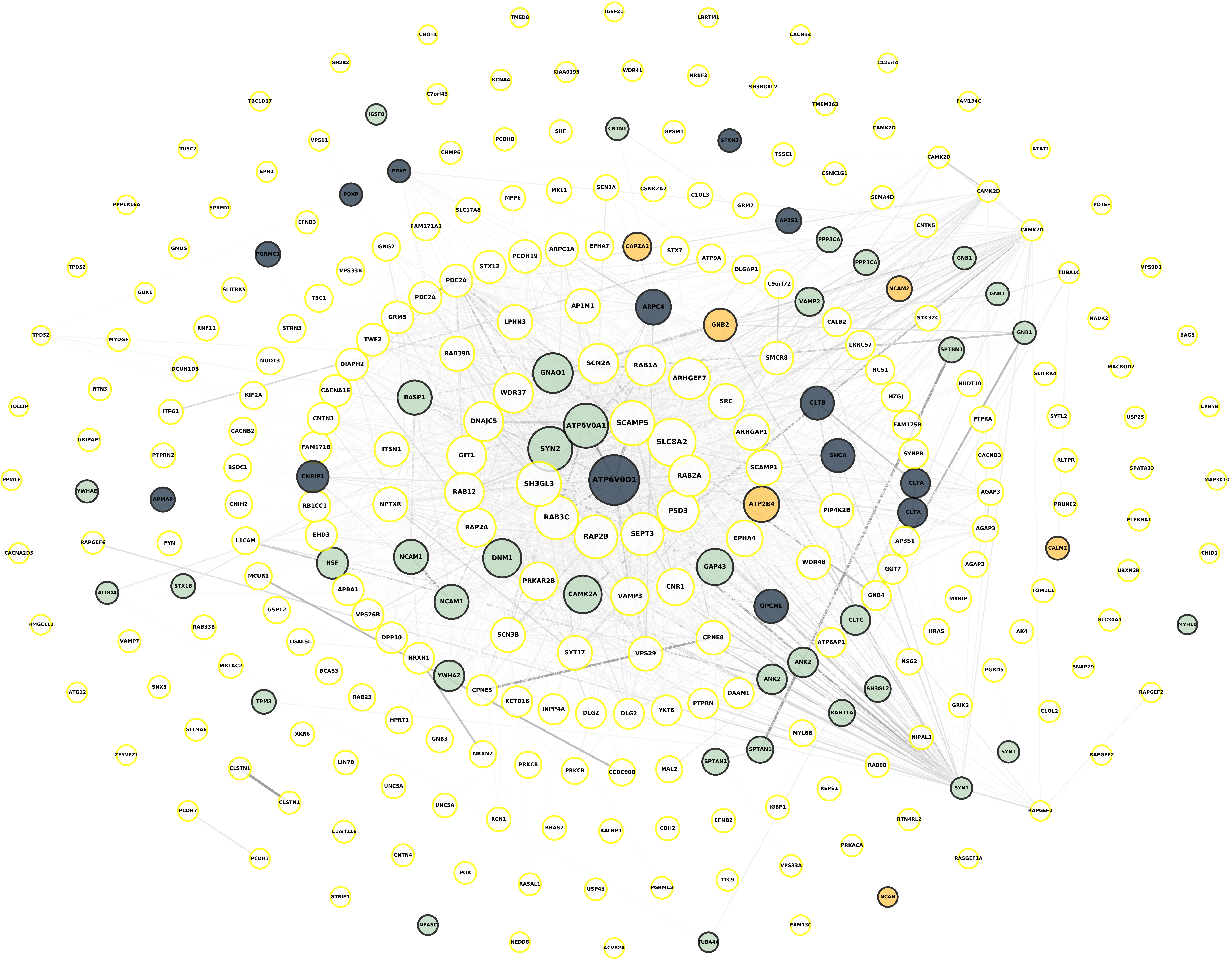

**Module 5, n = 249,**
**FDR Corrected pvalue 1**

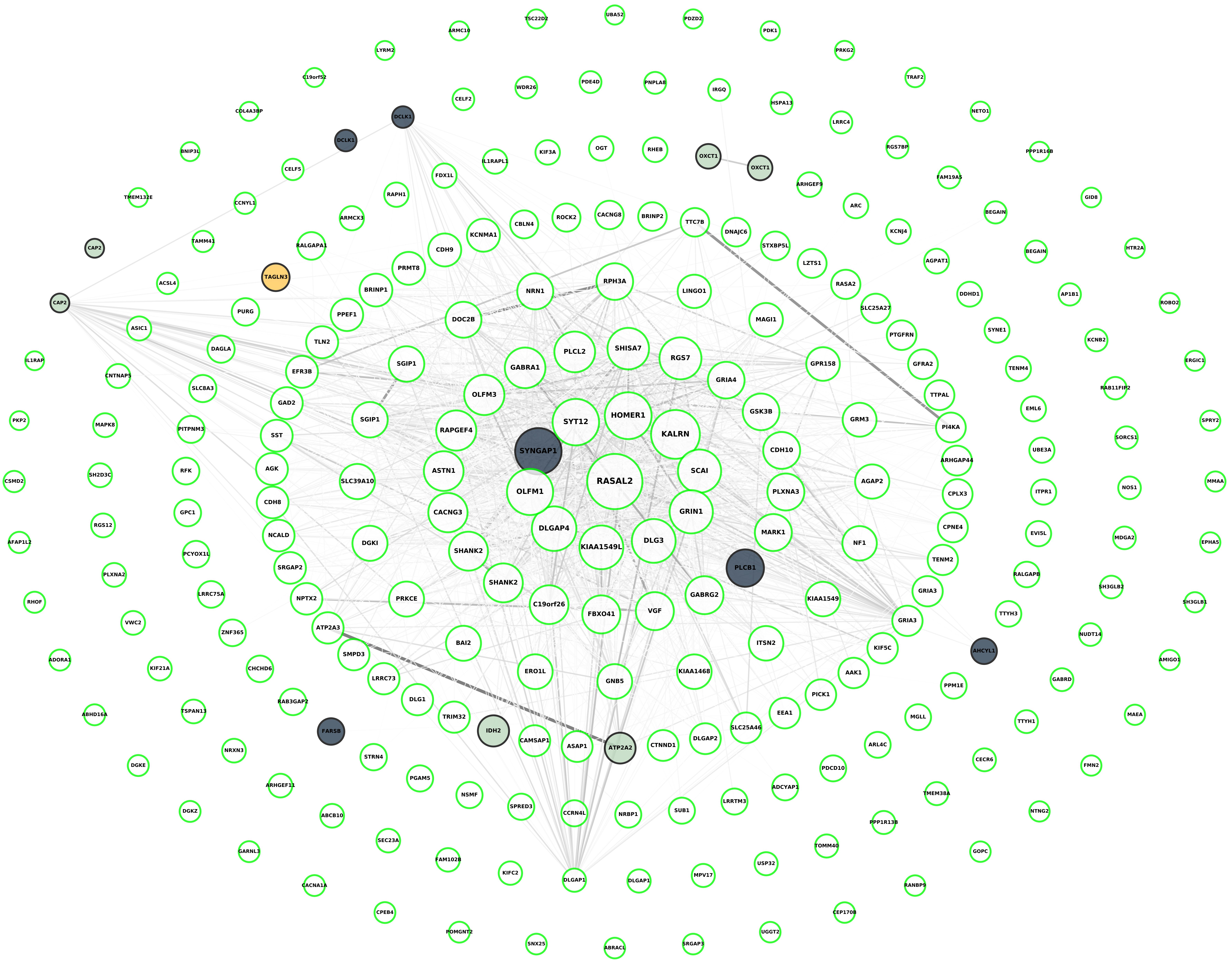

**Module 6, n = 248,
FDR Corrected pvalue 1**

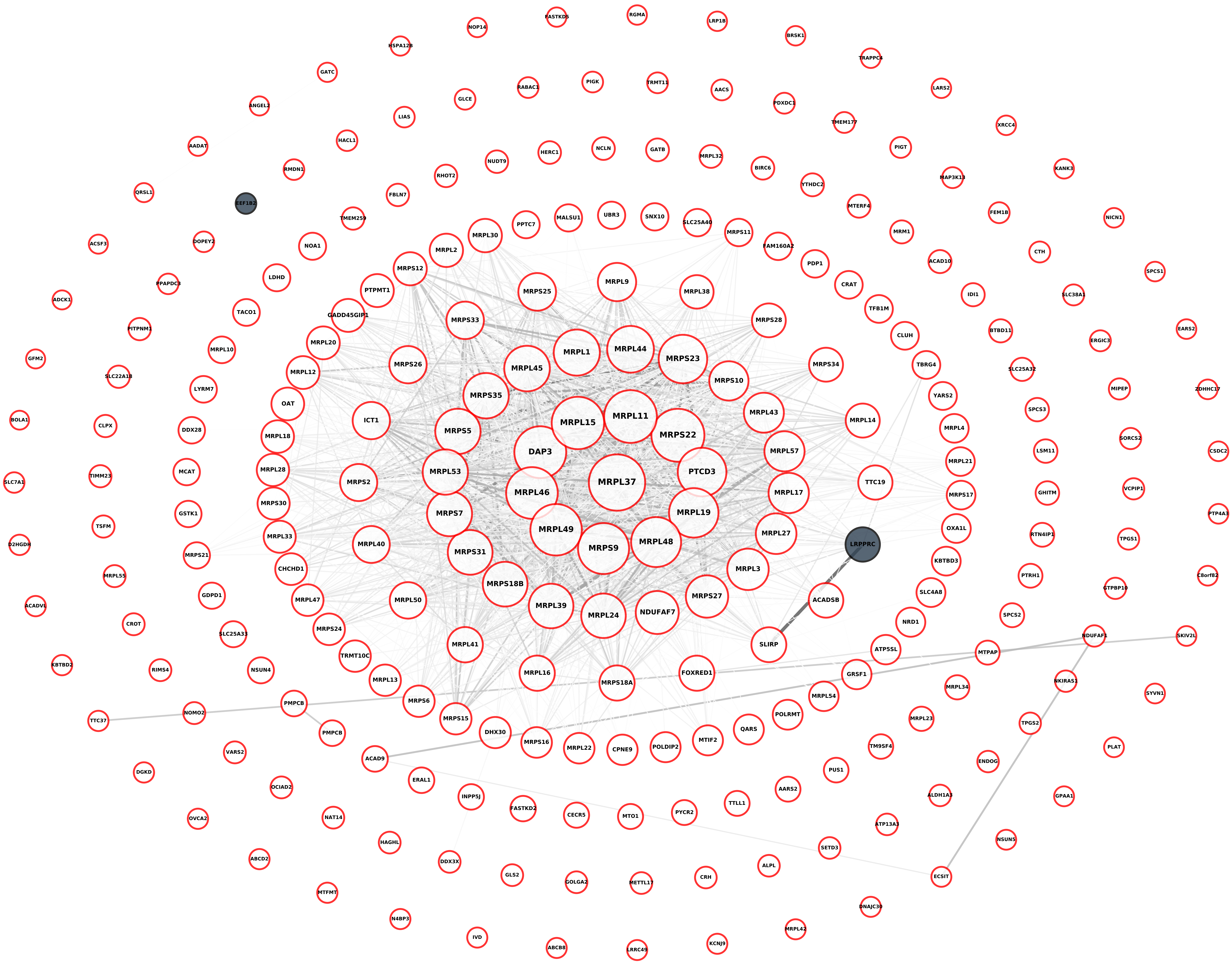

**Module 7, n = 234,**
**FDR Corrected pvalue 0.00044895**

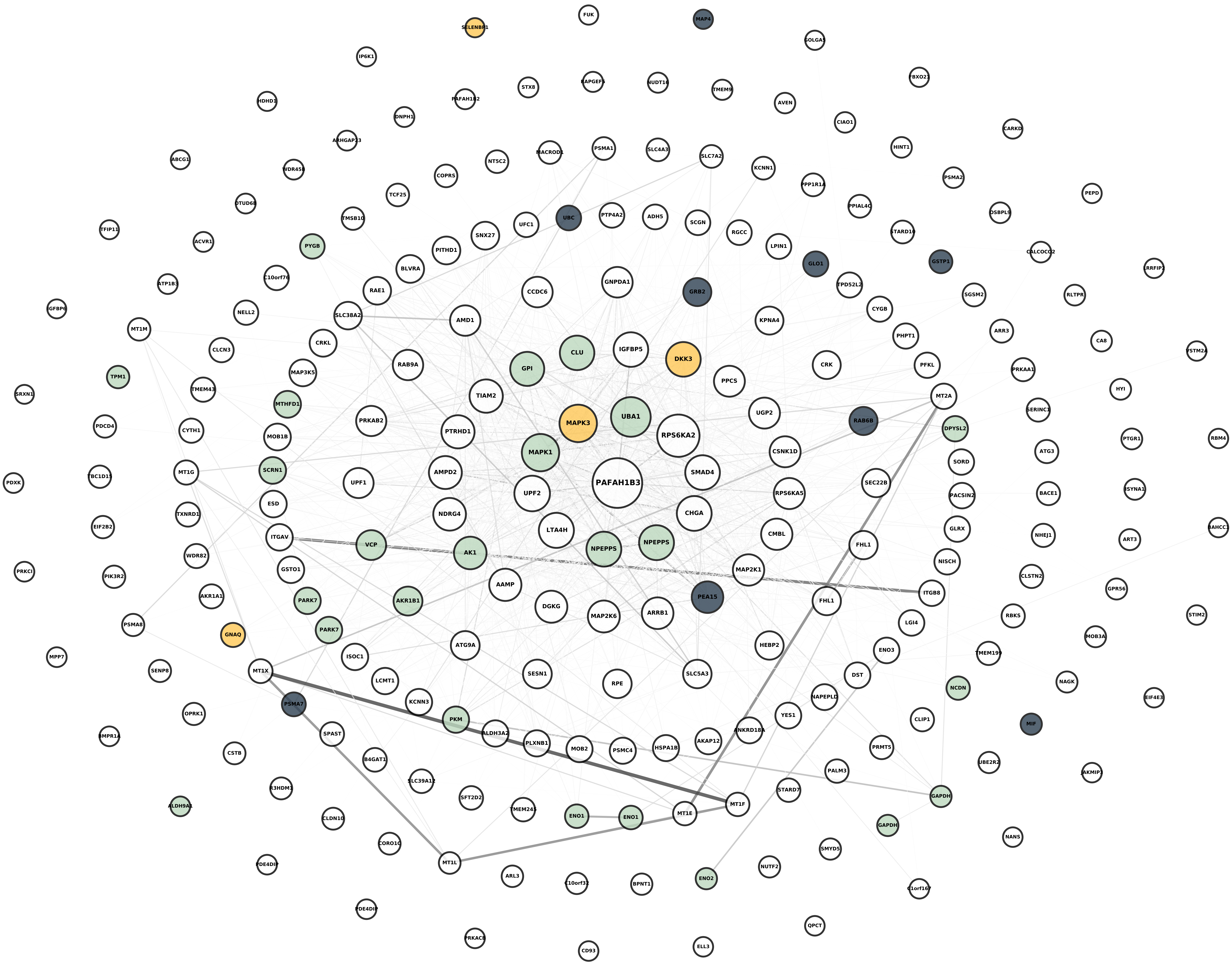

**Module 8, n = 232,**
**FDR Corrected pvalue 0.30558**

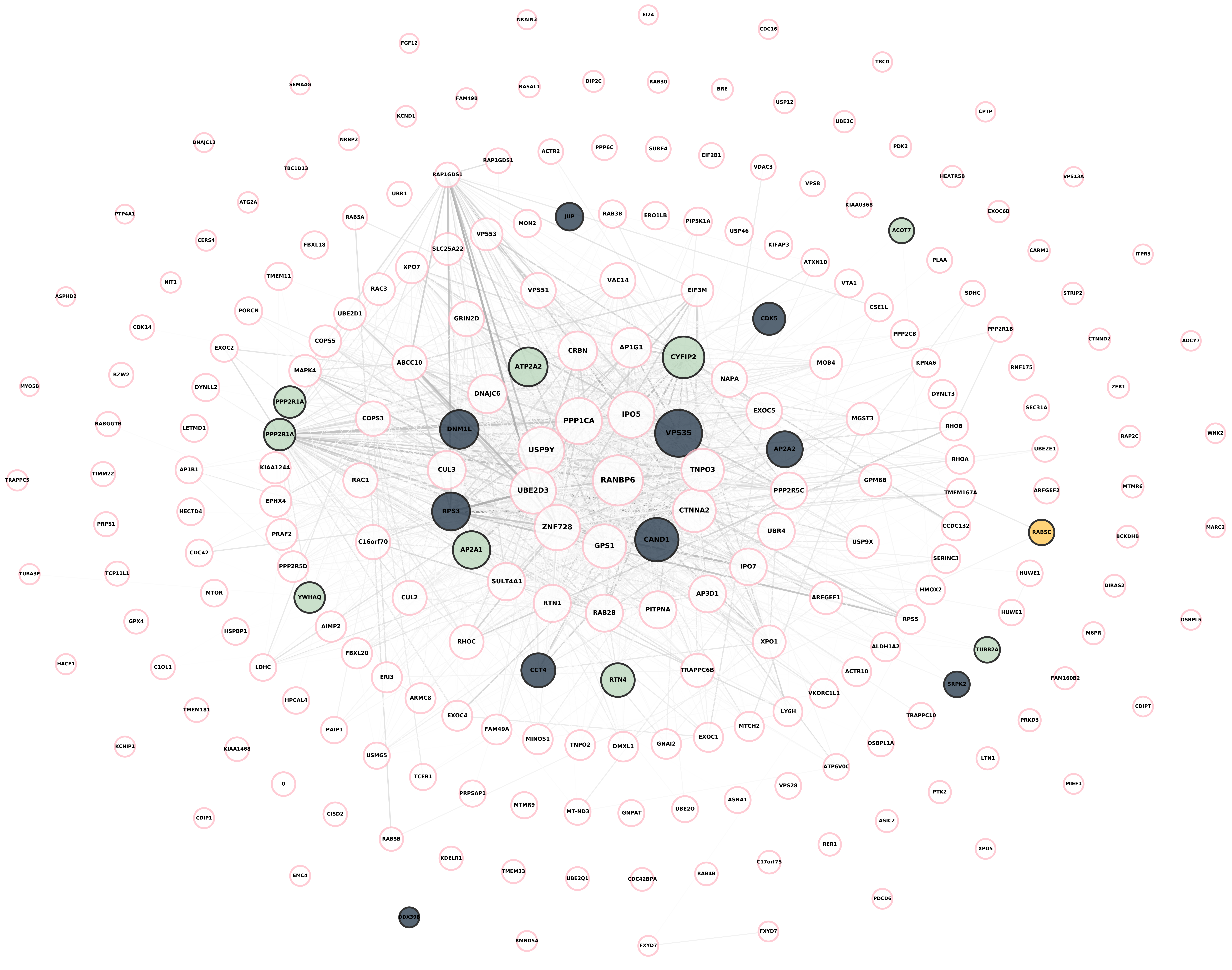

### Module 9, n = 227,
### FDR Corrected pvalue 1

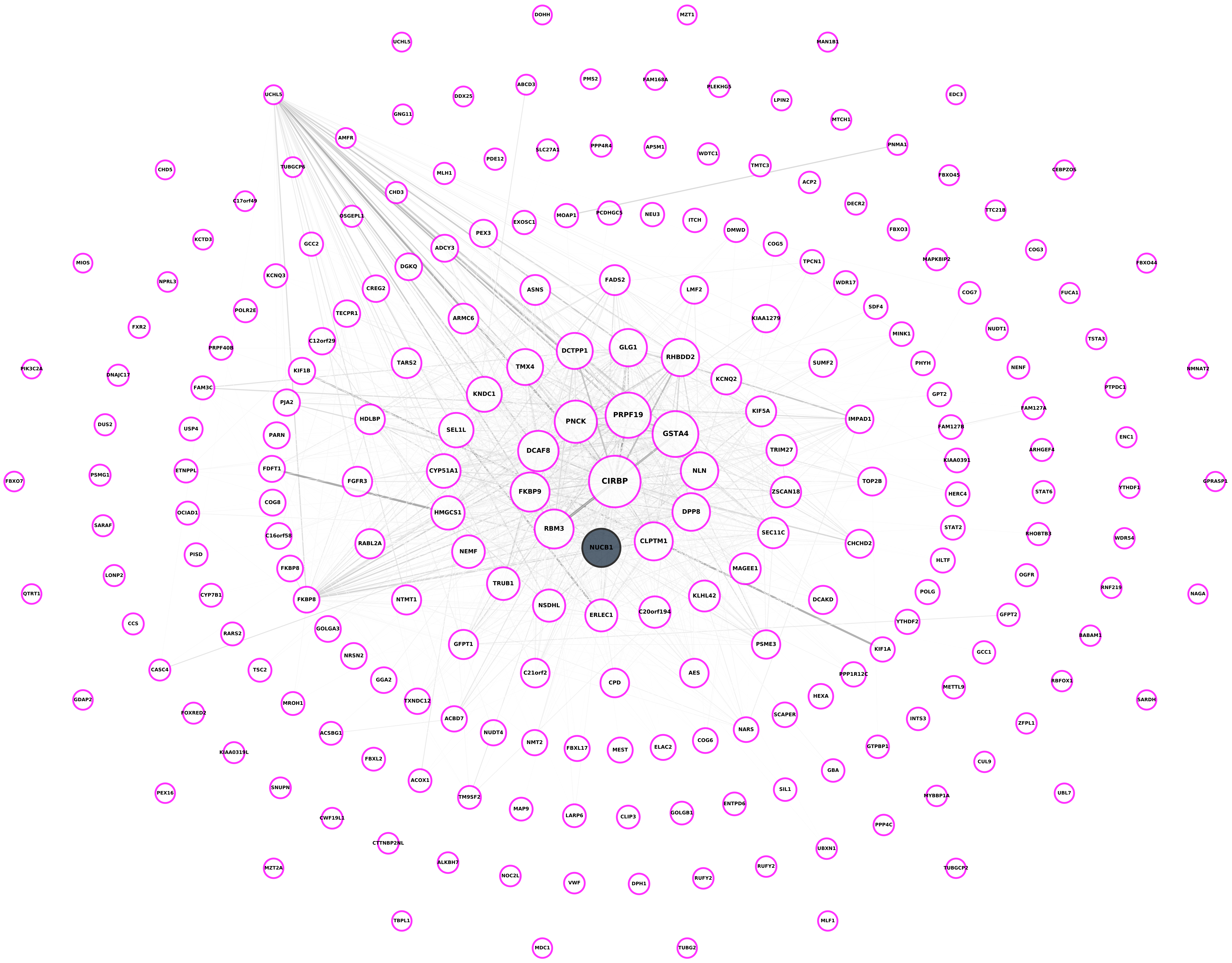

**Module 10, n = 200,**
**FDR Corrected pvalue 1**

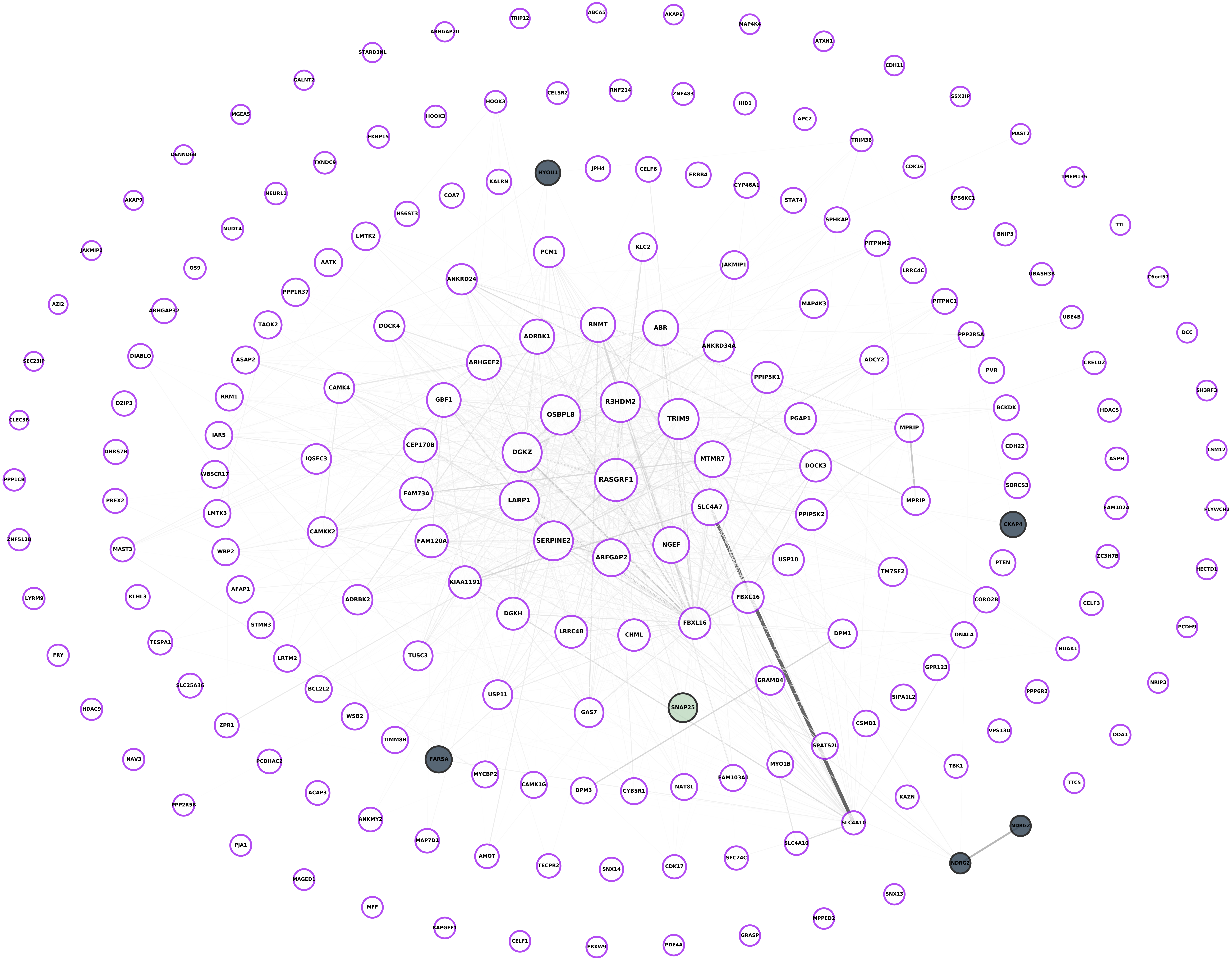

**Module 11, n = 198,**
**FDR Corrected pvalue 0.094128**

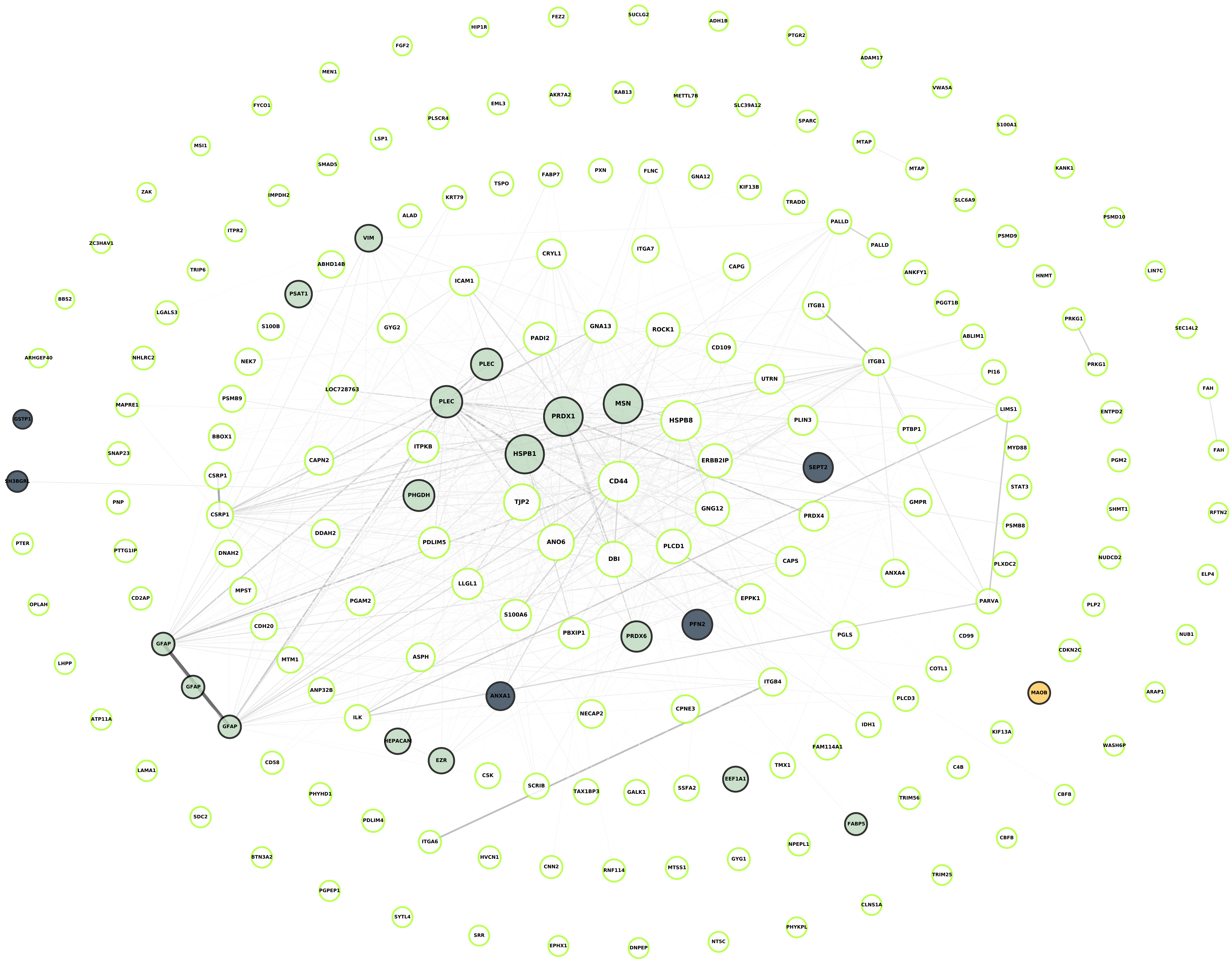

**Module 12, n = 164,
FDR Corrected pvalue 1**

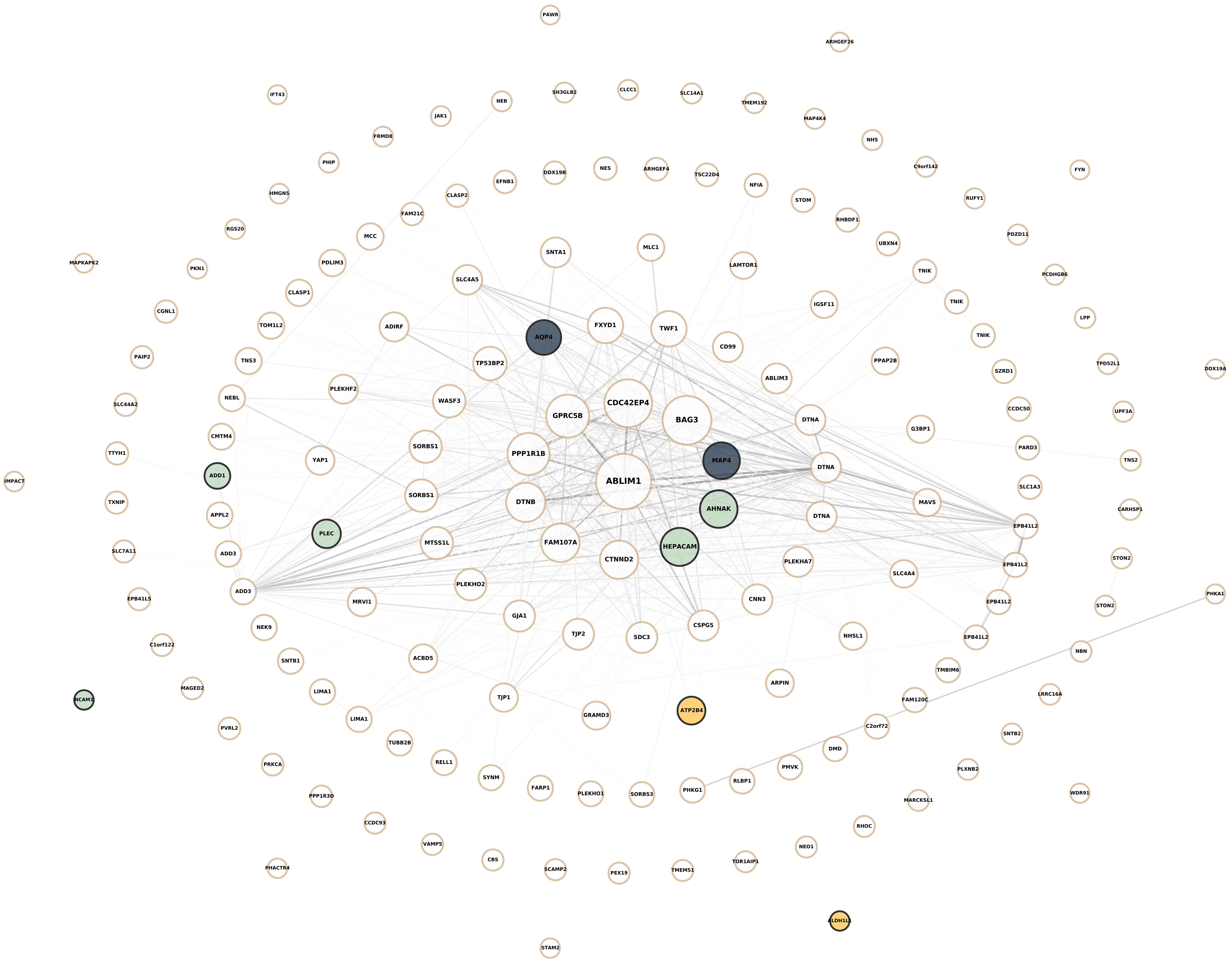

**Module 13, n = 162,**
**FDR Corrected pvalue 1.1211e-05**

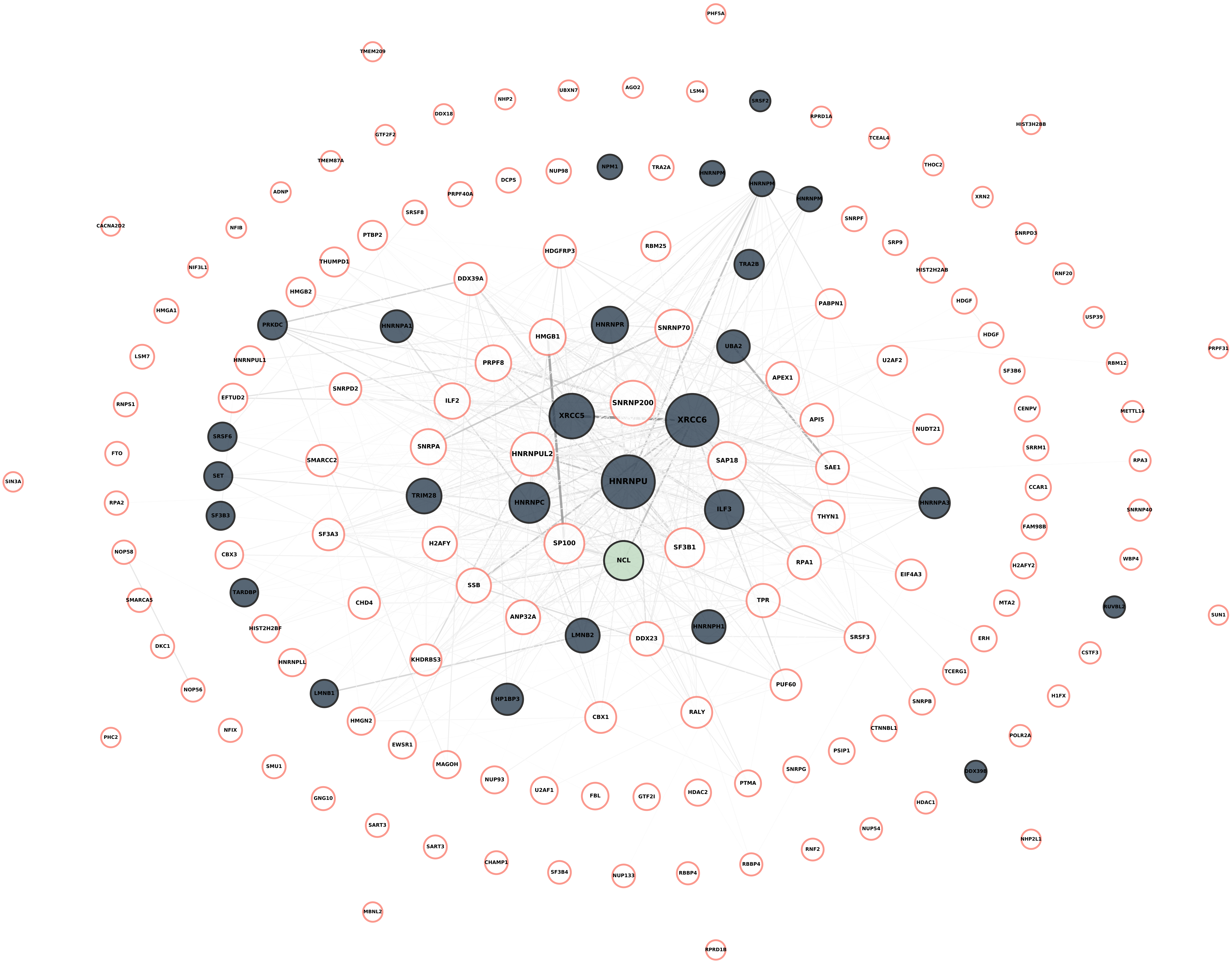

**Module 14, n = 162,**
**FDR Corrected pvalue 4.448e-14**

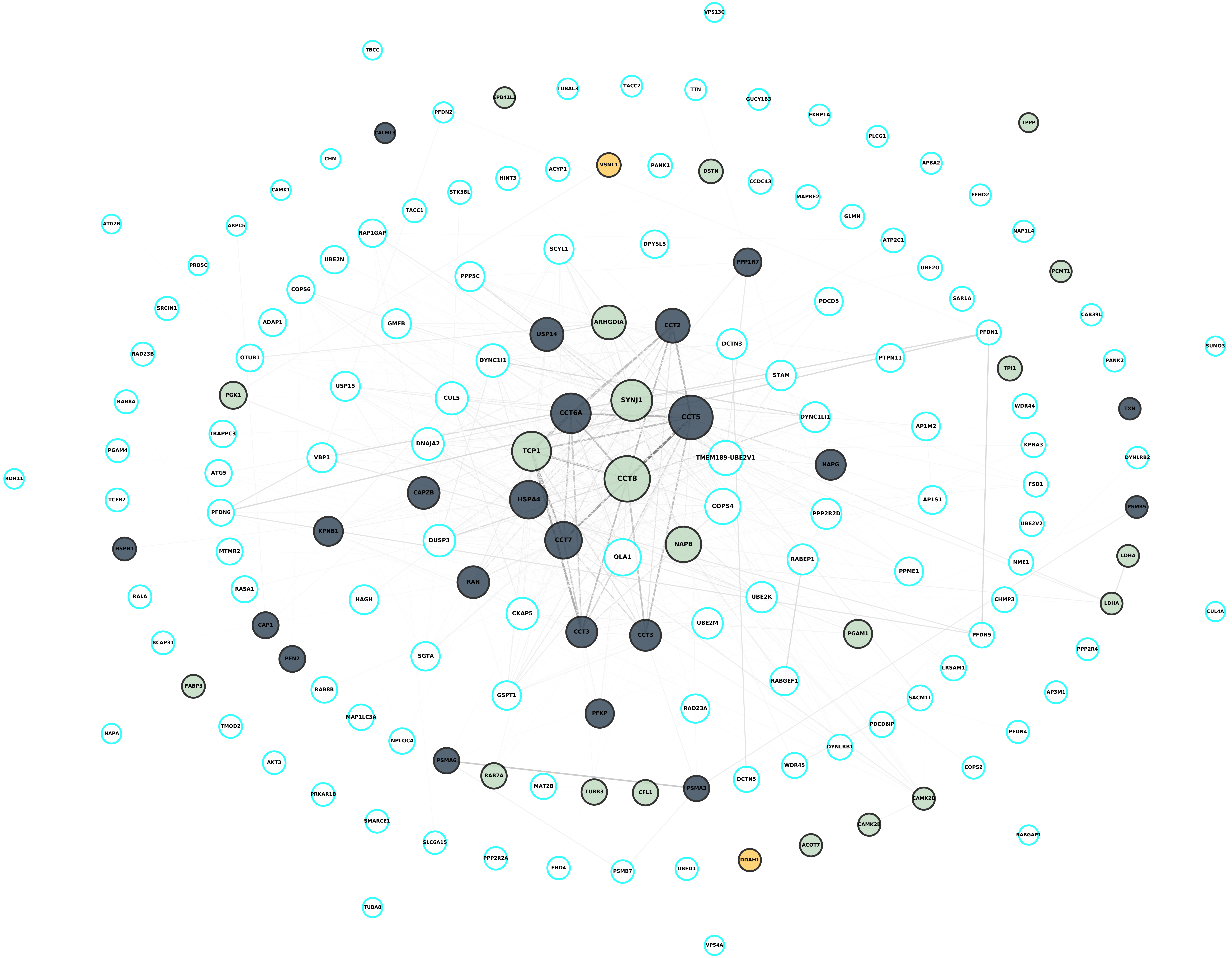

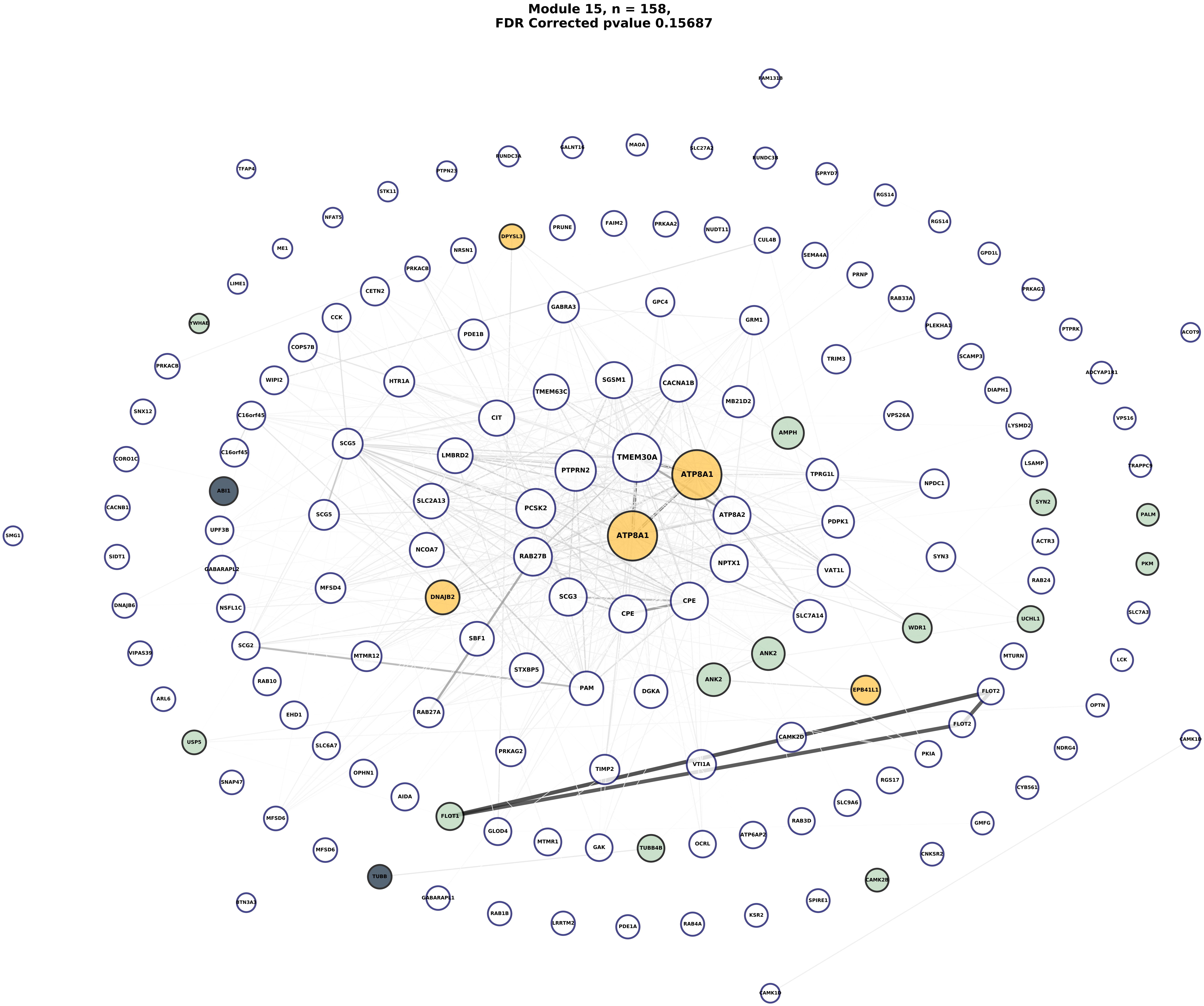
Module 15, n = 158,
FDR Corrected pvalue 0.15687
FAM131B
RUNDC3A
GALNT16
MAOA
SLC27A2
RUNDC3B
TFAP4
PTPN23
SPRYD7
STK11
RGS14
NFAT5
PRUNE
FAIM2
PRKAA2
NUDT11
RGS14
DPYSL3
CUL4B
ME1
NRSN1
SEMA4A
GPD1L
PRKACB
PRNP
LIME1
CETN2
RAB33A
PRKAG1
GABRA3
GPC4
YWHAE
CCK
GRM1
PLEKHA1
PTPRK
ACOT9
PDE1B
COPS7B
TRIM3
SCAMP3
PRKACB
ADCYAP1R1
SGSM1
WIPI2
HTR1A
CACNA1B
TMEM63C
DIAPH1
MB21D2
SNX12
C16orf45
CIT
VPS26A
VPS16
AMPH
LYSMD2
SCG5
C16orf45
LMBRD2
TMEM30A
CORO1C
PTPRN2
LSAMP
ATP8A1
TRAPPC9
TPRG1L
NPDC1
ABI1
SLC2A13
CACNB1
PCSK2
SYN2
ATP8A2
PALM
SCG5
UPF3B
PDPK1
SMG1
ATP8A1
ACTR3
NCOA7
SIDT1
RAB27B
SYN3
NPTX1
VAT1L
PKM
GABARAPL2
MFSD4
RAB24
DNAJB6
DNAJB2
SCG3
CPE
NSFL1C
SLC7A3
CPE
SLC7A14
UCHL1
WDR1
SBF1
SCG2
ANK2
VIPAS39
MTMR12
MTURN
LCK
STXBP5
RAB10
ANK2
PAM
DGKA
FLOT2
EPB41L1
ARL6
OPTN
EHD1
RAB27A
FLOT2
CAMK2D
CAMK1D
USP5
SLC6A7
PKIA
NDRG4
PRKAG2
VTI1A
TIMP2
RGS17
OPHN1
SNAP47
CYB561
SLC9A6
AIDA
MFSD6
FLOT1
RAB3D
GMFG
GLOD4
ATP6AP2
MTMR1
GAK
TUBB4B
OCRL
MFSD6
CNKSR2
TUBB
CAMK2B
GABARAPL1
SPIRE1
BTN3A3
RAB1B
LRRTM2
KSR2
PDE1A
RAB4A
CAMK1D

Module 16, n = 154,
FDR Corrected pvalue 1

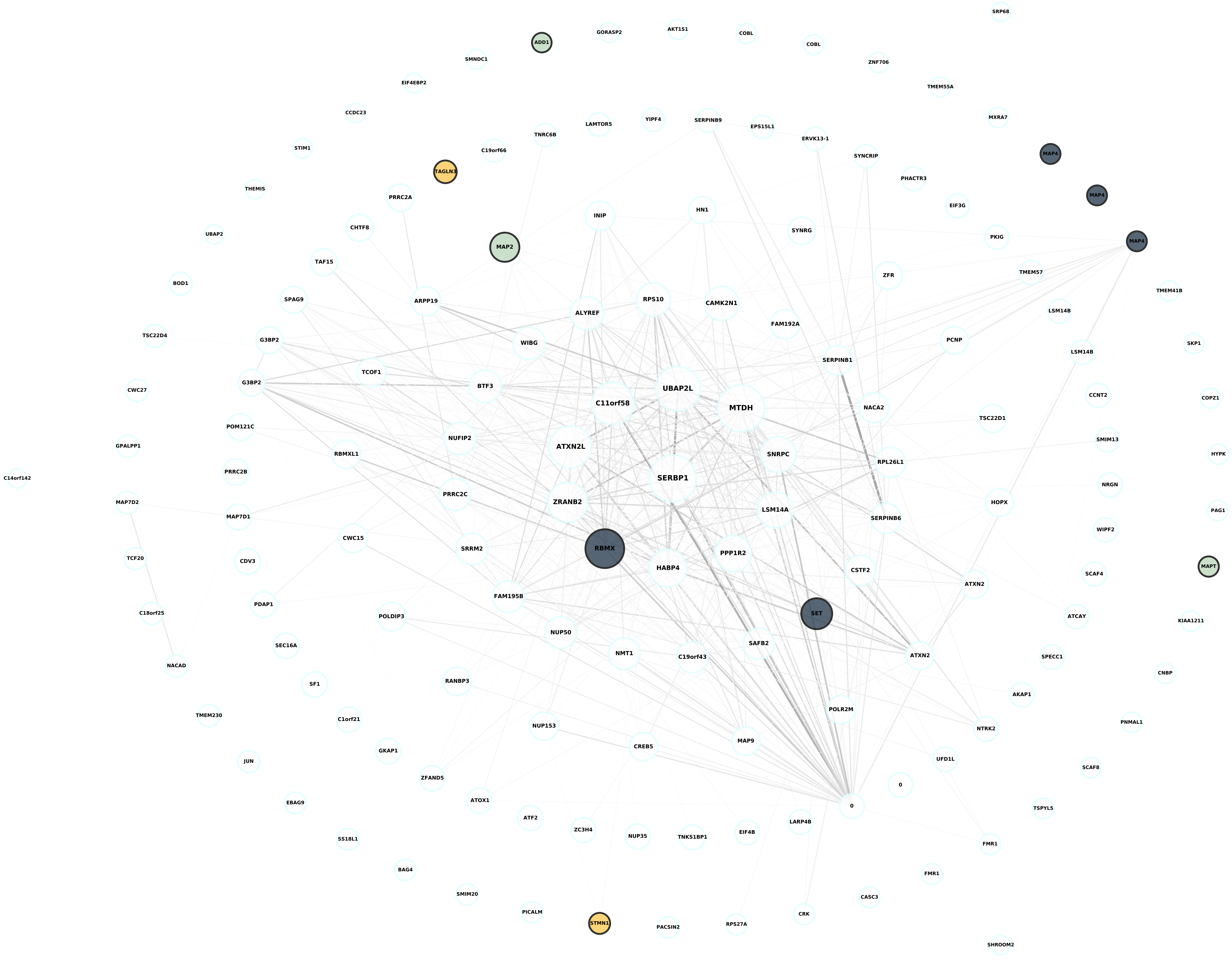

**Module 17, n = 153,**
**FDR Corrected pvalue 1**

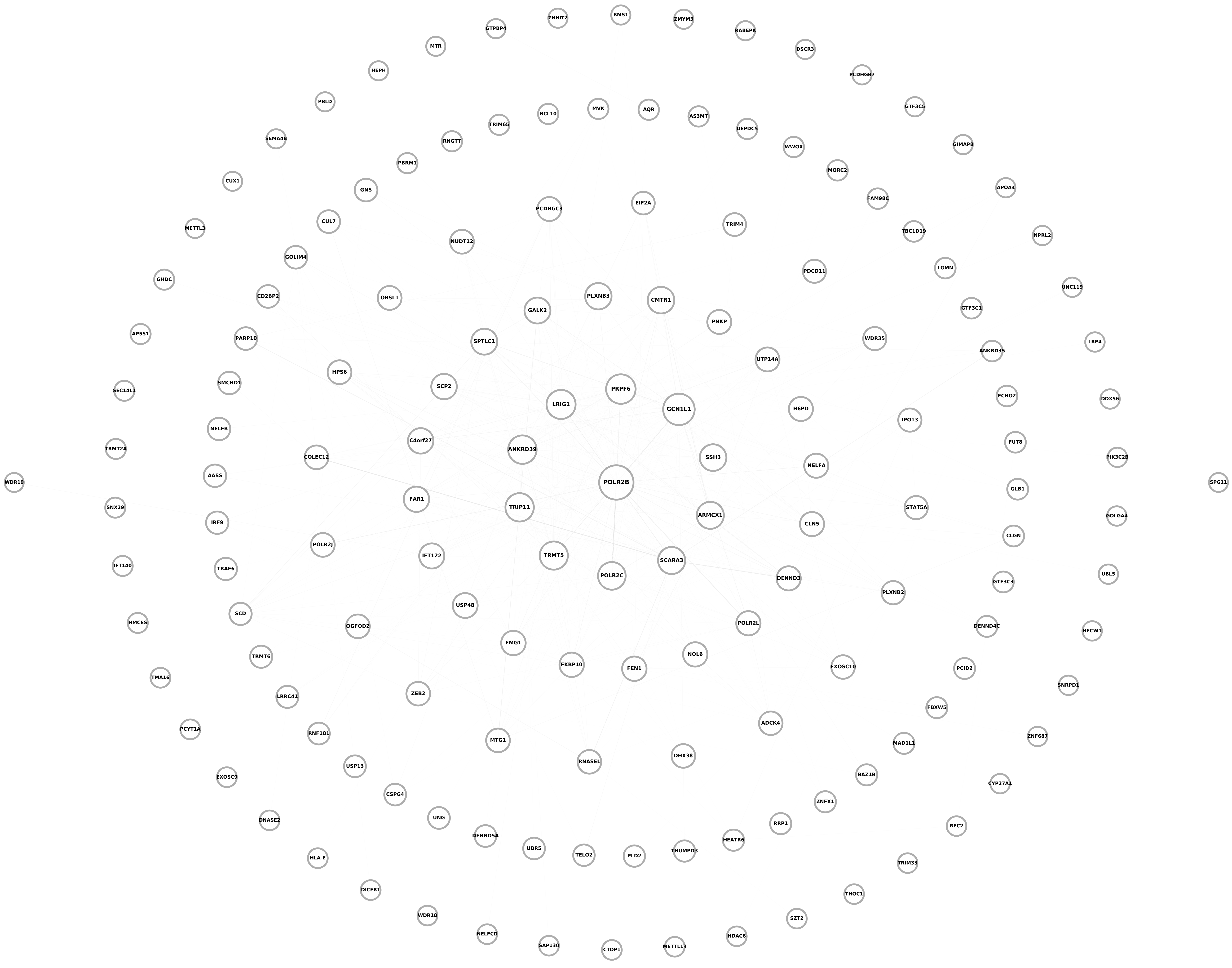

**Module 18, n = 148,**
**FDR Corrected pvalue 0.074783**

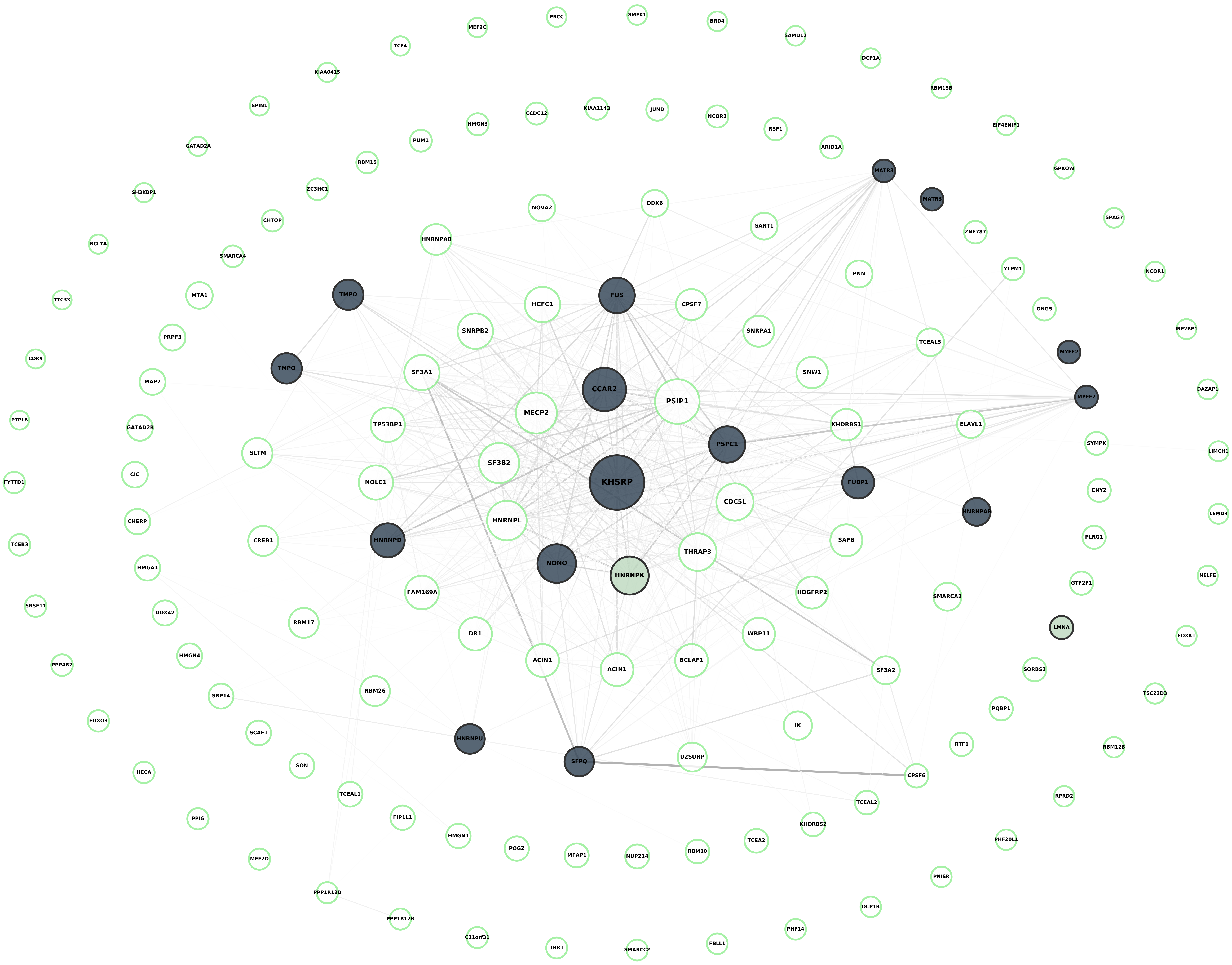

**Module 19, n = 142,
FDR Corrected pvalue 1**

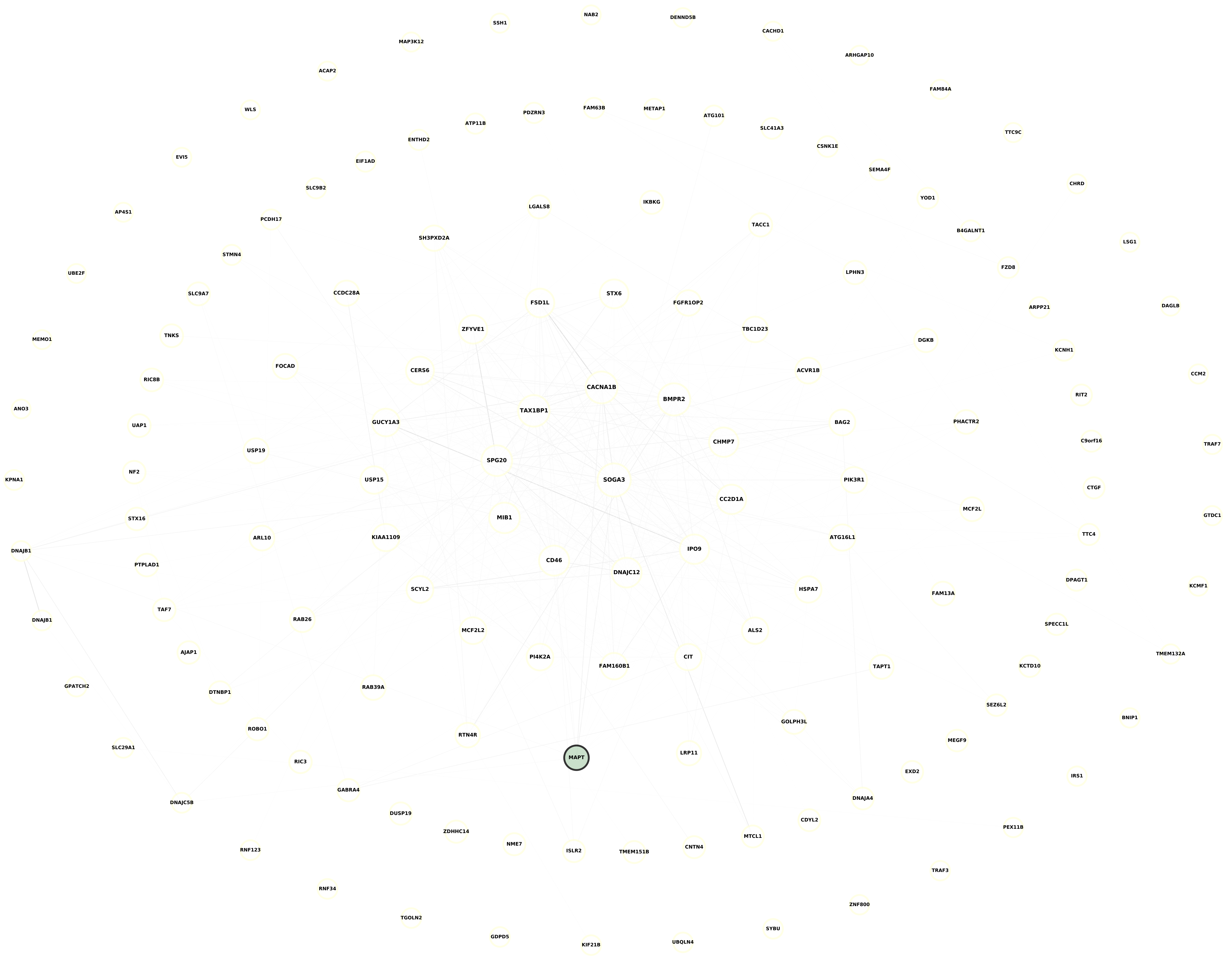

**Module 20, n = 141,**
**FDR Corrected pvalue 0.19461**

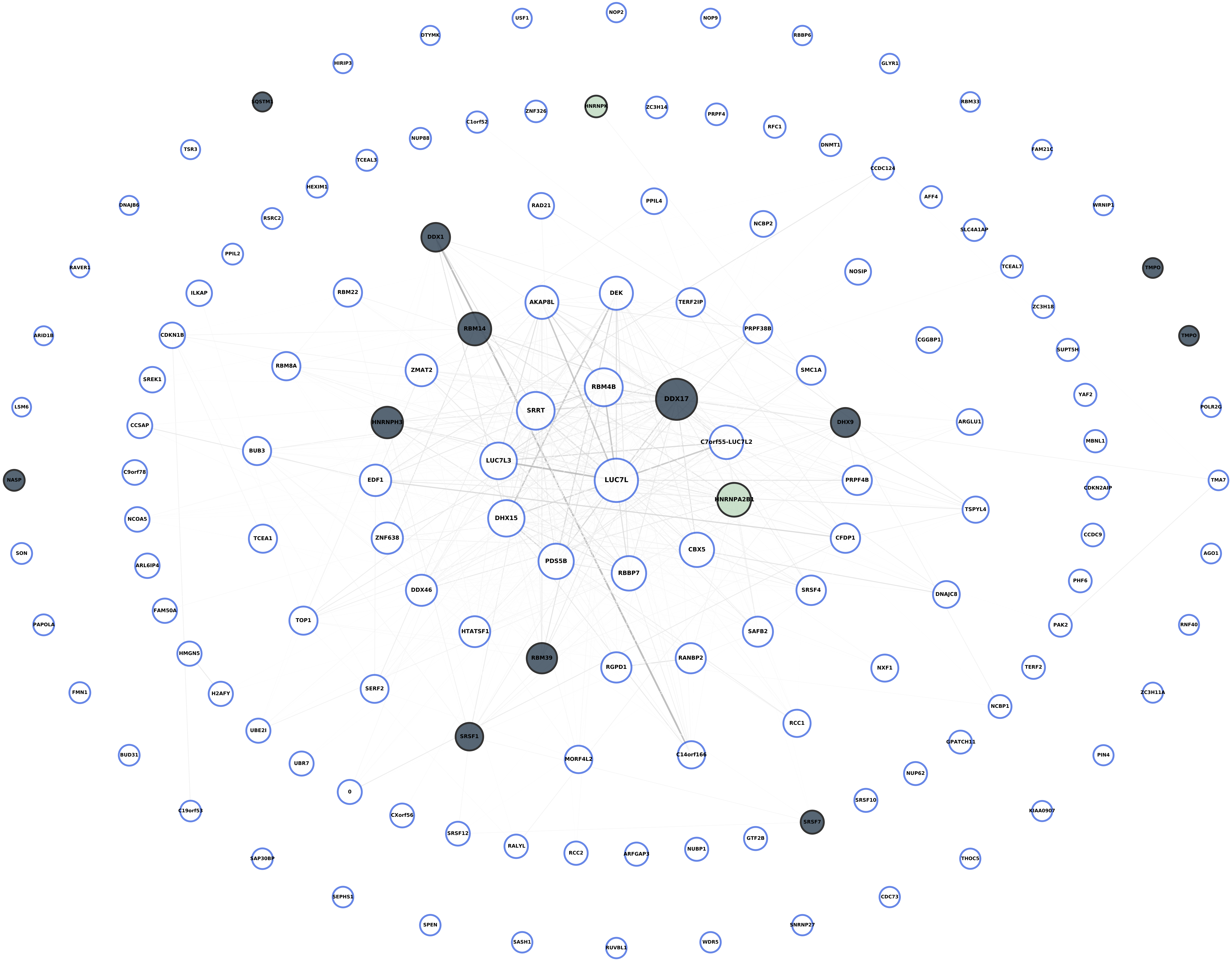

**Module 21, n = 126,**
**FDR Corrected pvalue 1**

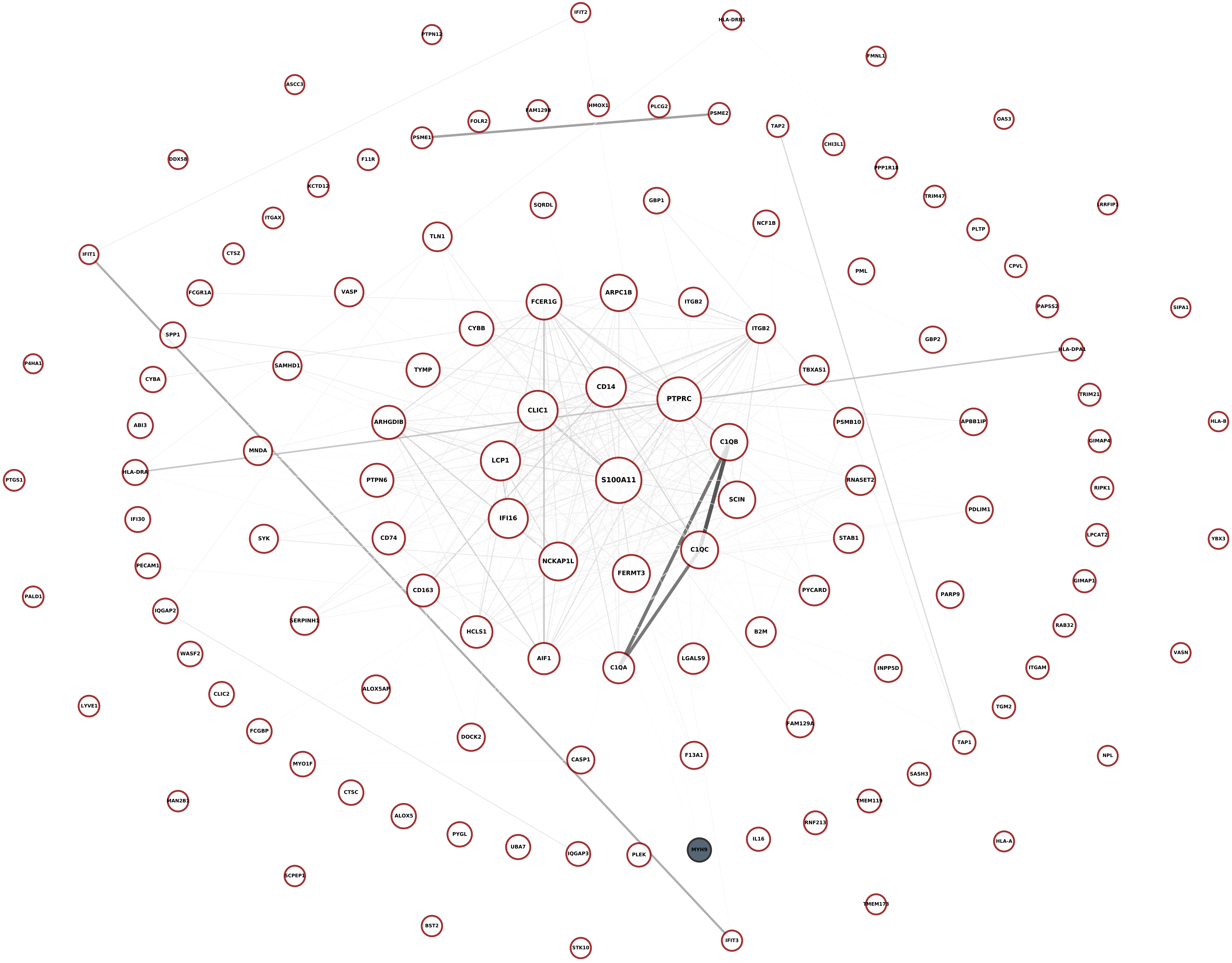

**Module 22, n = 118,
FDR Corrected pvalue 1**

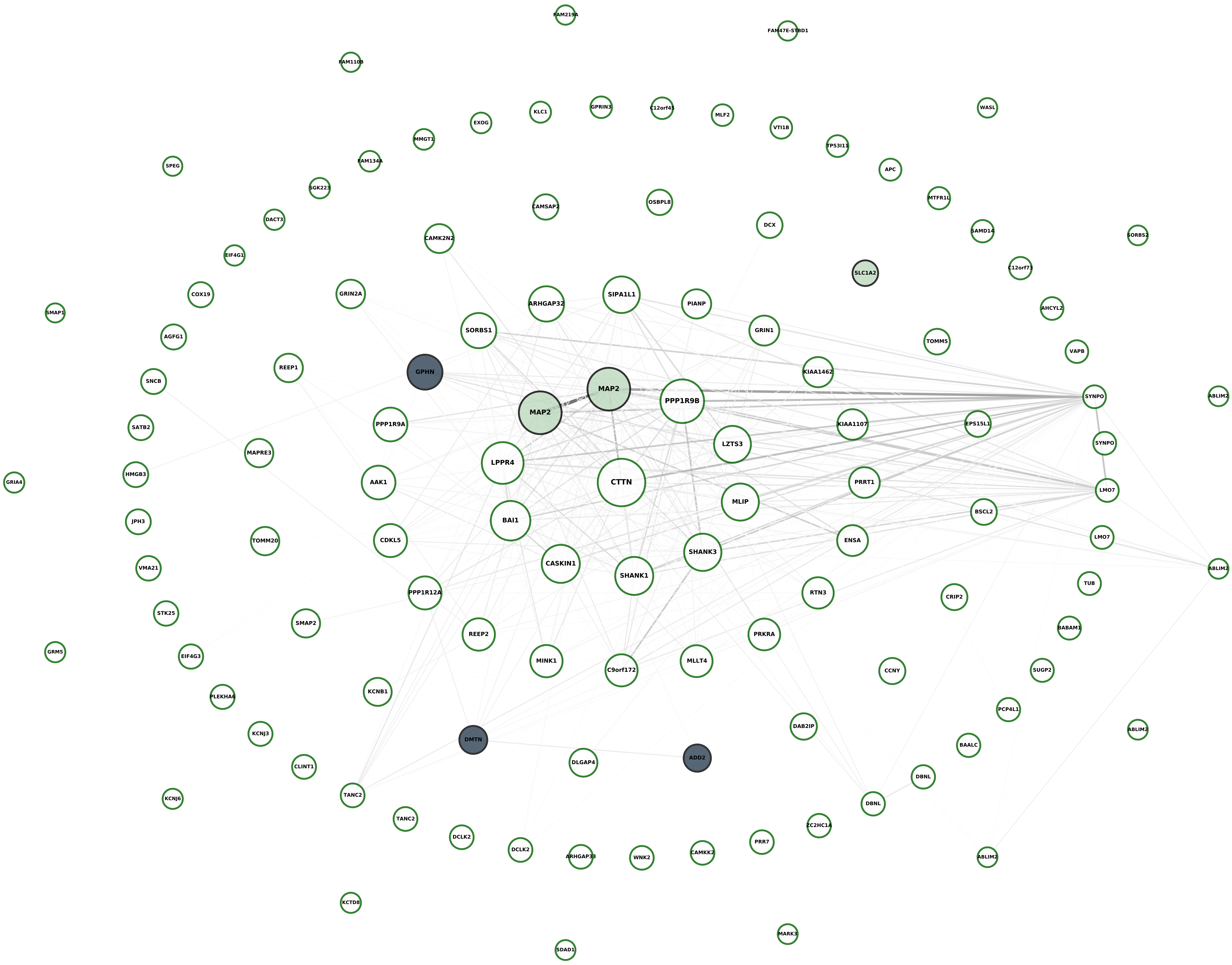

**Module 23, n = 111,
FDR Corrected pvalue 1**

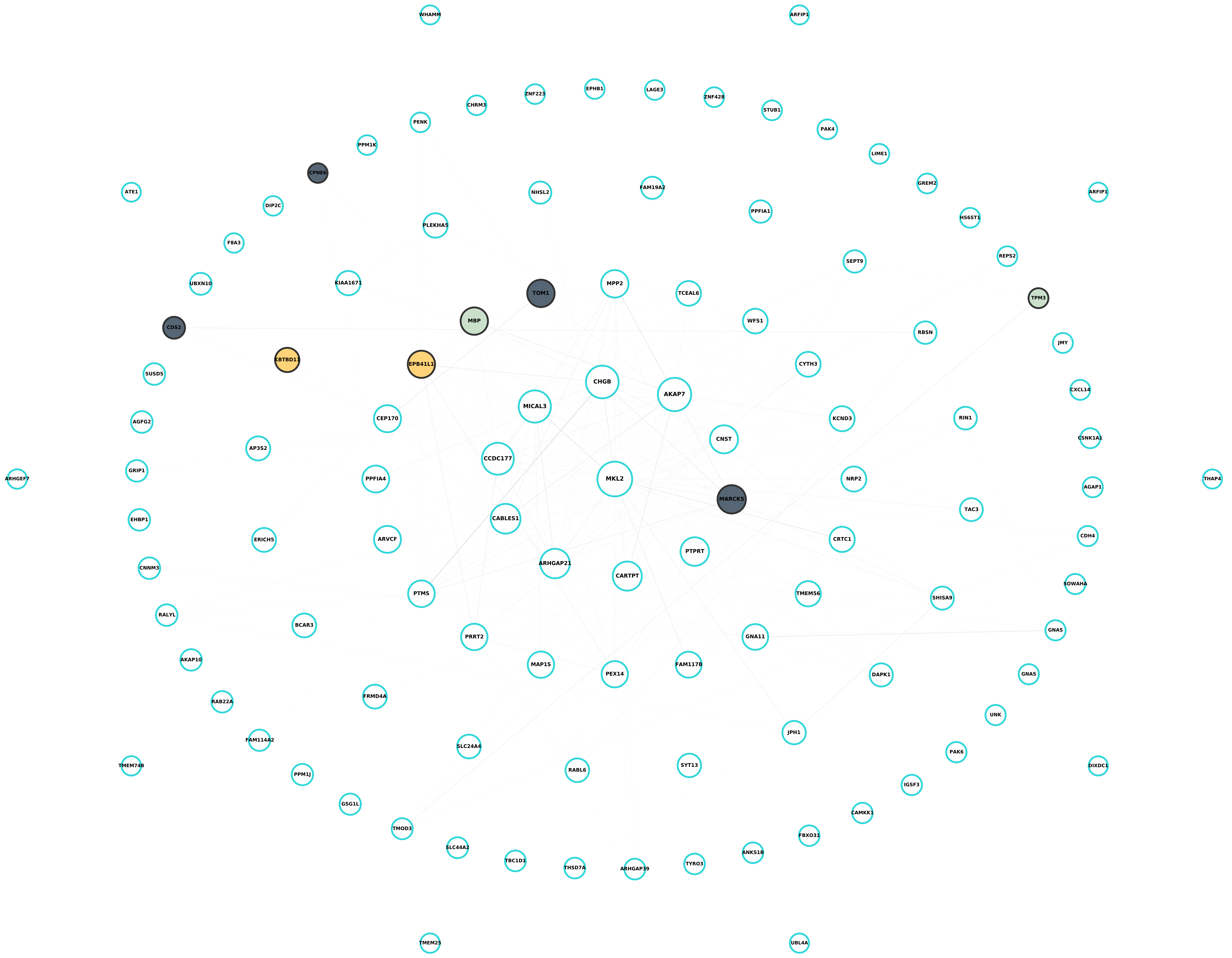

**Module 24, n = 111,**
**FDR Corrected pvalue 1**

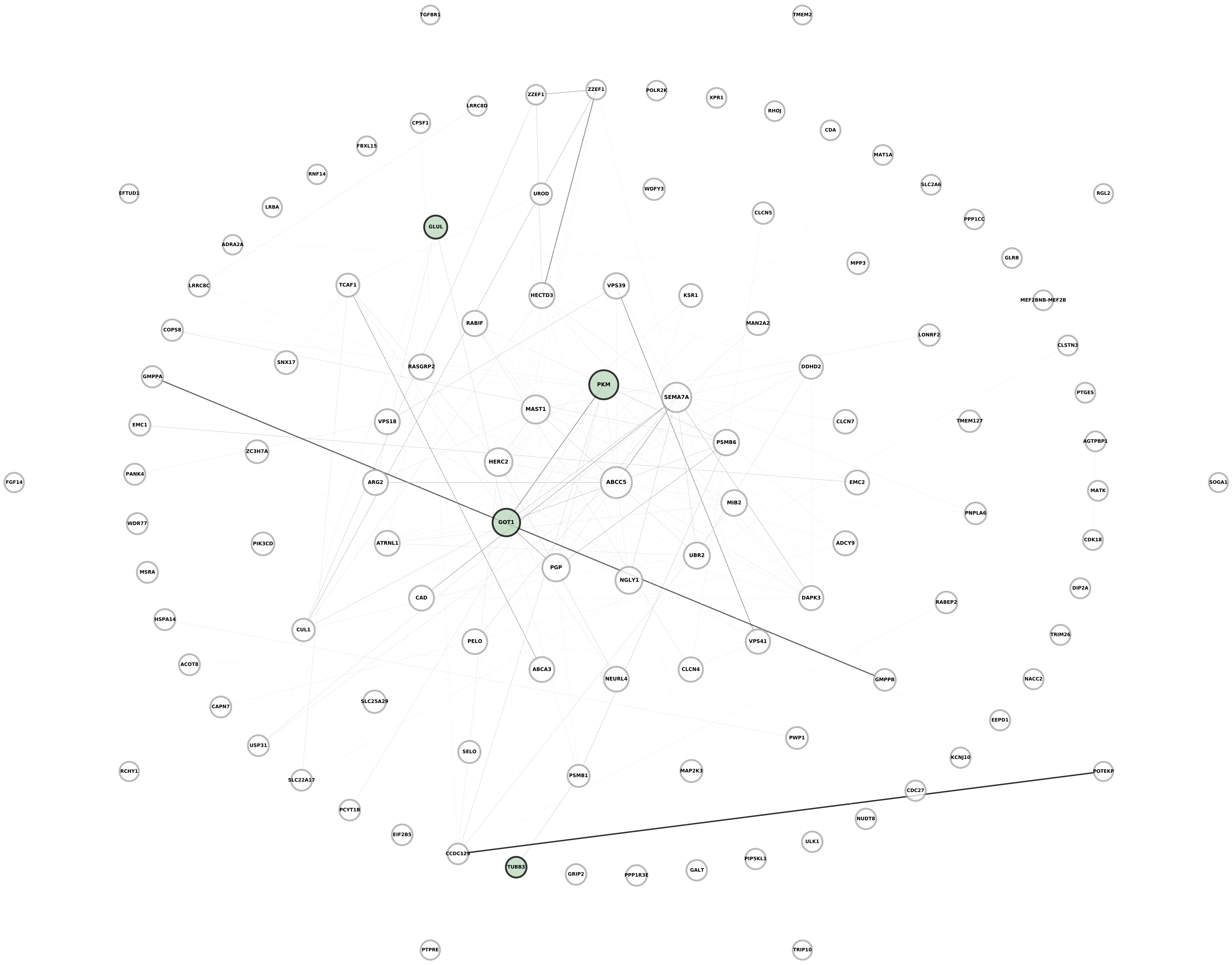

**Module 25, n = 104,**
**FDR Corrected pvalue 0.012325**

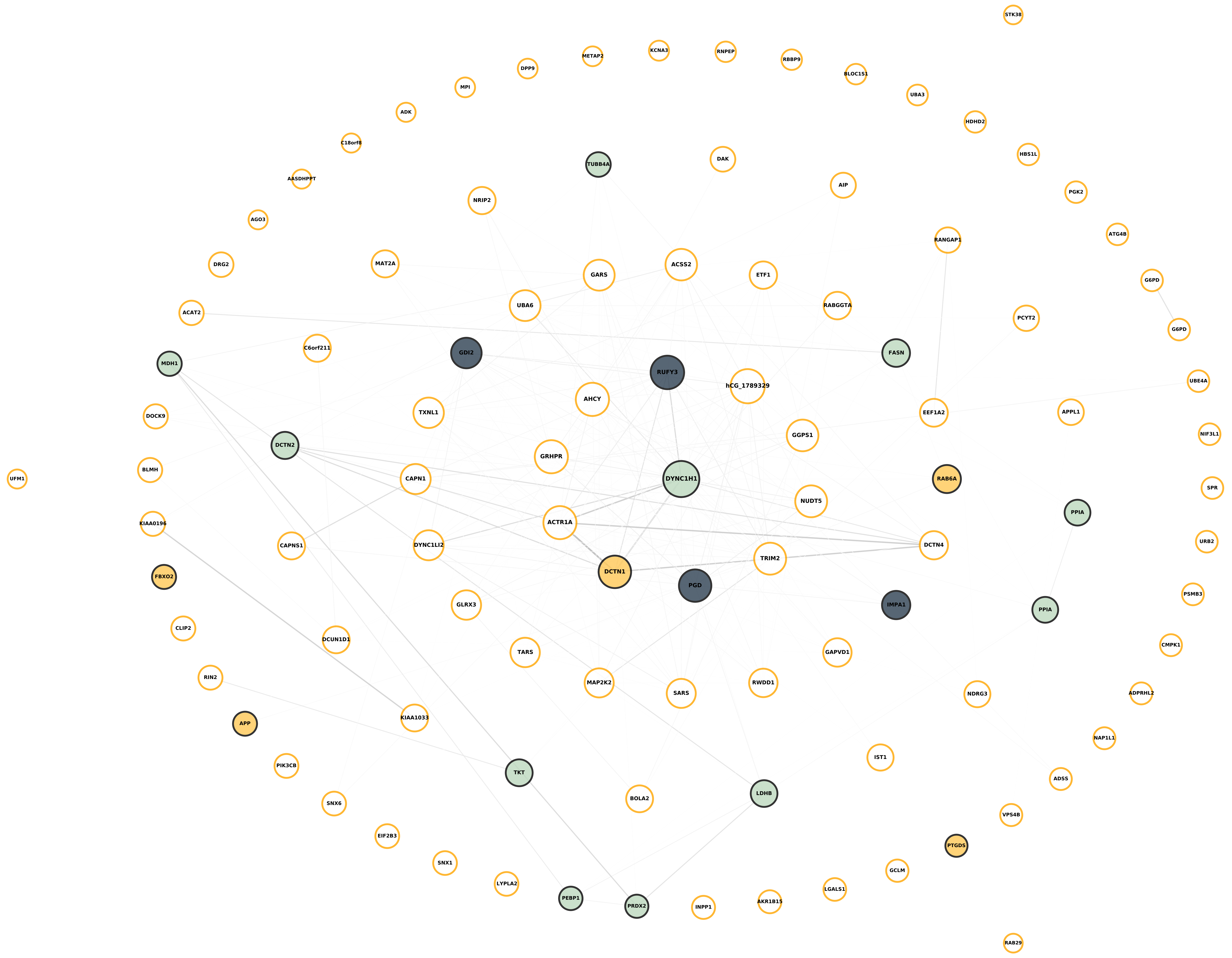

Module 26, n = 97,
FDR Corrected pvalue 1

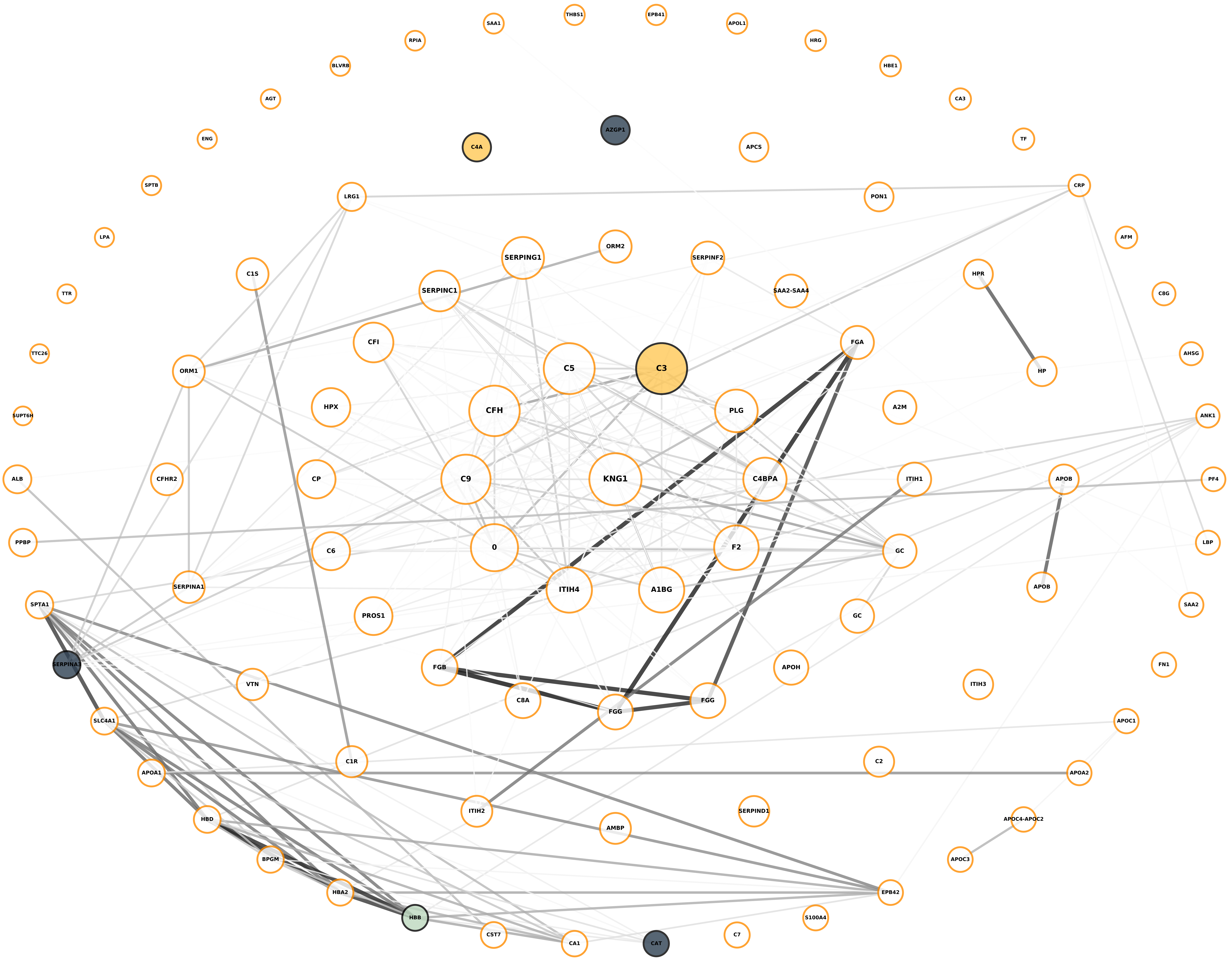

**Module 27, n = 93,**
**FDR Corrected pvalue 1**

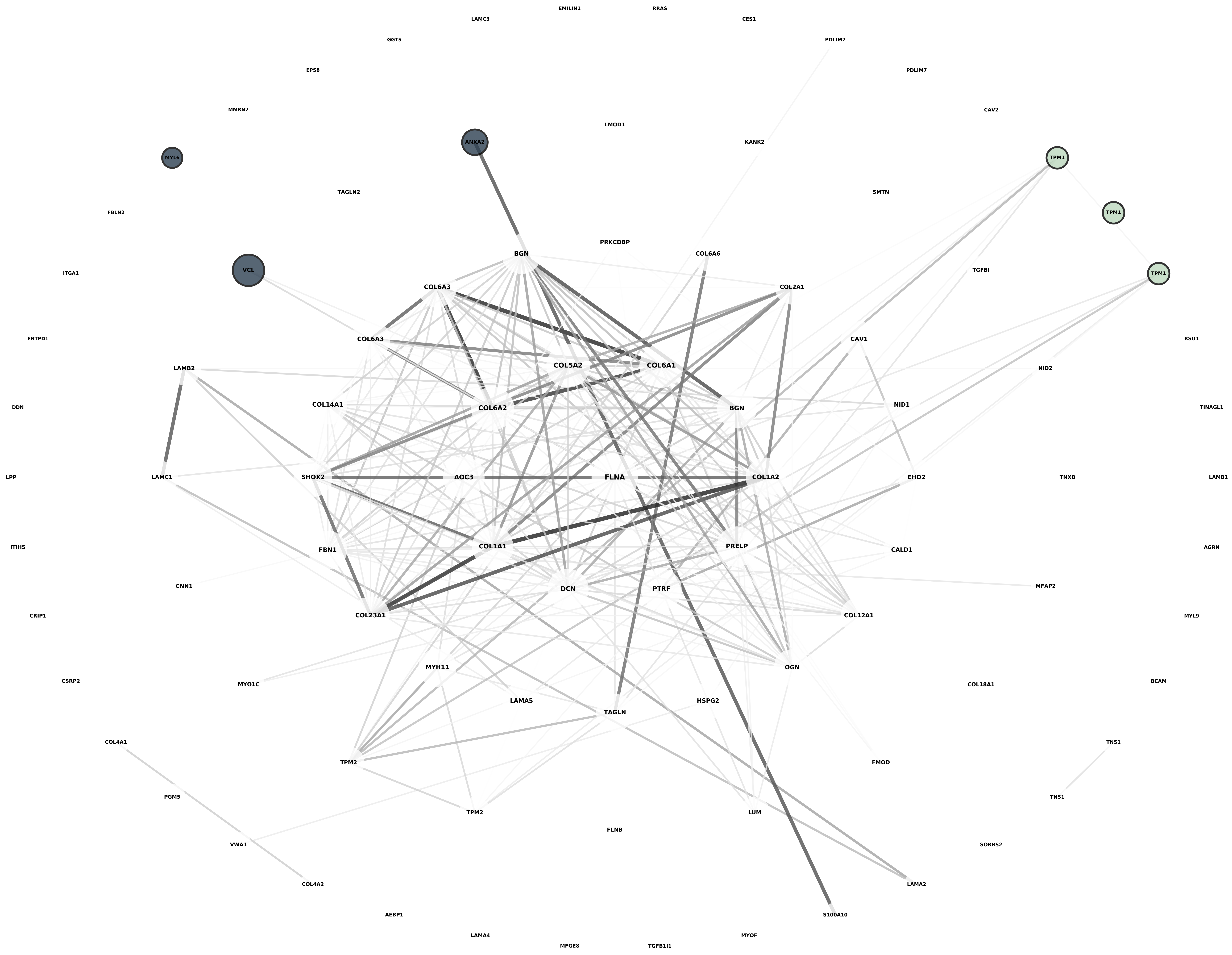

**Module 28, n = 86,**
**FDR Corrected pvalue 3.9464e-06**

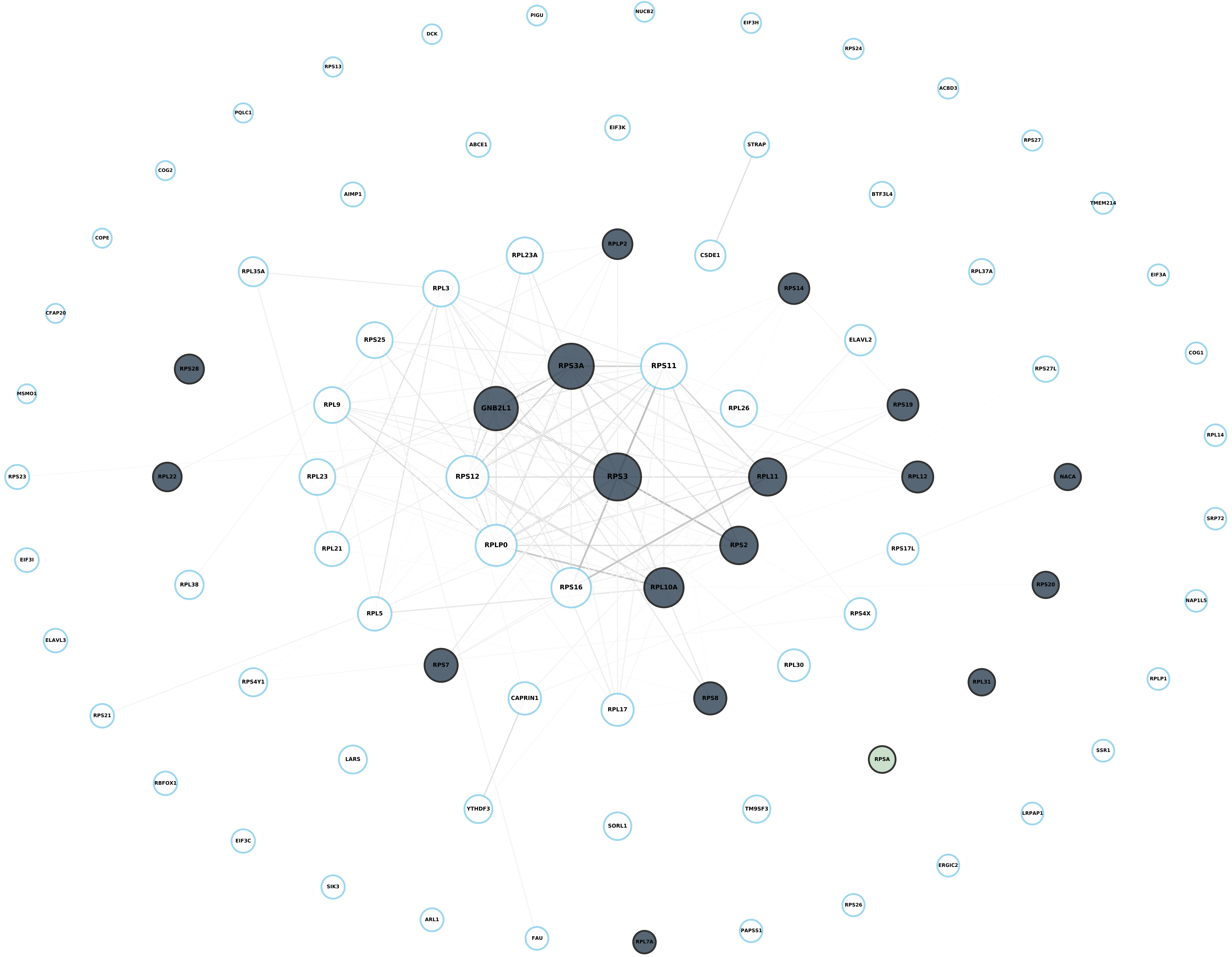

**Module 29, n = 81,**
**FDR Corrected pvalue 0.010991**

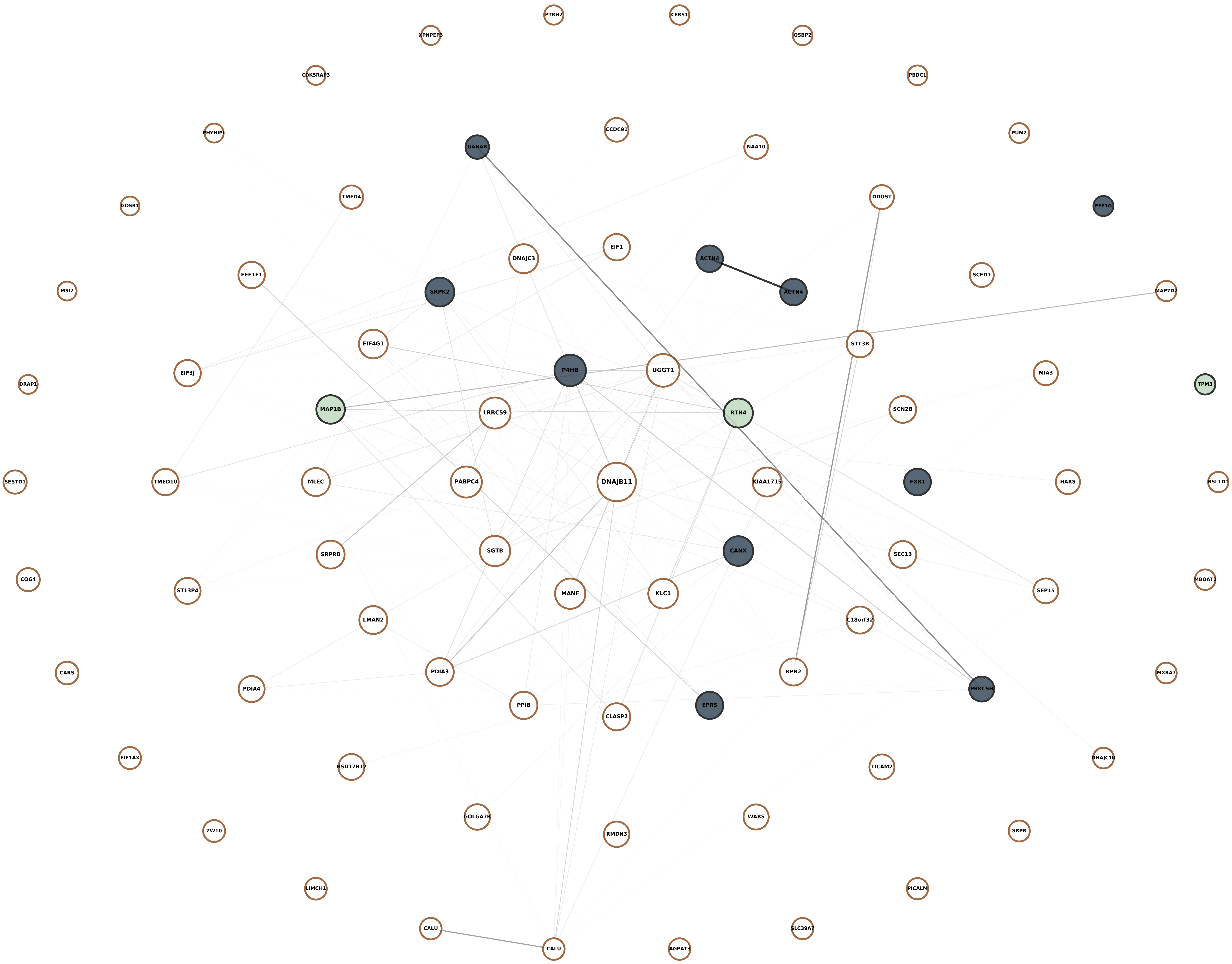

**Module 30, n = 73,**
**FDR Corrected pvalue 0.0016322**

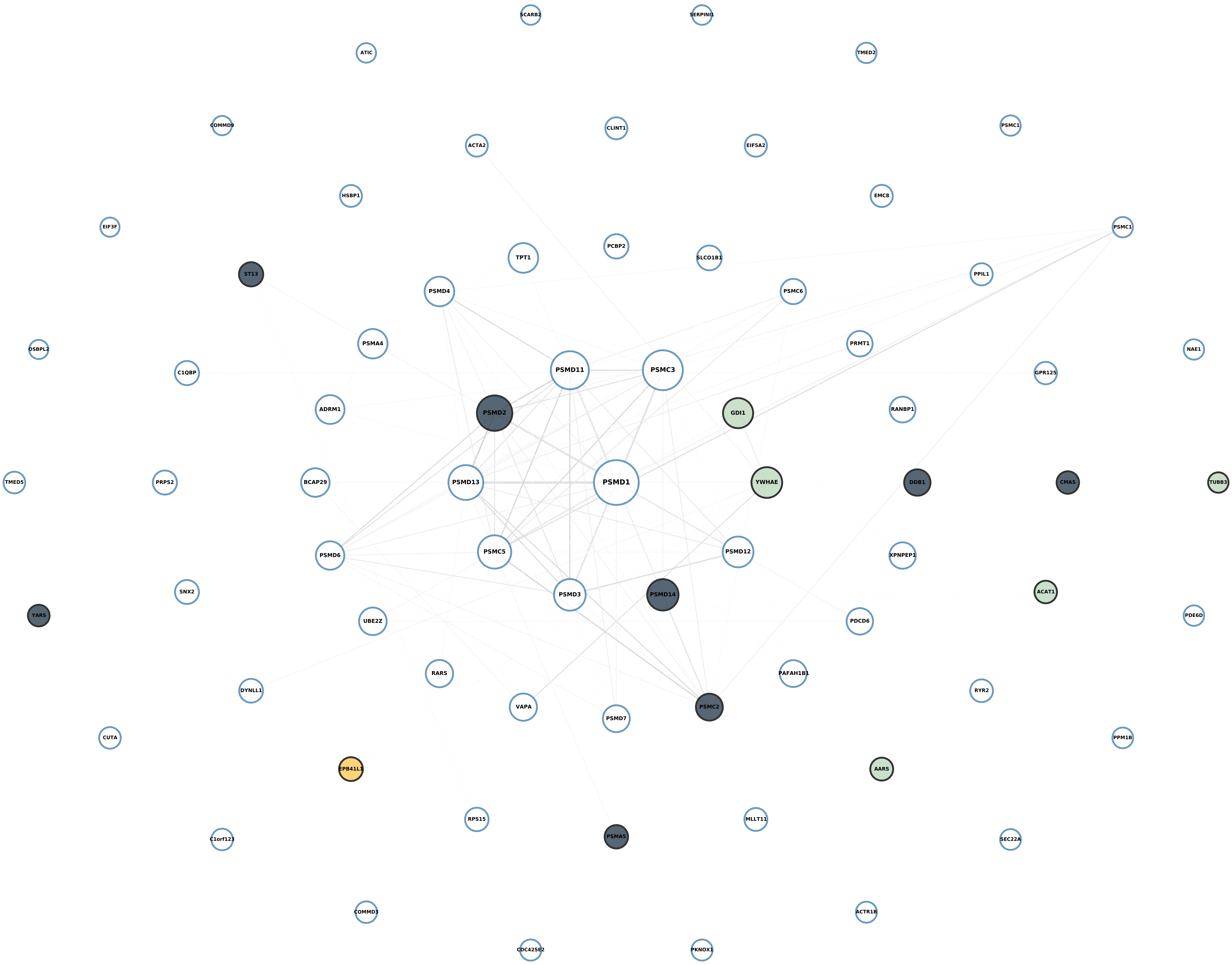

**Module 31, n = 72,**
**FDR Corrected pvalue 0.99401**

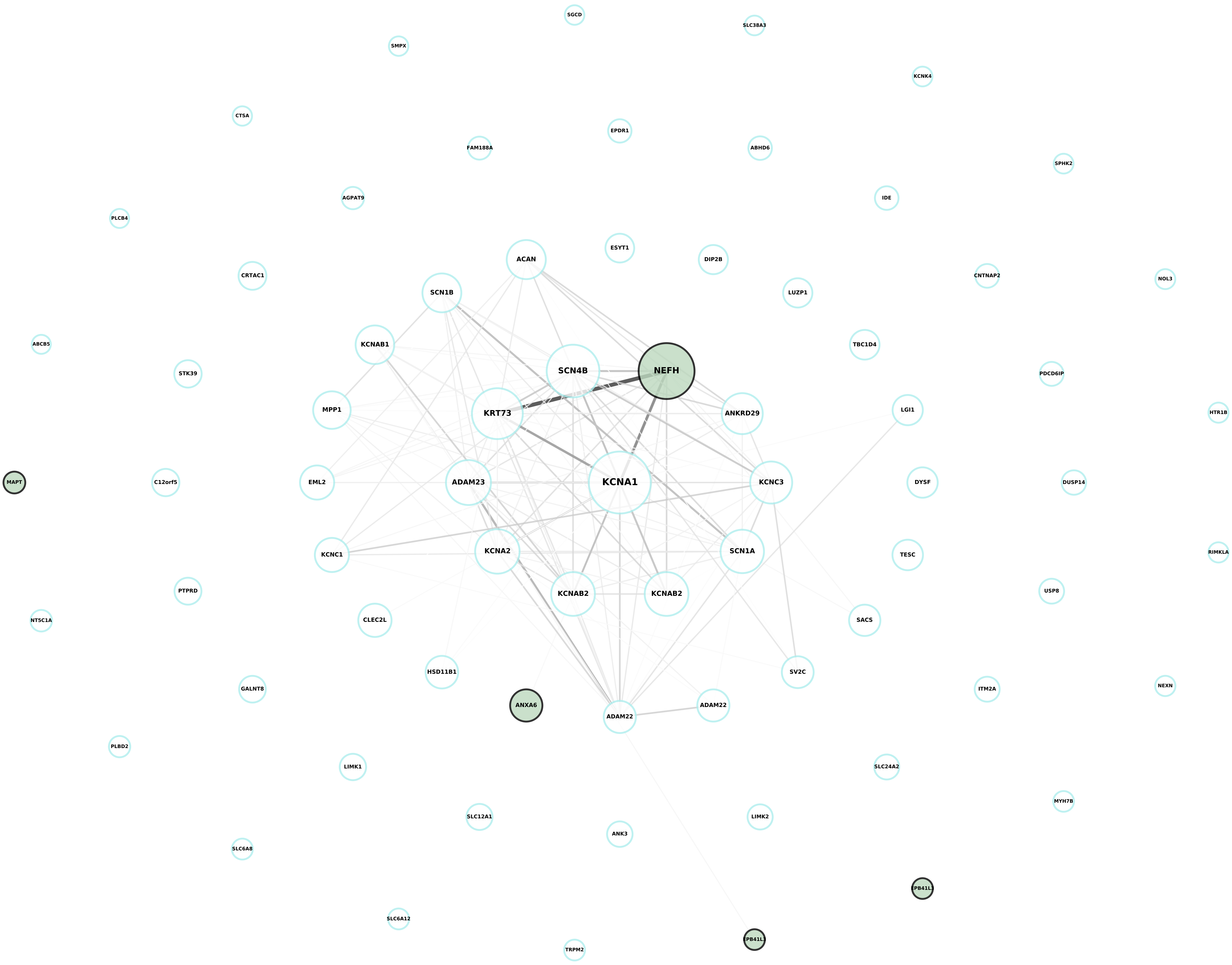

Module 32, n = 72,
FDR Corrected pvalue 1

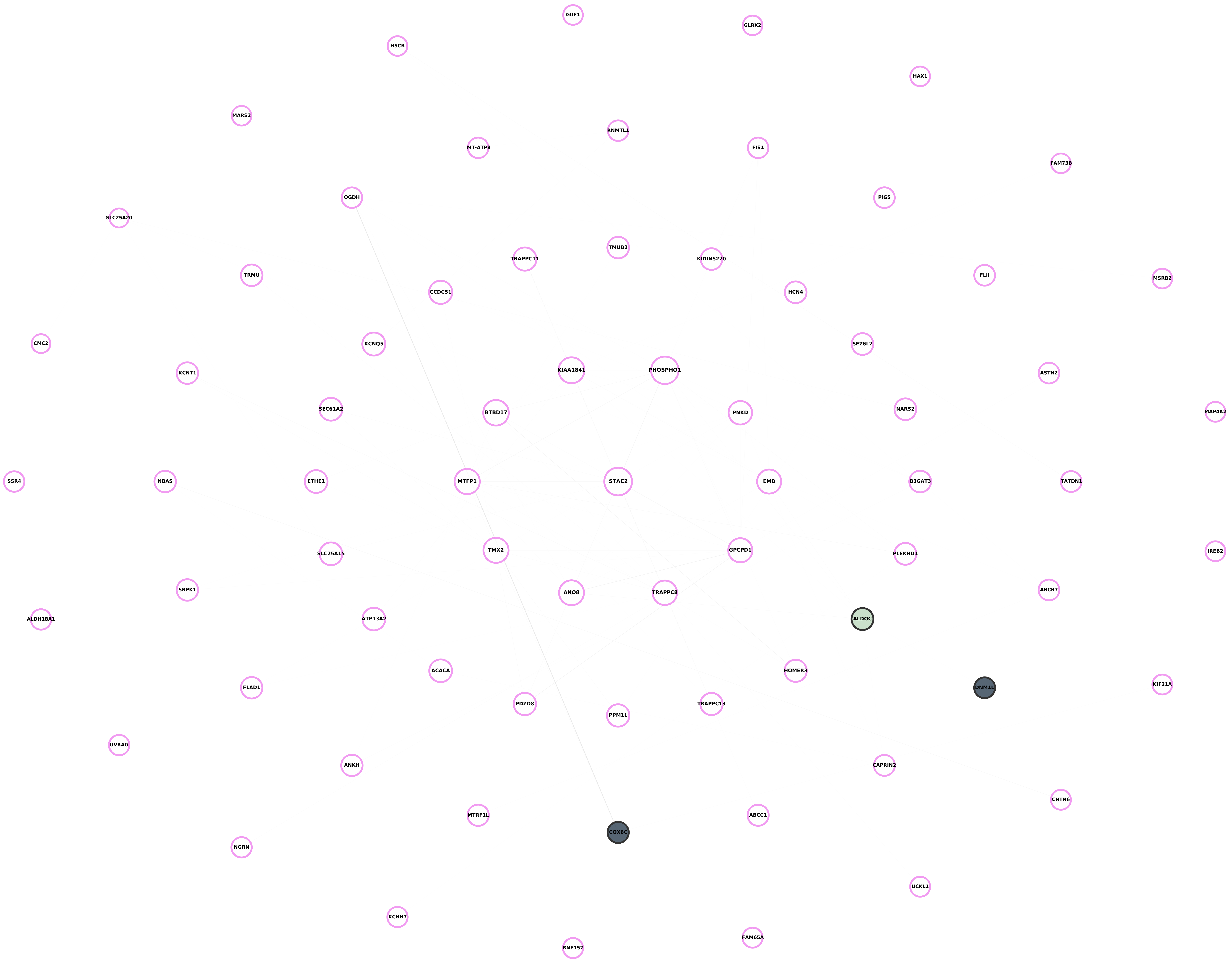

**Module 33, n = 65,**
**FDR Corrected pvalue 1**

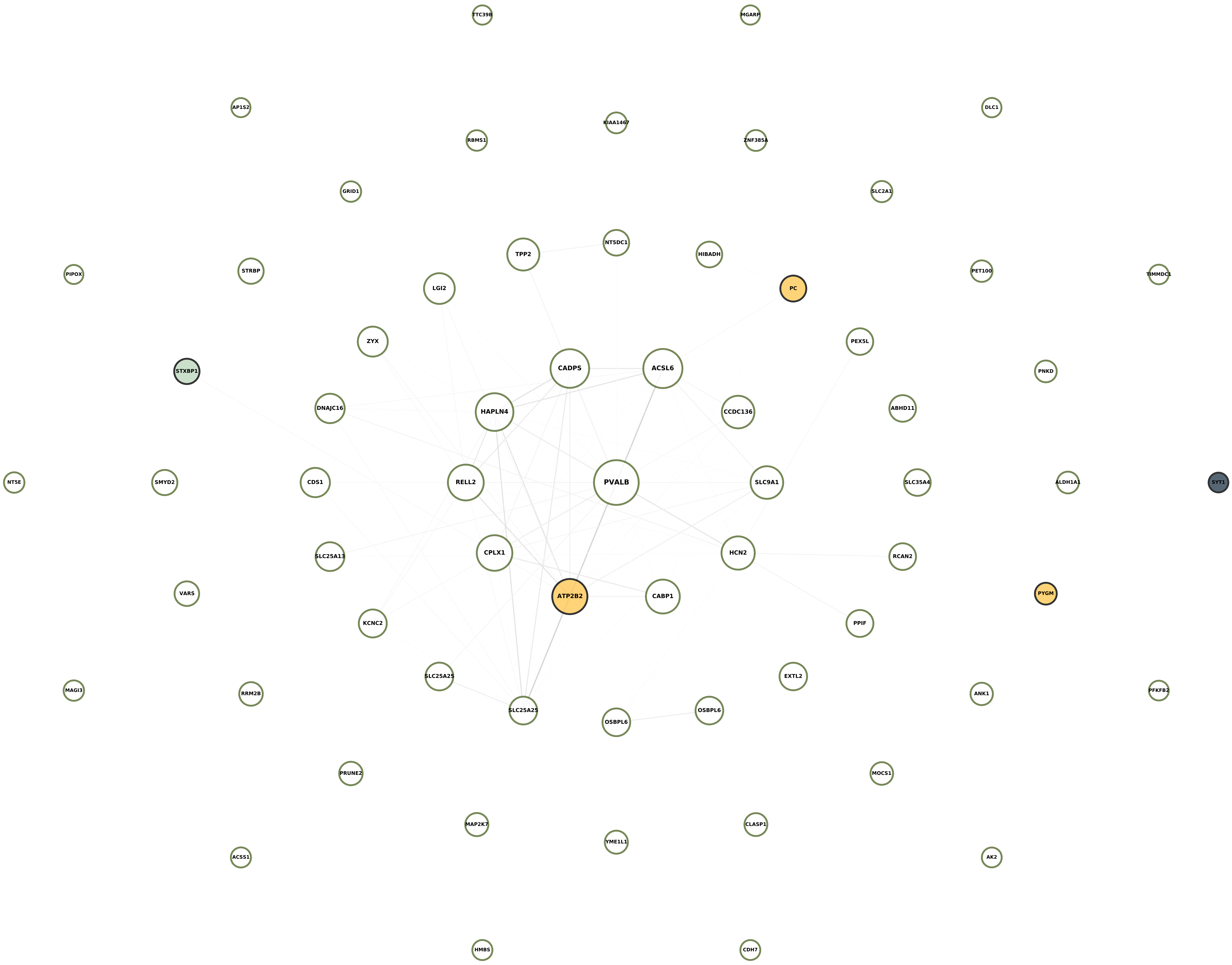

### Module 34, n = 61,
### FDR Corrected pvalue 0.86525

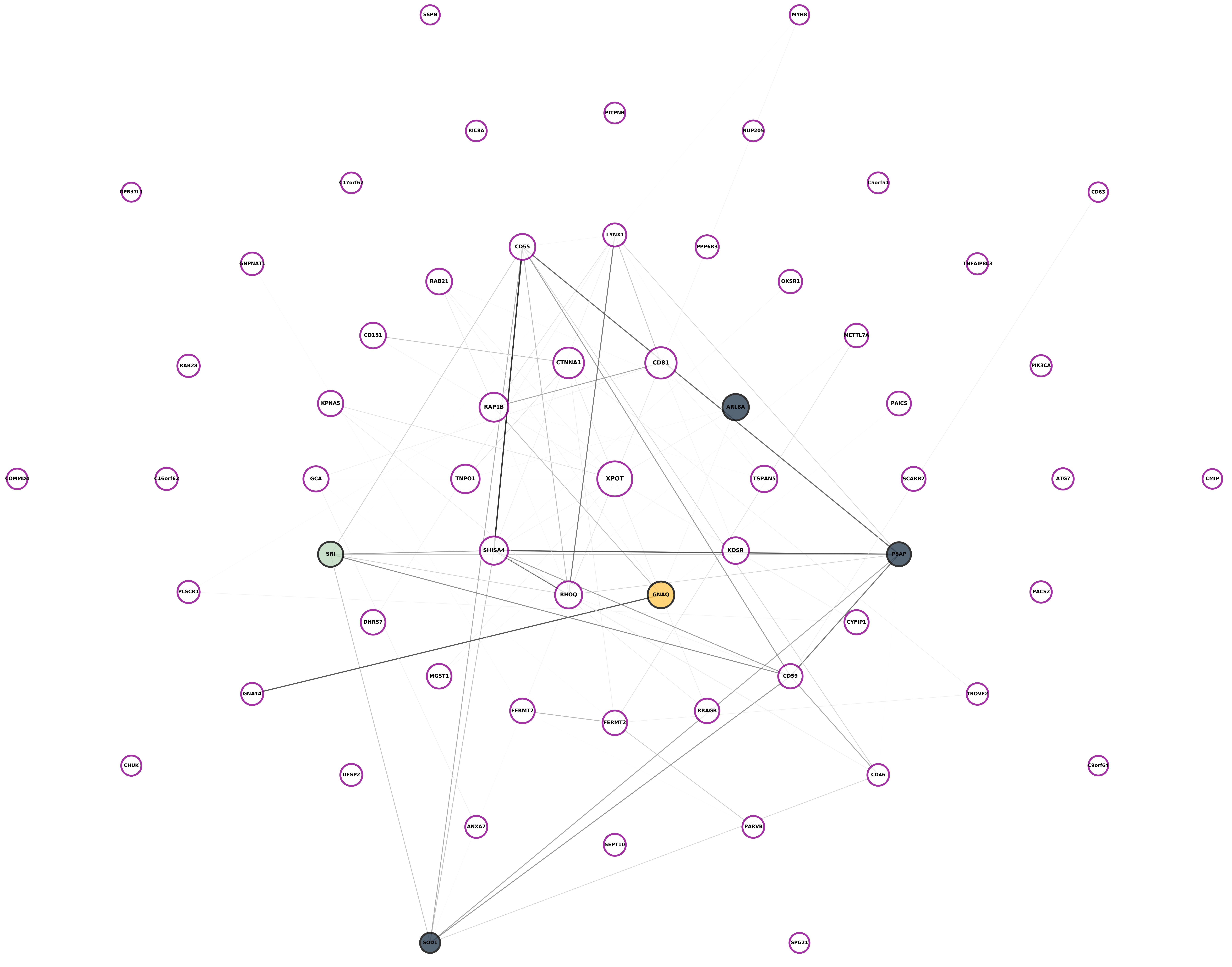

**Module 35, n = 60,**
**FDR Corrected pvalue 1**

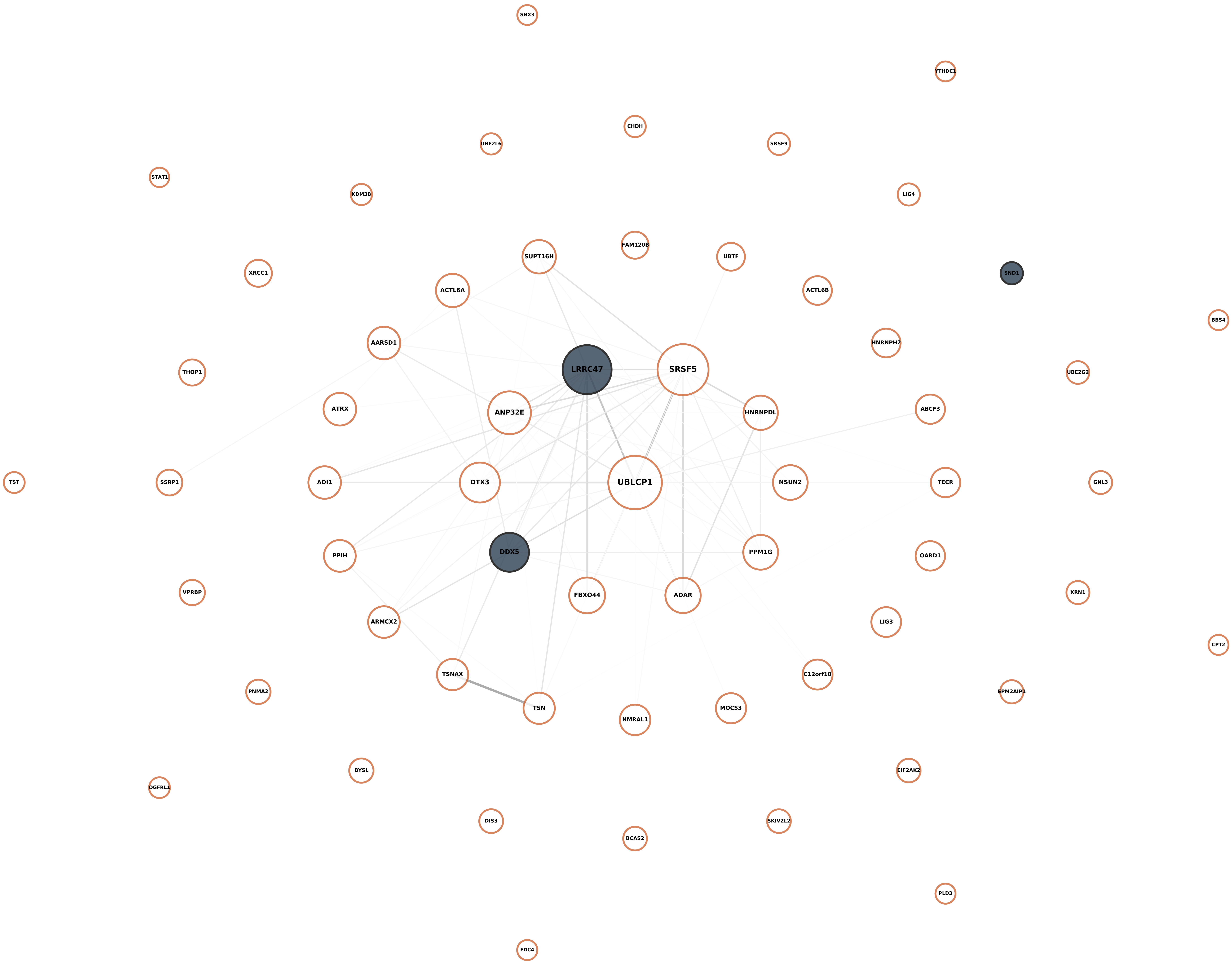

Module 36, n = 60,
FDR Corrected pvalue 0.12774

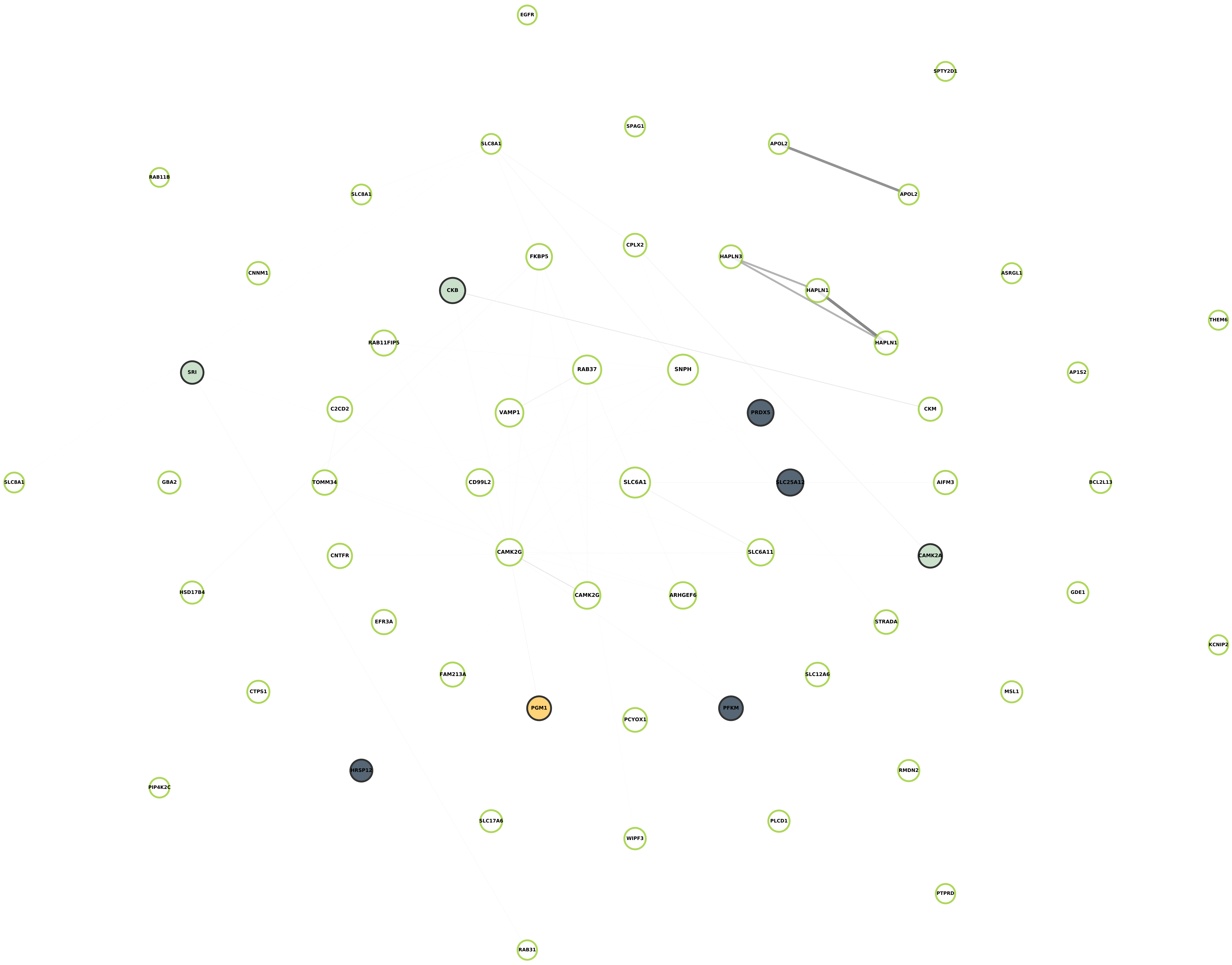

**Module 37, n = 56,**
**FDR Corrected pvalue 0.50737**

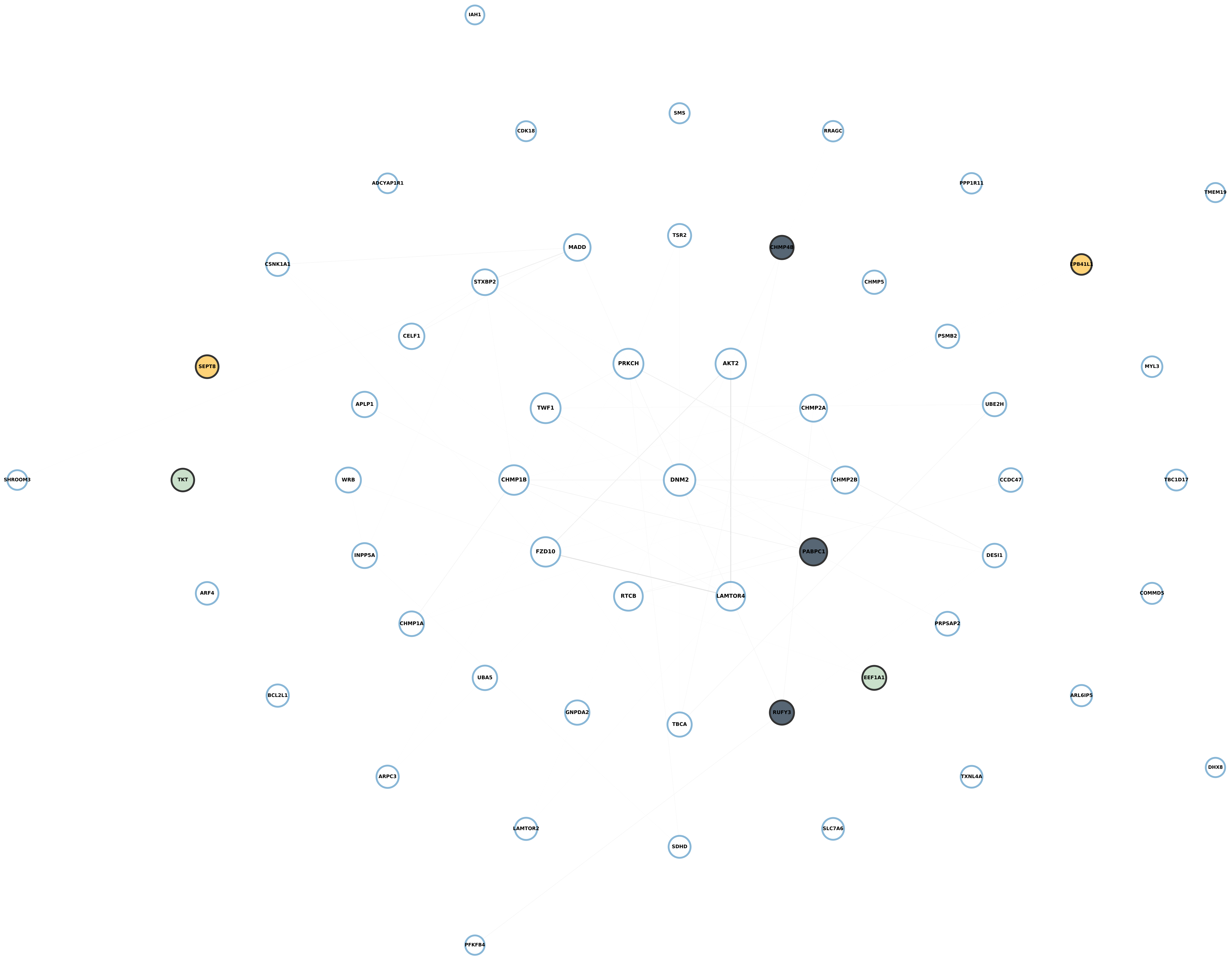

**Module 38, n = 53,**
**FDR Corrected pvalue 0.00025693**

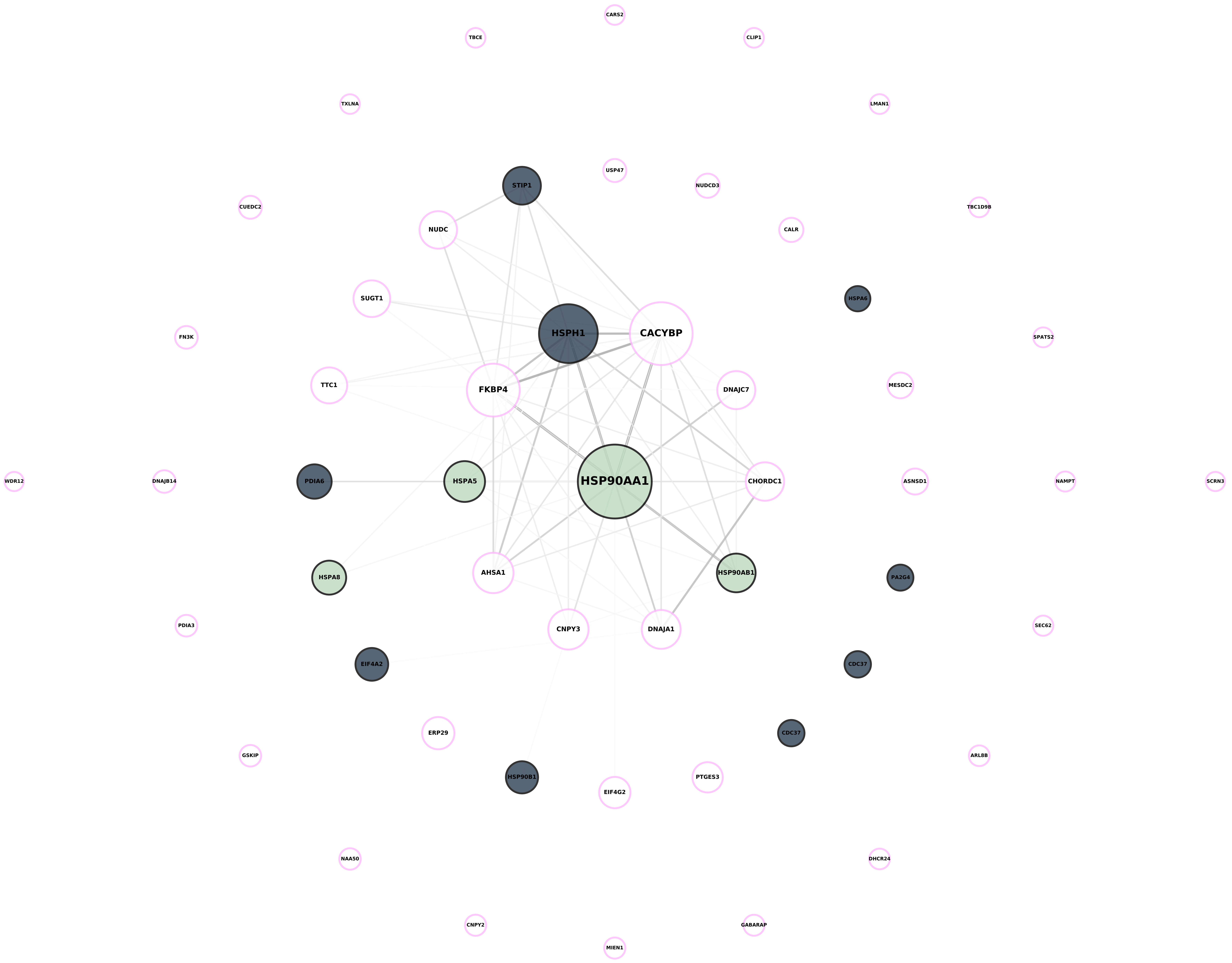

**Module 39, n = 48,**
**FDR Corrected pvalue 0.96802**

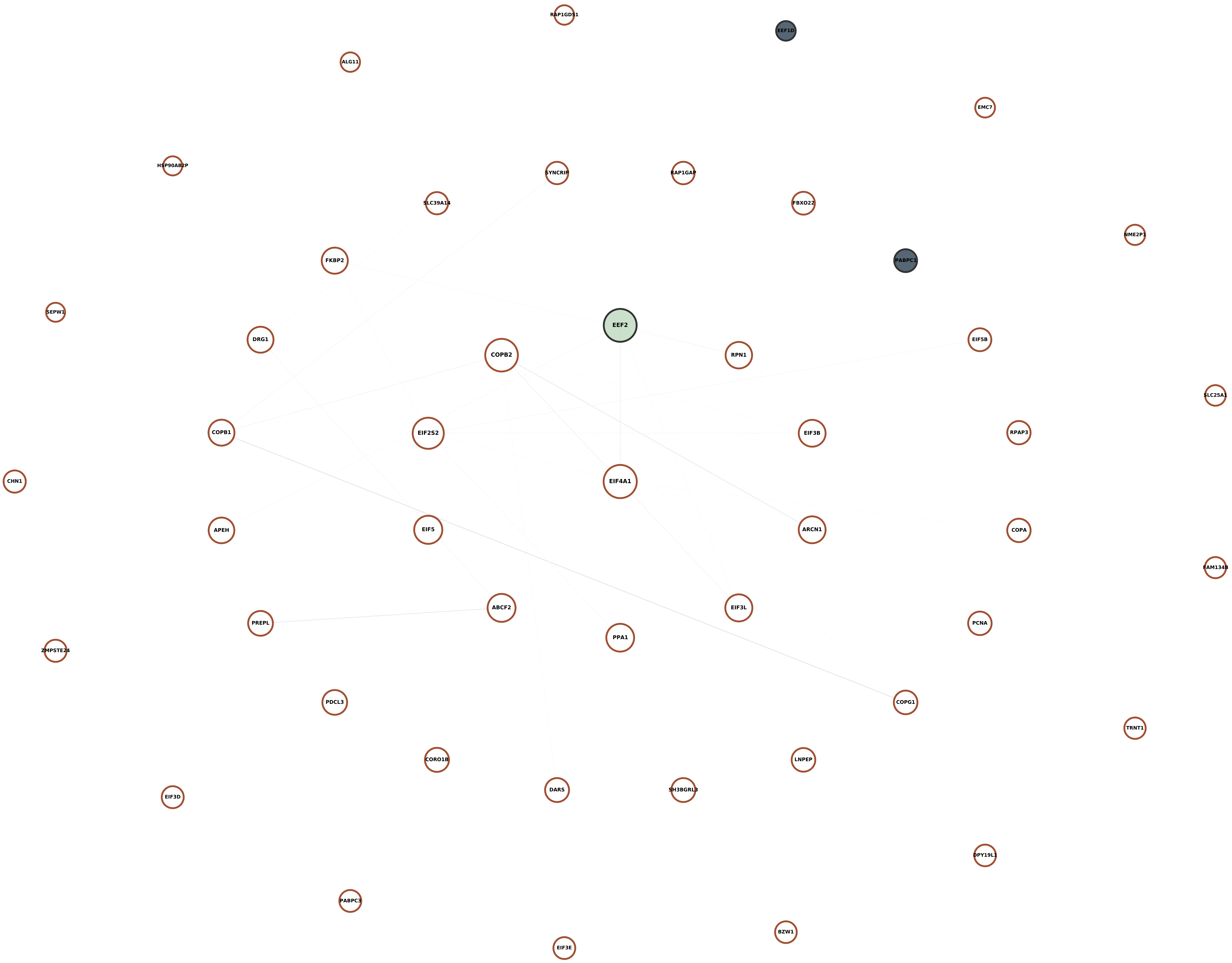

**Module 40, n = 41,**
**FDR Corrected pvalue 0.49975**

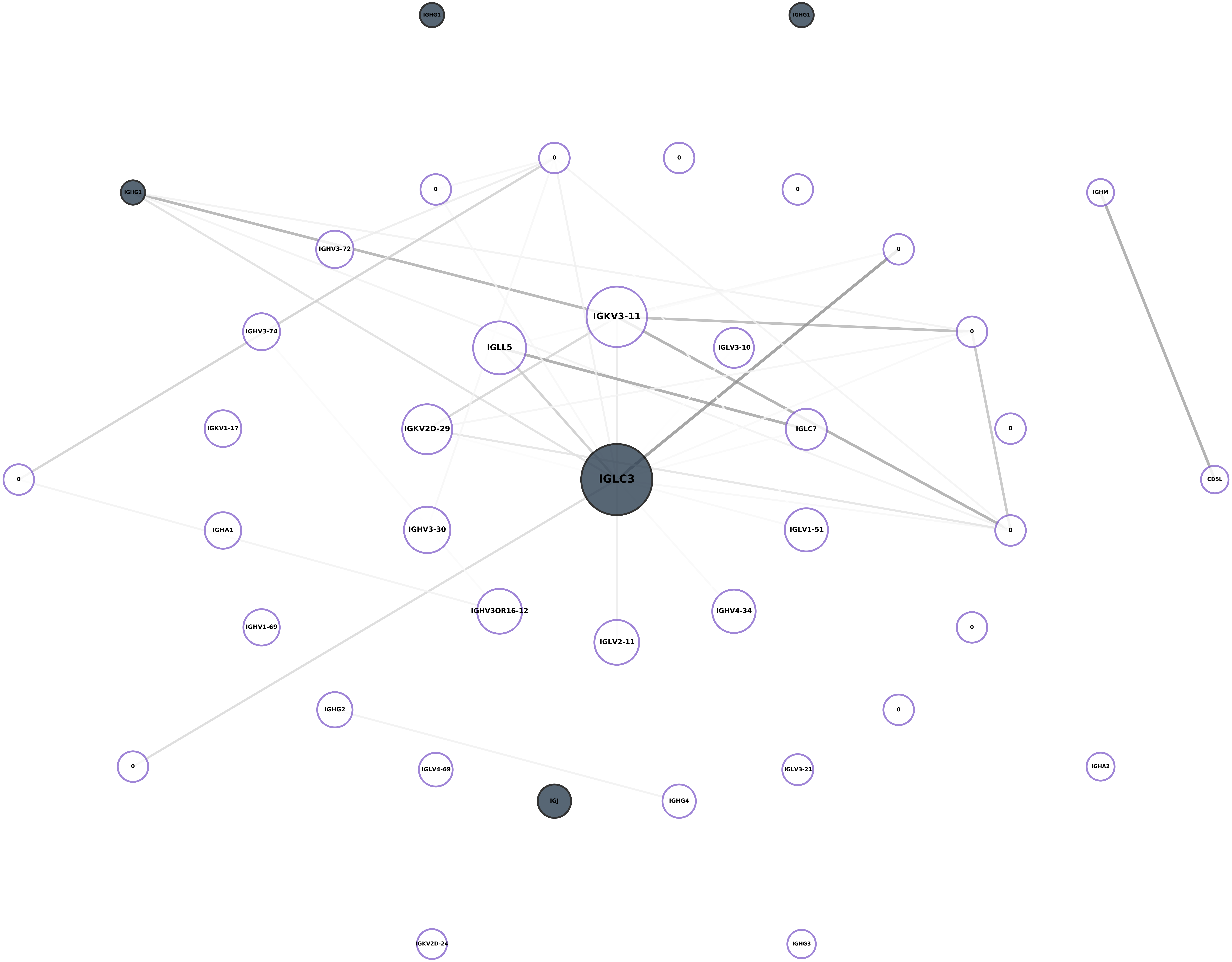

**Module 41, n = 37,**
**FDR Corrected pvalue 0.0064004**

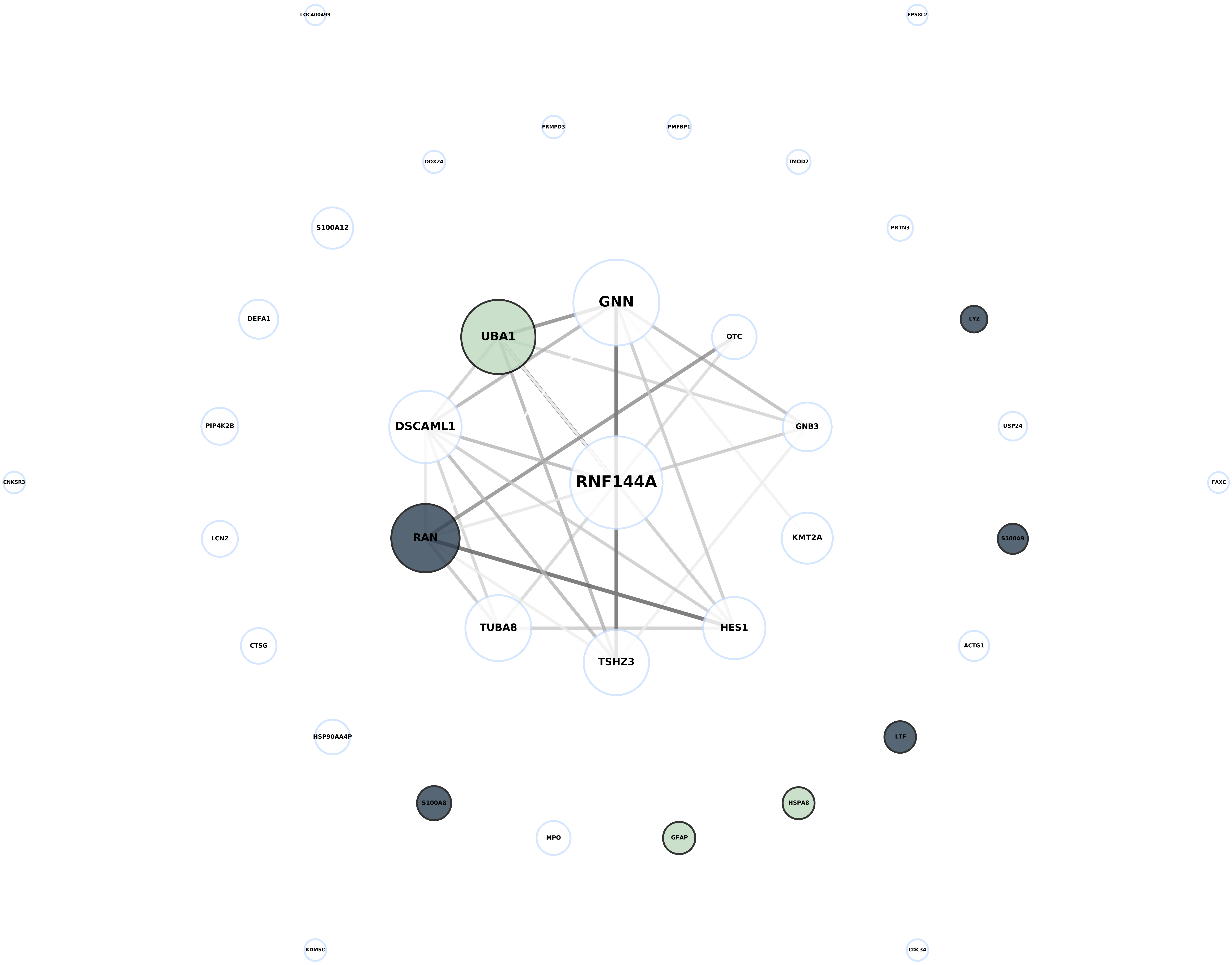

**Module 42, n = 32,
FDR Corrected pvalue 1**

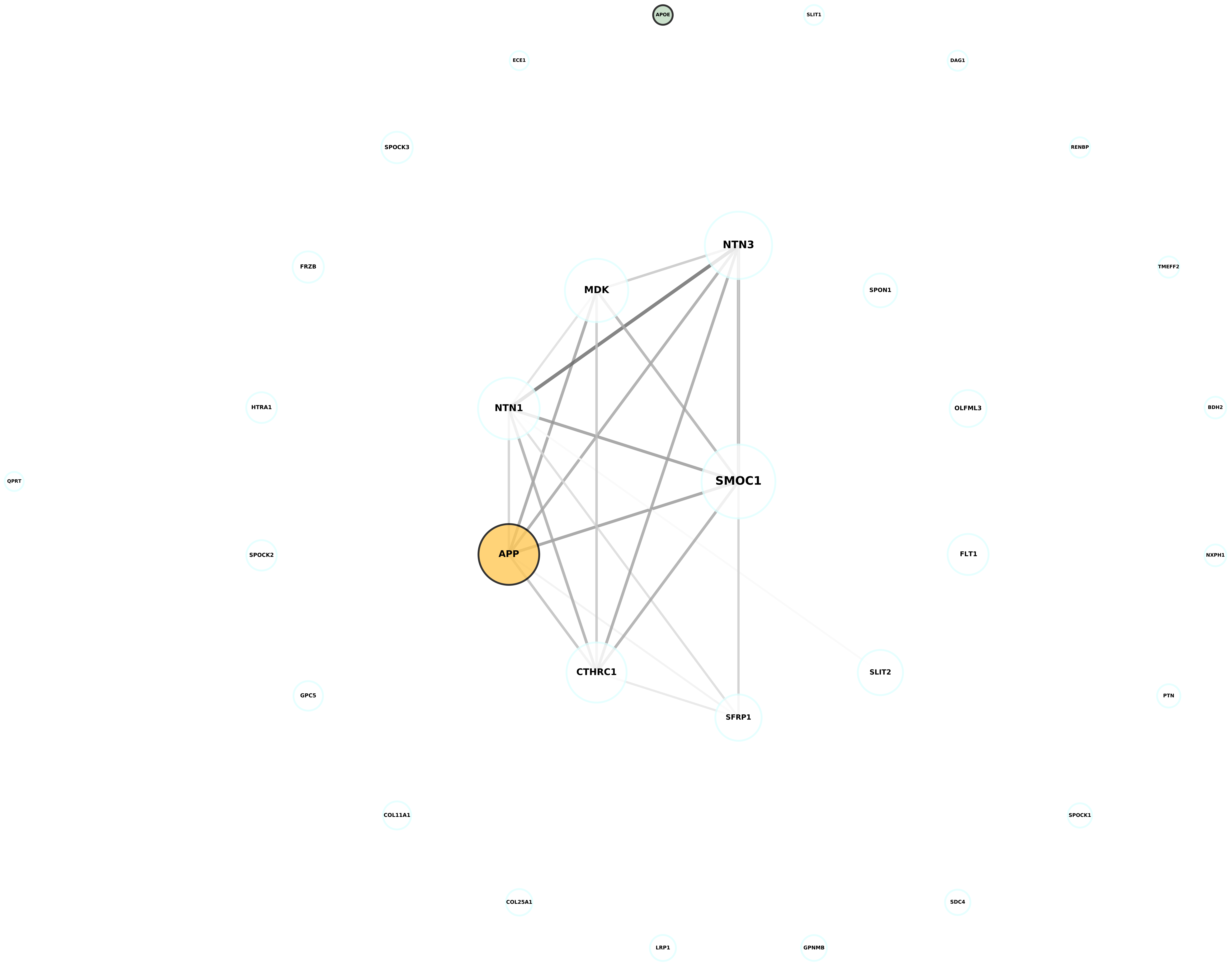

**Module 43, n = 30,**
**FDR Corrected pvalue 0.54857**

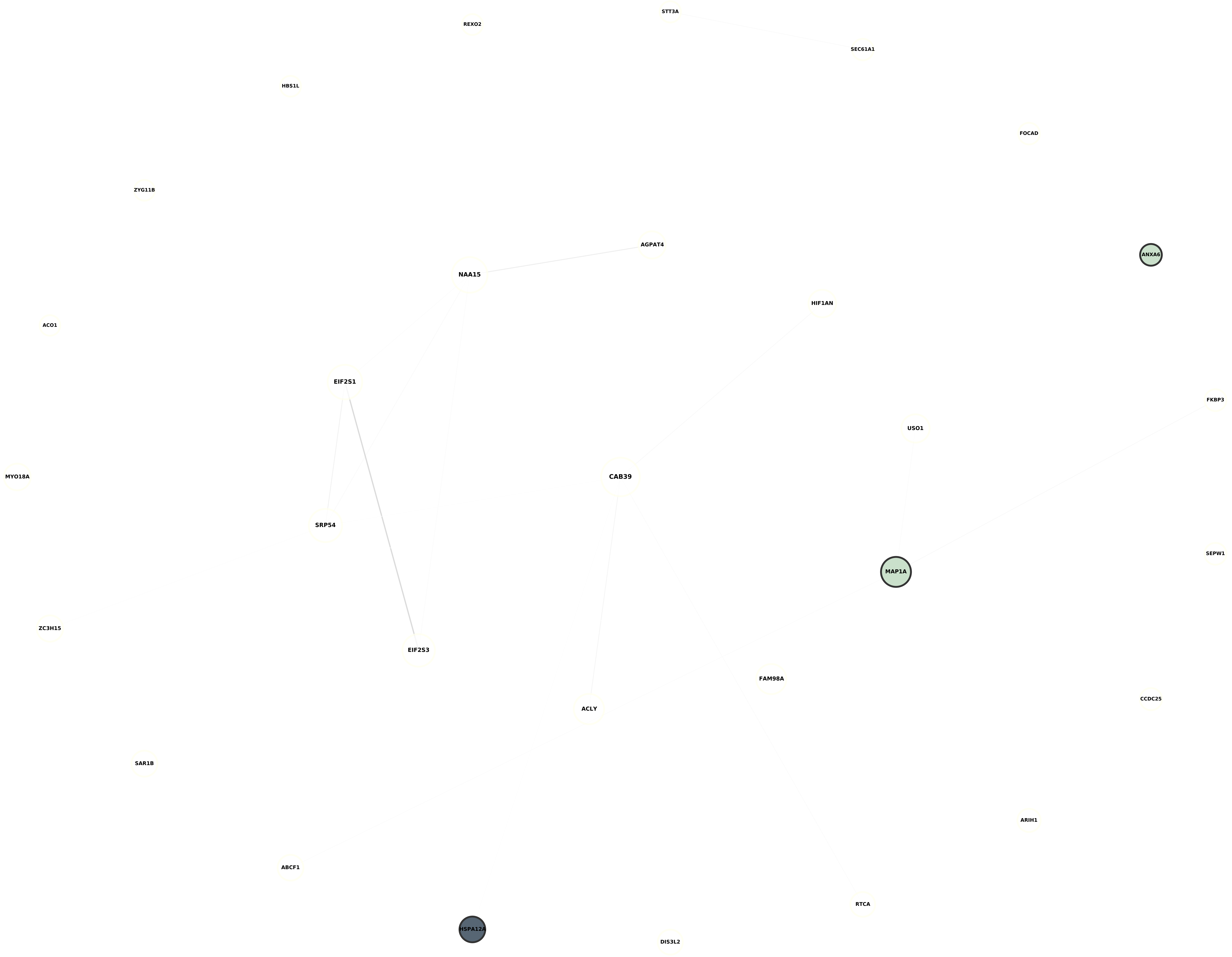

**Module 44, n = 28,**
**FDR Corrected pvalue 2.3129e-06**

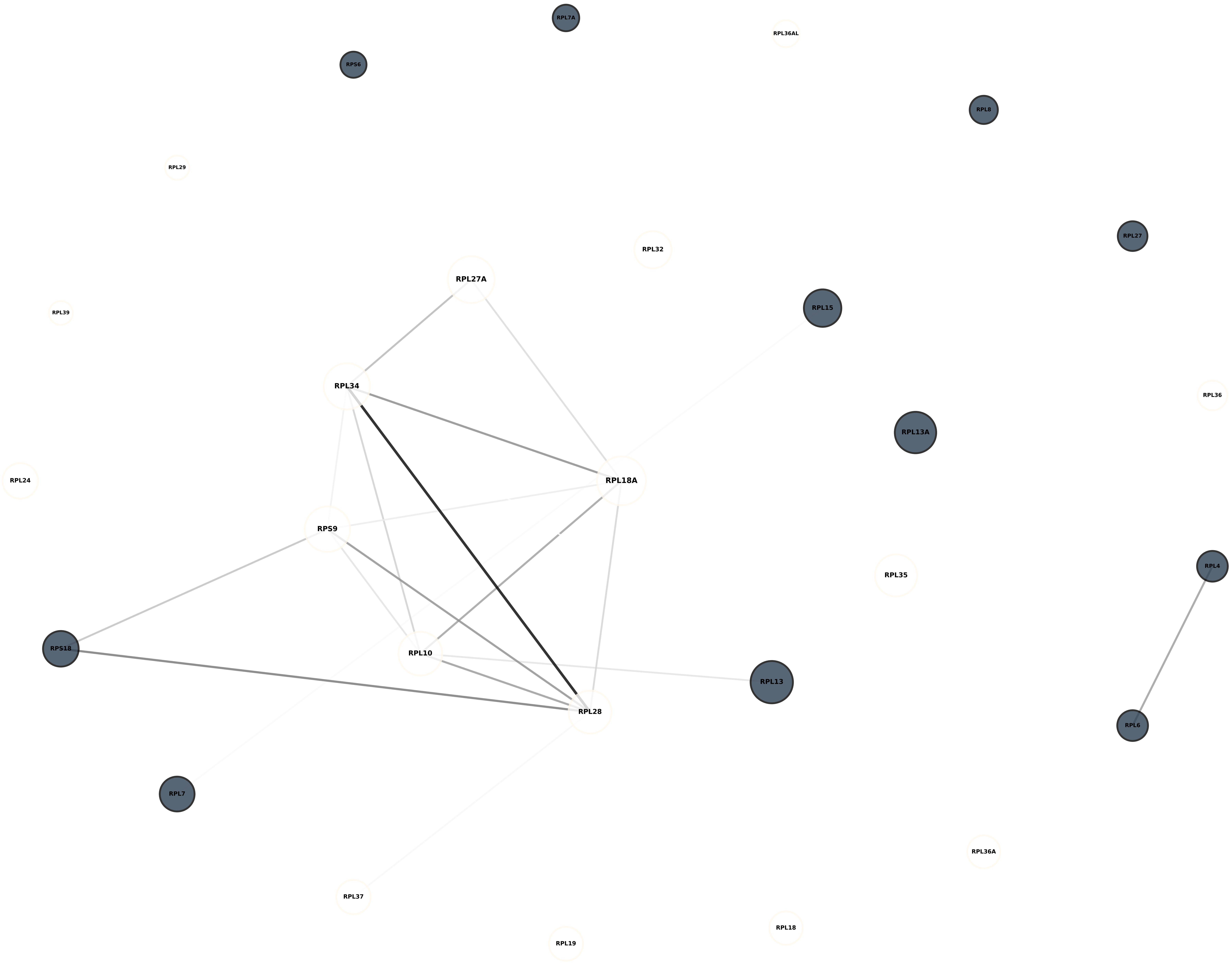
